# A near-chromosome level genome assembly of *Anopheles stephensi*

**DOI:** 10.1101/2020.04.27.063040

**Authors:** Afiya Razia Chida, Samathmika Ravi, Suvratha Jayaprasad, Kiran Paul, Jaysmita Saha, Chinjusha Suresh, Saurabh Whadgar, Naveen Kumar, Raksha Rao K, Chaitali Ghosh, Bibha Choudhary, Suresh Subramani, Subhashini Srinivasan

## Abstract

Malaria remains a major healthcare risk to growing economies like India and a chromosome-level reference genome of *Anopheles stephensi* is critical for successful vector management and an understanding of vector evolution. We report a chromosome-level assembly of an Indian strain from draft genomes of two strains using a homology-based iterative approach. The resulting assembly with an L50 of 9 had long enough scaffolds for building 90% of the three chromosomes using physical markers. The sequencing of individuals reveals a genetic diversity ten times higher than what is reported. Based on the developmental transcriptome and orthology of the 54 olfactory receptors (ORs) to those of other *Anopheles* species, we identify olfactory receptors with the potential for host recognition in the genus *Anopheles*. A comparative analysis of the genomes suggests limited inter-chromosomal gene flow and chromosomal arm switching with a potential role in adaptive radiation within the genus *Anopheles*.

## Introduction

There are ~450 recorded species in the genus *Anopheles* (*An*.) with roughly 100 being vectors of malaria in various endemic regions in India, Africa and elsewhere. Among the malaria vectors, only 30-40 transmit parasites of the genus *Plasmodium*. Efforts to obtain reference genome sequences for various vectors that spread malaria by transmitting *Plasmodium* are under way. Chromosome level assemblies of *An. gambiae* and *An. funestus*, both malaria vectors from Africa, have been reported^1,2^. Draft genomes of an *An. stephensi* strain from India^3^ and sixteen other *Anopheles* genera^4^ have also been reported.

*An. stephensi* is an urban species of mosquito that is responsible for causing roughly 12% malaria in India. The major genome resource so far available for *An. stephensi* is the draft genome reported in 2014^3^. In this report, a number of sequencing technologies including Pacbio, 454, Illumina and BAC-end sequencing were used to generate the draft assembly. The 454 reads were sequenced from multiple mate-pair libraries and include 12.2X coverage from single-end reads, 2.2X coverage from 3 kilobase (kb) paired-end reads, 3.4X coverage from 8 kb paired-end reads, and 1.7X coverage from 20 kb paired-end reads. The majority of 454 reads was in the range of 194 to 395 base-pairs (bp) in length. Illumina short reads with a coverage of 86.4X, read length of 101 bp paired end reads and an average insert size of approximately 200 bp were sequenced. Also, ten cells of PacBio reads using RS1 sequencing for male genomic DNA produced 5.2X coverage with a median read length of 1,295 bp. A hybrid assembly combining 454 and Illumina data was further improved by filling the gaps with error corrected PacBio reads and scaffolding using BAC-ends with an insert size of ~140 kb. The resulting assembly contained 23,371 scaffolds spanning 221 Mb, including 11.8 Mb (5.3%) of gaps filled with Ns (unspecified nucleotides). The L50 of the assembly was 37. The N50 scaffold size was 1.59 Mb and the longest scaffold was 5.9 Mb. Considering that the assembly included various sequencing technologies and strategies, the assembly has the least bias from the use of particular technology platforms.

More recently, a draft genome assembly of sixteen diverse species of *Anopheles* mosquitoes was generated, including a strain of *An. stephensi* from Pakistan (SDA-500)^4^. The contigs for the Pakistani strain from this report were produced using short read sequences from Illumina sequencer with libraries ranging from small to medium and large insert sizes with a coverage of about 100X and read length of 101 bp. From this, the scaffold level assembly for the Pakistani strain of *An. stephensi* was reported by assembling reads with ALLPATHS-LG (P)^4^. The study reported an N50 of 0.8 Mb and the L50 of 85.

Yet another valuable resource for *An. stephensi* is the low-resolution physical map of the chromosomes^5^. This study reported a physical map consisting of 422 DNA markers hybridized to 379 chromosomal sites of the *An. stephensi* polytene chromosomes providing a resolution of 0.6 Mb. Of these, 241 are cDNA markers for which both location and sequences are accessible^3^.

In the past, experimentally derived mate-pair libraries of varying insert sizes have been used in assembling useful draft genomes for many species from short paired-end reads. Tools like SOAPdenovo use reads from mate-pair libraries to connect contigs from short paired-end reads into scaffolds based on the insert size information to create hundreds of draft genome assemblies. The quality of draft assemblies was mainly proportional to the insert size of the mate-pair libraries. It should be mentioned here that the gap between contigs measured by insert size of a given mate-pair read anchoring the two contigs is filled with Ns. However, the advent of cost effective long read sequencing technologies has made the creation of mate-pair libraries, an arduous step, obsolete. Long reads, along with technologies such as HiC, are generating high quality reference genomes of many non-model organisms. This has created an opportunity to provide chromosomal context to draft scaffolds, the holy grail of all genome assemblies and for other strains, cultivars and landraces at reduced cost.

Reference-guided improvement of draft genome assembly of individuals from the same species is becoming routine. Mate-pair libraries from one *Arabidopsis thaliana* strain were shared across many strains to build super-scaffolds for all individuals^6^. Also, assisted assembly of closely related species significantly improved the contiguity of low coverage mammalian assemblies^7^. For example, the draft genomes of four species including bush baby, African elephant, rabbit and guinea pig from the “Mammal24 - 2X” project were built using both human and canine references^7^. Reference-based assembly relies on DNA level homology between the reference and the draft genomes, which can only be expected if both are from the same species. In the absence of a reference genome from the same species, a draft assembly can be improved using the synteny and protein level homology between species to provide chromosomal context to scaffolds. Recently, a chromosome level genome of *Lates calcarifer* was assembled from a draft genome using long read sequencing, transcriptome data, optical/genetic mapping and synteny to two closely related seabasses^8^. In yet another report, 16 out of 60 chromosomes of the Tibetan antelope were reconstructed from draft assemblies using its homology to cattle^9^. In fact, using independent mapping data and conserved synteny between the cattle and human genomes, 91% of the cattle genome was placed onto 30 chromosomes^10^. In a review article, synteny has been used to filter, organize and process local similarities between genome sequences of related organisms to build a coherent global chromosomal context^11^. Similarly, the malarial strain, *Plasmodium falciparum* HB3, was improved using the reference of *P. falciparum* 3D7 combined with an assisted assembly approach that significantly improved the contiguity of the former^7^. More recently, it has been shown that the scaffolds from draft genomes of 20 *Anopheles* species could be improved considerably by providing chromosomal context using synteny to each other^12^.

Here, we have created a near-chromosome level assembly of *An. stephensi* Indian strain maintained at Walter Reid (IndV3s) using draft assemblies of two strains downloaded from VectorBase.

## Results

### Homology-based assembly and pseudomolecule generation

We utilized contigs/scaffolds from draft genomes of two different strains of *An. stephensi* to iteratively improve the draft assemblies of both obtained from public repositories using complementary information from one to the other. Table 1 describes the assembly metrics through the iterative improvement process. The initial assembly of Indian strain (IndV1) downloaded from VectorBase had a L50 of 37 and that for SDA-500 (PakV1) assembly was 85. In the first iteration, the L50 of IndV1 was reduced to 19 (IndV2) with the help of simulated mate-pairs of varying insert sizes from PakV1. During the second iteration, the L50 of PakV1 was reduced to 22 (PakV2) using the simulated mate-pairs of varying insert sizes from IndV2 of the assembly of Indian strain obtained after first iteration. The subsequent iteration resulted in assemblies with L50 of the Indian strain dropping to 9 from 37 and that for the Pakistani strain to 12 from 85. Further iteration did not show significant improvement suggesting saturation of complementing information in the initial draft assemblies of the two strains.

**Table 1:**
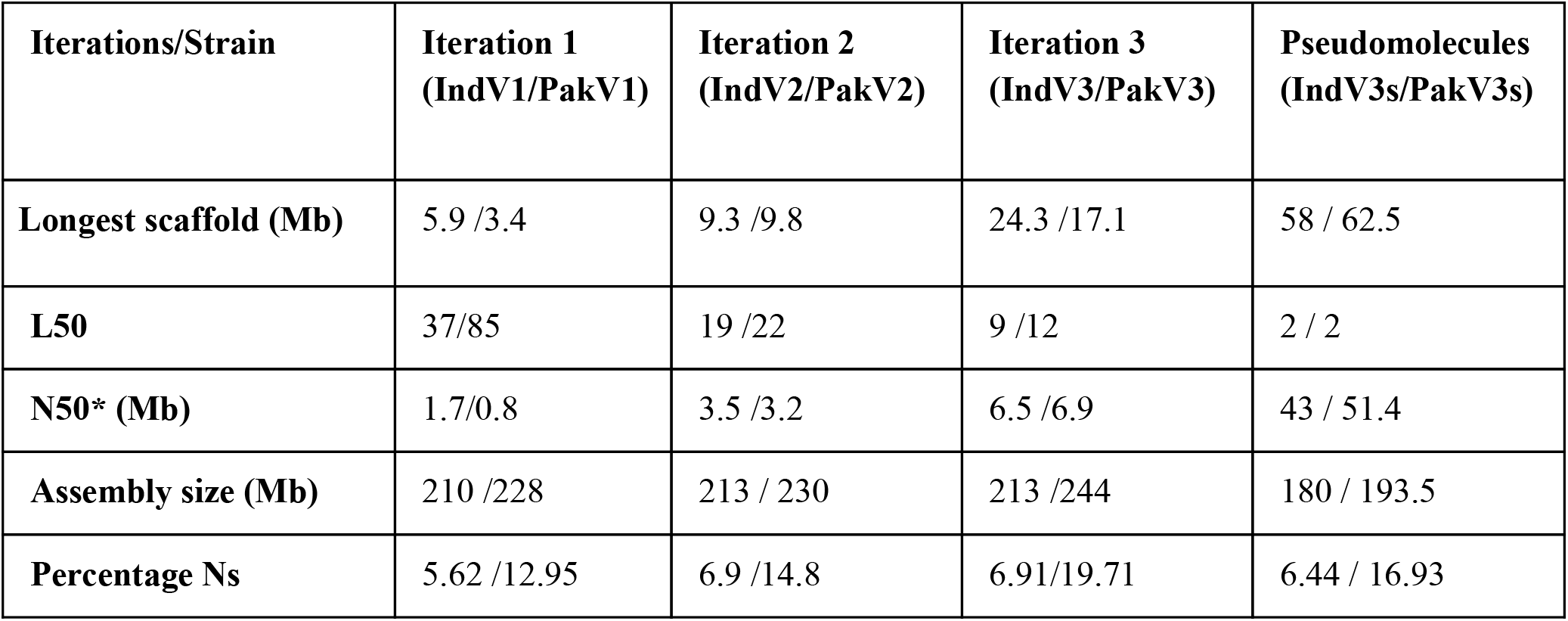
Homology-based iterative assembly. Improvement in assembly metrics for Indian and Pakistani strains after every iteration. *N50 is the scaffold length, x, such that 50% of the genome is assembled on scaffolds of length x or longer.

In the absence of HiC data and/or optical mapping data, the published physical mapping data was used to order the scaffolds and stitch them into pseudomolecules. It is important to note here that the scaffolds were long enough for super scaffolding using the placement of 230 out of the 241 cDNA markers^5^ for which the sequences were available^3^. Out of 230 markers, 209 (90%) could be mapped with confidence to scaffolds. Figure 1a shows the overlap between sequences from the physical mapping information for *An. stephensi* and the scaffolds generated from the third iteration of homology-based assembly for both the Indian (IndV3) and Pakistani strains (PakV3) of *An. stephensi*. The almost null overlap of physical maps placed on scaffolds, as shown in Figure 1a, helped in uniquely assigning chromosomes to scaffolds. However, physical markers from multiple arms hit a few scaffolds challenging the assignment of chromosomes for some long scaffolds (see Supplementary Table S6). As shown in Figure 1b, the scaffolds have been assigned orientation within the chromosomes, such as “f” (forward) and “r” (reverse), based on the linearity of physical maps. The equation for stitching scaffolds is elaborated along with the scaffold IDs (the numbers following ‘f’ and ‘r’). The karyogram in Figure 1c shows the final order and orientation of the physical markers on each chromosome after building the pseudomolecules. The total bases incorporated into pseudomolecules after stitching constituted more than 85% of the estimated genome size excluding Y chromosome.

**Figure 1:**
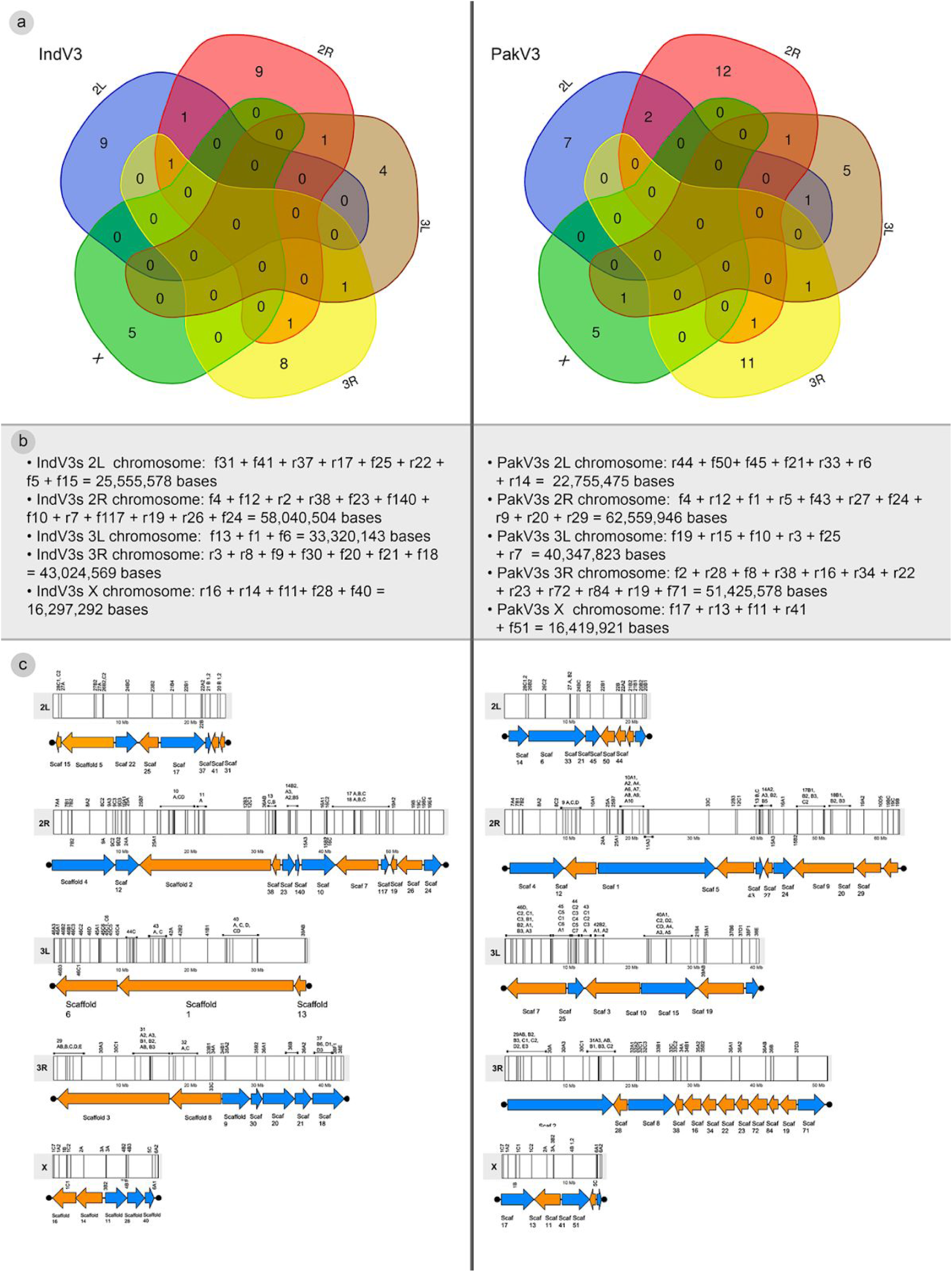
Generation of pseudomolecules. a) Intersection of scaffolds from IndV3 and PakV3 strains generated by homology-based assembly with marker data from physical maps for *An. stephensi*. b) Ordering of scaffolds into five chromosome arms based on the linearity of physical map markers using position and orientation for IndV3 and PakV3. c) Karyogram showing stitched near-complete chromosomes, IndV3s and PakV3s with DNA marker location on scaffolds from IndV3 and PakV3.

### Validation using synteny

The assemblies of IndV3s and PakV3s are validated using synteny to each other and synteny to other completed *Anopheles* genomes. In Figure 2, we show the synteny of IndV3s against PakV3s (2a), *An. gambiae* (2b) *and An. funestus* (2c). The synteny of the X-chromosome and the 3L arm of IndV3s shows the least variation with PakV3s assembly, suggesting that the overall assembly quality is good. The not-so-perfect synteny between the two strains of *An. stephensi* in 2R, 2L and 3R may reveal real differences between the genomes of the two strains and/or potential mis-assembly. Synteny of chromosomal arms 2L and 3L between IndV3s and PakV3s with insertions in IndV3s suggests incomplete assembly of PakV3s as expected because of the lack of long reads in the original SDA-500 assembly (Figure 2a). The synteny between chromosomes of IndV3s against *An. gambiae* and *An. funestus* suggests a near complete assembly of IndV3s without major gaps (Figure 2b and 2c). Figure 2 also shows potential intra-species (2d) and inter-species (2e, 2f) inversion events.

**Figure 2:**
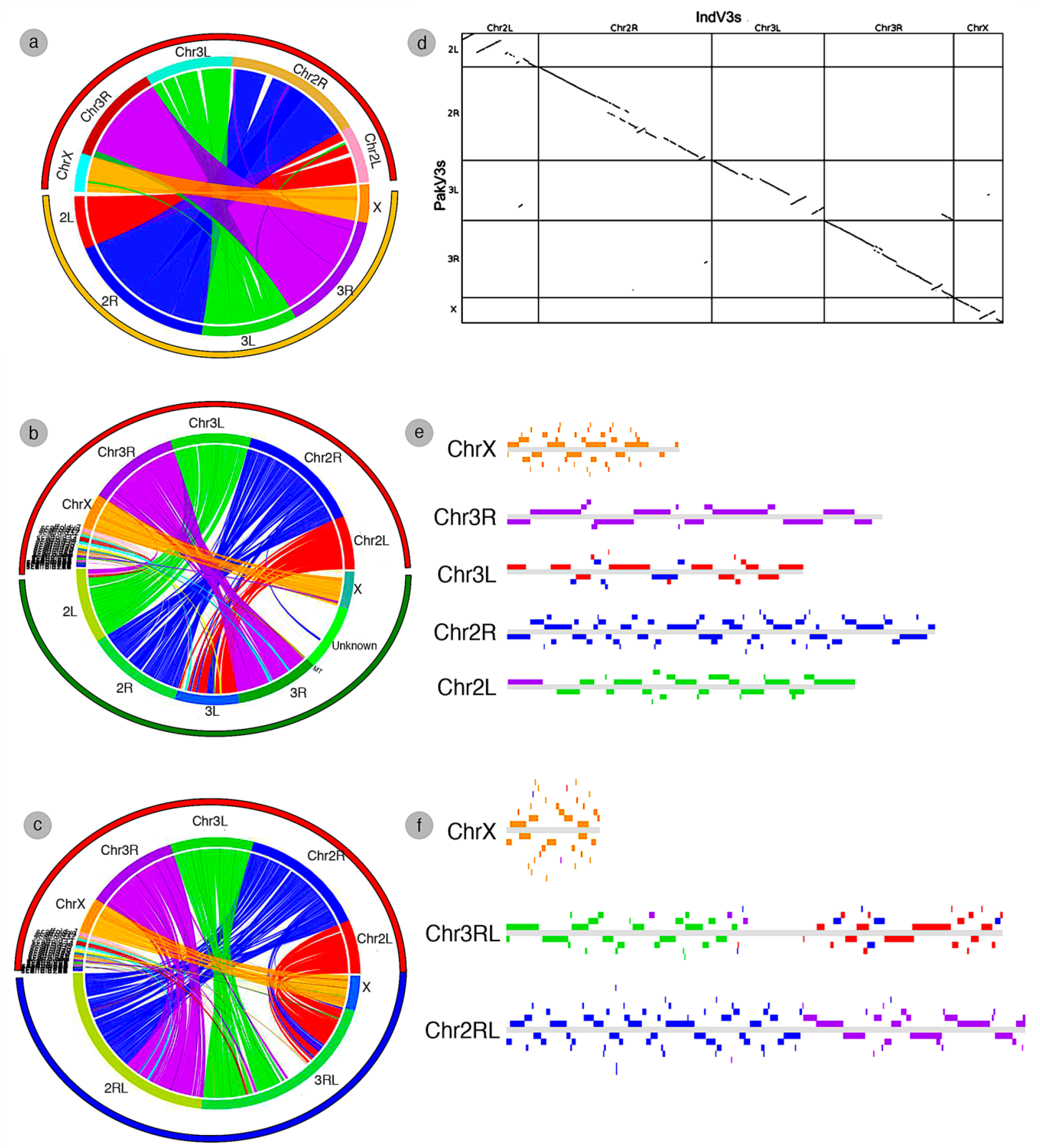
Assembly validation using synteny. a) Circos plot showing synteny between IndV3s (brown arc) against PakV3s (yellow arc) b) Synteny of IndV3s (brown arc) against *An. gambiae* (green arc) and c) Synteny of IndV3s against *An. funestus* (blue arc). d) represents the dot plot of IndV3s on PakV3s. e) and f) represent block chromosomal diagrams of IndV3s on every chromosome of *An. gambiae* and *An. funestus* respectively.

The scaffolds classified as unknown in *An. gambiae* genome, which may represent Y-chromosome and/or microbial genomes, is not syntenic with unplaced scaffolds of IndV3s suggesting the possibility of unique microbial compositions in *An. stephensi* compared to *An. gambiae*. However, several unplaced scaffolds from IndV3s are syntenic to *An. gambiae* and *An. funestus* chromosomes, suggesting that they are part of *An. stephensi* chromosomes but could not be placed on chromosomes in IndV3s assembly because of the low-resolution physical map (Figure 2b, 2c). The synteny of IndV3s with the rather high-resolution assembly of *An.funestus* with chromosomal arms assembling into a single scaffold reveals that the centromeric region between the two R and L arms of chromosome 3 is much larger than that of chromosome 2 (Figure 2c). Interestingly, in *An. funestus* the R arm of chromosome 3 maps to 3L arms of *An. stephensi* and 2L of *An. gambiae* respectively. Also, the 2L arm of *An. funestus* is syntenic with 3R arms of both *An. stephensi* and *An. gambiae*. Thus, chromosomal arm switching (twisting) must be common among *Anopheles* genus. However, high synteny between arms show limited gene flow between chromosomal arms.

Physical markers from the end of both 3R and 3L arms of *An. stephensi* hit scaffold18 of length 3.8 Mb of IndV3 and scaffold19 of length 4.2 Mb of PakV3 and hence had to be further resolved using numbers/linearity of markers and level of homology to marker DNAs. After resolving the conflict between 3R and 3L, scaffolds were assigned to the end of chromosomal arm 3R. Upon creating synteny between scaffold18 with chromosomes 3 and chromosome 2 of *An. funestus* (Supplementary Figure S1), it is clear that scaffold18 in IndV3 and scaffold19 of PakV3 joins 3R and 3L arms of chromosome 3 of *An. stephensi*.

### Completeness and annotation

In this section, we evaluated the overall completeness of the two assemblies IndV3s and PakV3s using diverse approaches. These include i) percentage of physical markers hitting the scaffolds, ii) total predicted proteins, iii) a set of elite proteins from BUSCO, iv) synteny to other *Anopheles* genomes, summarized in Table 2. Additionally, we assessed the gene structures for a set of predicted genes from IndV3s to validate the quality of assembly with respect to gene loci (Table 3).

**Table 2:** Completeness of assemblies. Metrics of IndV3s and PakV3s using various approaches including physical markers, predicted proteins from Augustus, validated proteins from public databases and ATGC content from synteny analysis.

| Parameter | IndV3s assembly | PakV3s assembly |
| --- | --- | --- |
| **Completion based on physical map** | 87%<br>21/28 (75%) on 2L, 84/88 (95%) on 2R, 52/56 (92%) on 3L, 37/40 (92%) on 3R, 15/18 (83%) on X | 84%<br>21/28 (75%) on 2L, 74/88 (84%) on 2R, 50/56 (89%) on 3L, 36/40 (90%) on 3R, 15/18 (83%) on X |
| **Predicted proteins** | 21,378 (20,318 in *An. funestus*) | 20,083 |
| **Validated proteins** | 12,148 | 11,303 |
| **BUSCO (out of 1013 *Arthropoda* specific)** | 98.2% including extra scaffolds<br>90.2% in chromosomes | 85.3% including extra scaffolds<br>82% in chromosomes |
| **Total ATGC content based on synteny to *Anopheles* genomes** | 168/180 million bases (93%) | 160/193 million bases (84%) |

**Table 3:**
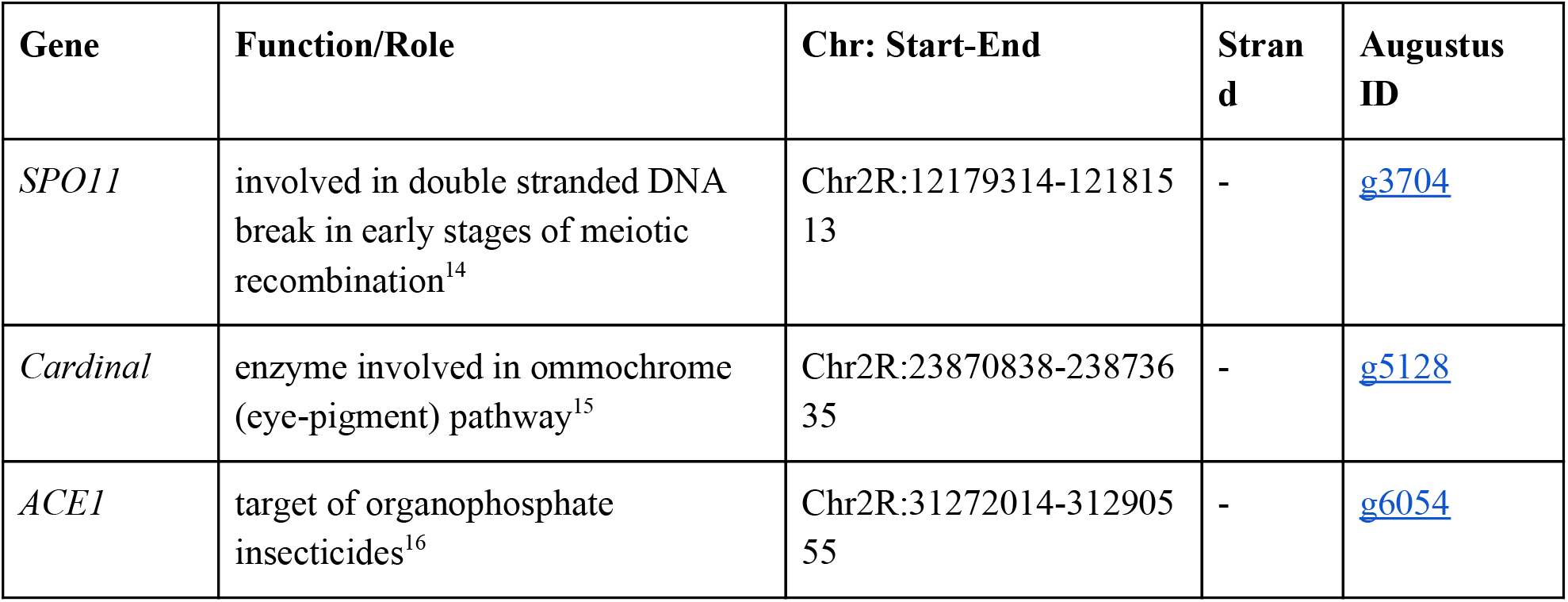
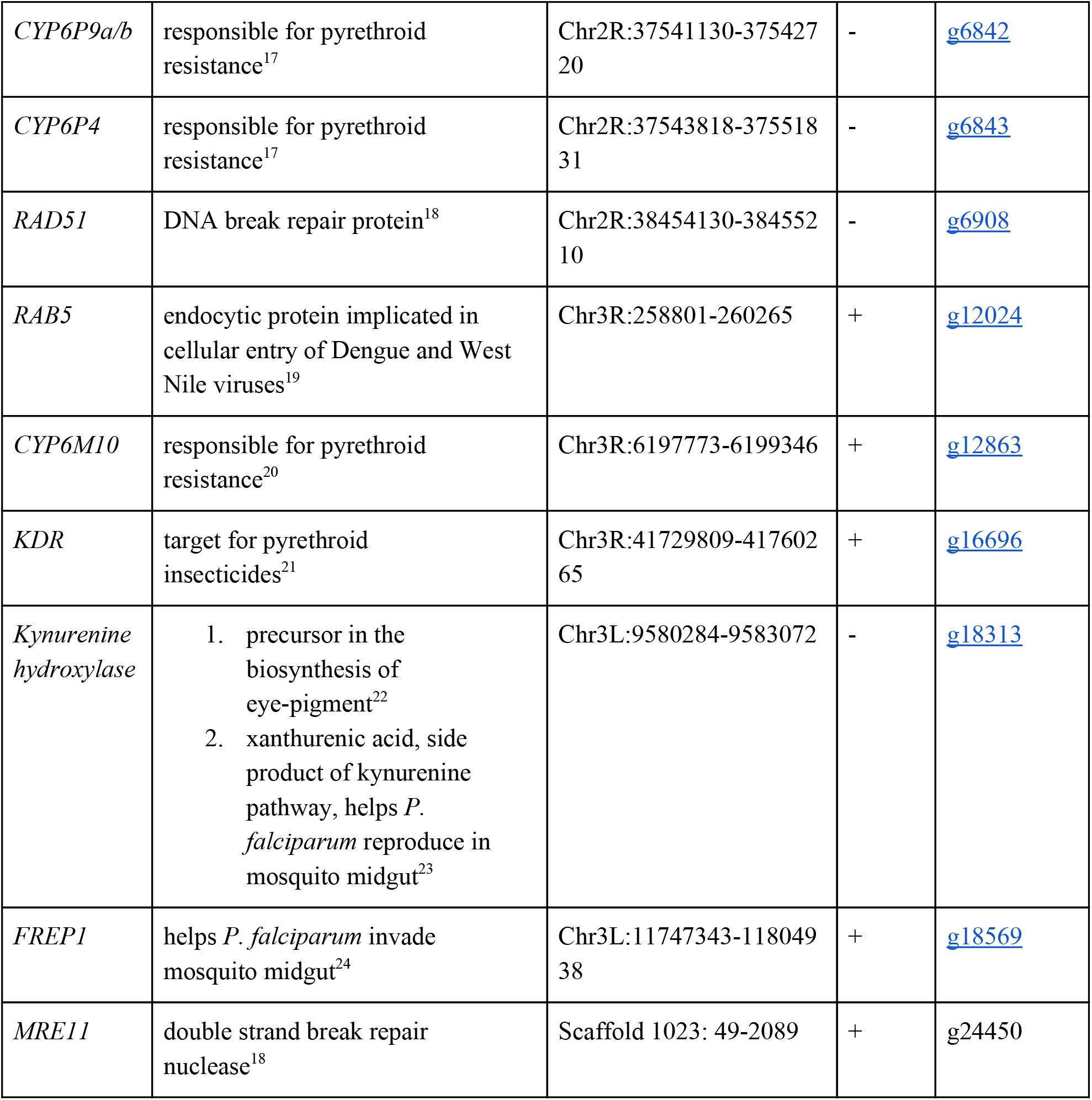
Genes of interest from a functional perspective in IndV3s. Details of genes along with their function, chromosomal context in IndV3s and a link to the gene structure is provided in the column with their Augustus ID.

The number of physical markers mapped to each assembly, 87% to IndV3s and 84% to PakV3s, provides a macroscopic measure of the completeness of each chromosome. For example, 15/18 markers (83%) from X chromosome mapped to IndV3s assembly and 84/88 (95%) of 2R, 37/40 (92%) from 3R, 52/56 (92%) from 3L and 21/28 (75%) markers from 2L were mapped to IndV3s assembly. Furthermore, a total of 180 Mb and 194 Mb of scaffolds from IndV3 and PakV3 (Figure 1b) could be placed on the chromosomes using physical markers to obtain IndV3s and PakV3s assemblies (Table 2).

Gene prediction using Augustus with *Aedes aegypti* as a model, the species closest to *An. stephensi* offered by the tool, resulted in the prediction of 21,378 and 20,083 proteins for IndV3s and PakV3s, respectively. These are comparable to the 20,318 proteins predicted in-house from the high-resolution chromosome-level assembly of *An. funestus* using exactly the same method. The proteins were validated using both publicly available transcriptome data from *An. stephensi* and orthology to the proteomes of other species. For transcriptome-based validation midgut transcriptome and Indian peptides database from VectorBase was used and for orthology, predicted genes from other available *Anopheles* genus in Swiss-Prot and TrEMBL was used. Table 2 shows the number of predicted proteins in IndV3s and PakV3s, respectively. Supplementary Figure S2 depicts the intersection of predicted proteins for IndV3s (S2a) and PakV3s (S2b) validated by each database. Of the 21,378 genes predicted for IndV3s, 12,148 are validated using one and/or all the above databases. For PakV3s, 11,303 of the 20,083 proteins predicted by Augustus were validated using these databases. An annotated GTF file is made available via the genome browser site listed under Data Availability section.

The completeness of the proteome in IndV3s and PakV3s assemblies was also assessed with BUSCO, which utilizes evolutionarily informed expectations of gene content. Out of the total of 1013 complete single-copy elite marker genes for *Arthropoda* used by BUSCO, 90.2% were placed on chromosomes and 98.2 % in all scaffolds of IndV3s, which is a very good indication of the completeness of the assembly.

### Genes of interest to vector biologists

To assess the completeness of assembly across gene loci, we assessed gene structures of selected predicted genes across IndV3s. The selected genes are of importance based on functional implications in the development of antimalarial strategies. These include genes that are implicated in insecticide resistance, parasitic infection and other DNA break repair mechanisms as listed in Table 3. Based on the well-studied orthologs, we found that the gene structure for all the genes of interest (see Table 3) is predicted at full length. For the FREP1 gene, no ortholog could be found and hence, the amino acid sequence for the domain was taken from Nui et al. to obtain a predicted gene with 214 amino acid (463-677) said to be conserved within this genus^13^. The links listed for Augustus ID in Table 3 can be used to view the gene structure on the browser and download sequences. For example, all 20 exons are predicted intact for the *KDR* gene with exon structures and sizes similar to orthologs from other species. With the exception of the *MRE11* gene, all are found in assembled chromosomes. The protein sequences of the genes listed below are given in Supplementary Text S2. The details of the online source of these sequences are given Supplementary Table S1.

### Developmental transcriptome analysis

Developmental transcriptome data for various stages from embryonic to adult stages were obtained from public sources (SRP013839, Supplementary Table S2). The coverage of Illumina data for each stage varied from 10X to 35X and is computed based on the assumption that 20% of the genome is transcribed. Transcriptome data summarizing sequencing parameters and the various developmental stages including embryonic stages 0-1 hour, 2-4 hours, 4-8 hours and 8-12 hours, larva, pupa, adult female, adult male, PBM (ovary Post Blood Meal), PEM (ovary post emergence), FeCa (female carcass) are detailed in Supplementary Table S2. The mapping was done on the IndV3s reference genome and the mapping percentages are given in Supplementary Table S3.

It is interesting to note that all the genes of interest listed in Table 3 show stage specific expression (Figure 3a). For example, *CYP6M10a, CYP6P4* and *CYP6P9a/b* are highly expressed in the larva stage compared to all other stages (Figure 3a3, 3a4, 3a5). On the other hand, the *KH* gene shows low expression in larva and adult males (Figure 3a8). Genes *ACE1, FREP1, KDR*, and *SPO11* show very low overall expression as can be seen by the range in the Y-axis (Figure 3a1, 3a6, 3a7 and 3a12). However, *SPO11* expression is higher in embryonic early stage (Figure 3a12) and in post blood meal ovary. *ACE1* and *KDR* show detectable peaks in adult male (Figure 3a1, 3a7, Supplementary Figure S5) and larva. Among other interesting findings is that *MRE11* and *RAD51*, both implicated in DNA repair, are upregulated in post blood meal ovary and the profiles of these genes correlate well across stages (Figure 3a9, 3a11). The genes *cardinal* and *KH* show negative correlation except in adult female, with *KH* upregulated in PBM and PEM stages and downregulated in adult male (Figure 3a2, 3a8). The gene *cardinal* is upregulated in adult male and is downregulated in PBM and PEM stages (Figure 3a2).

**Figure 3:**
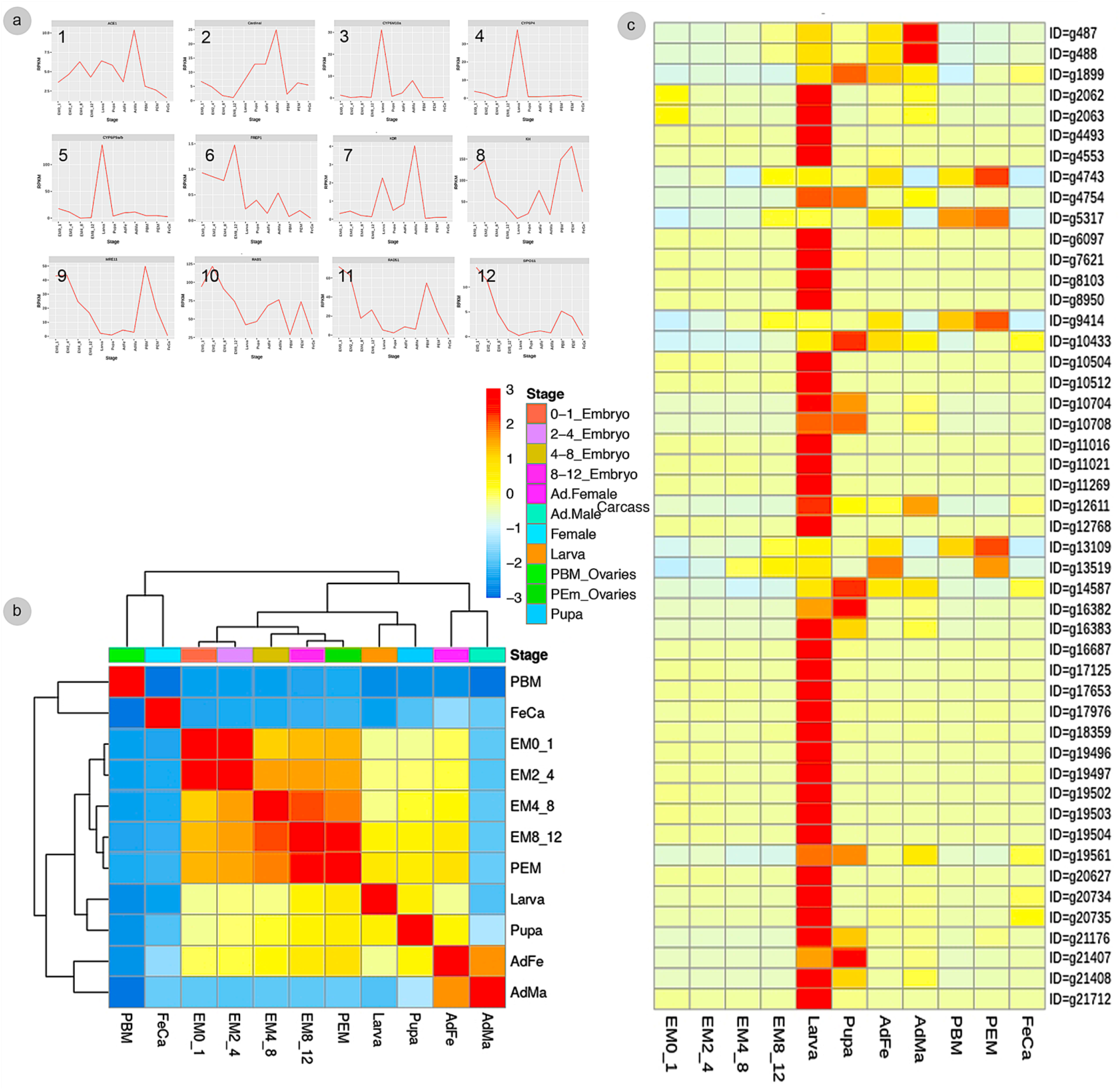
Transcriptome analyses using IndV3s reference genome and gene annotations. a) Expression profiles across developmental stages of genes of interest listed in Table 3, b) Correlation of genome-wide gene expression between different developmental stages, c) Heatmap of gene expression sorted by larva.

Genome-wide gene expression patterns show that adult females correlate with pupa and larva stages while adult male, PBM and FeCa show no correlation to early developmental stages (Figure 3b). As expected, there is good correlation between various embryonic stages (Figure 3b). Also, the overall gene expression profile of adult male and adult female correlates well without any correlation to female carcass (FeCa). Genome-wide gene expression profiles also provide opportunities to discover stage-specific novel target genes. For example, Figure 3c shows genome-wide gene expression profile sorted descending order of gene expression (RPKM) in larva, suggesting stage-specific expression of several genes in each stage. Gene expression data sorted by other stages are provided in a link listed under Data Availability.

### Evolution of Olfactory receptors (ORs)

Olfactory receptor genes (ORs) are of interest from an evolutionary point of view in vectors because of their role in host choice and therefore disease transmission. The number of ORs in different vectors vary widely. *Ae. aegypti* has 110 ORs^25^ whereas *An. gambiae* encodes for only 79 ORs^26^. The total number of OR proteins predicted from IndV3s that are orthologous to *An. gambiae* ORs is 54. Using a similar approach, we found 42 unique ORs from the recently reported high-quality genome of *An. funestus*^2^. The protein sequences of the 54 ORs from *An. stephensi* and 42 from *An. funestus* (Supplementary Text S1) are used to create a phylogenetic tree (Figure 4), which shows the orthology of the 54 *An. stephensi* ORs with respect to 42 from *An. funestus*. The corresponding orthologs of *An. gambiae* ORs are in green at the branch points. Supplementary Figure S3 shows a phylogenetic tree including orthologs of ORs from all three *Anopheles* species. Supplementary Table S7 shows the BLAST-based orthology used in decorating the tree in Figure 4 with *An. gambiae* accessions, which is consistent with the ClustalW based tree presented in Supplementary Figure S3.

**Figure 4:**
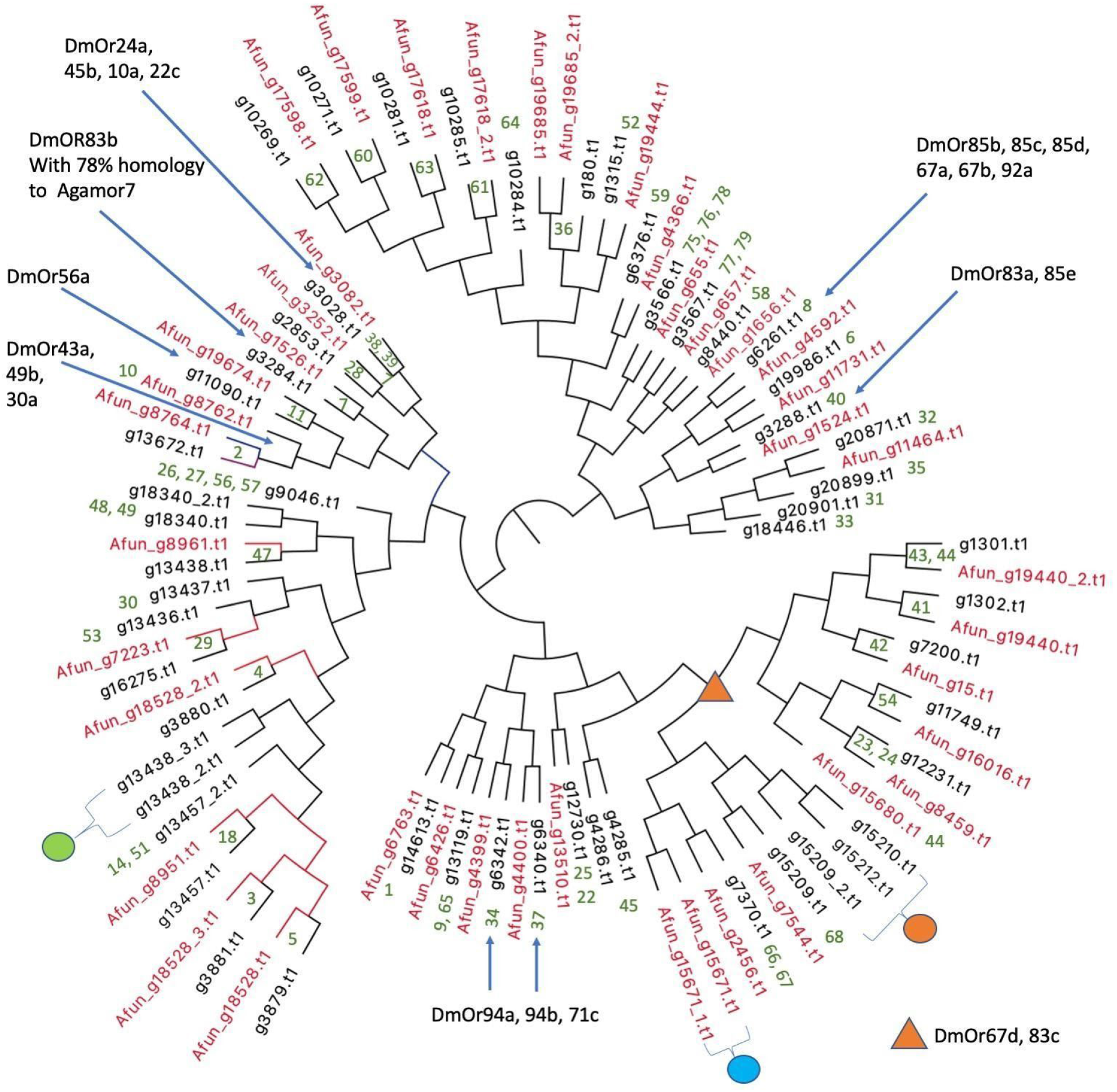
Phylogenetic relationships of olfactory receptors. The 54 ORs of *An. stephensi* (black) predicted from IndV3s with the 42 ORs predicted for *An. funestus* (red). Embedded numbers in green are the accession IDs of *An. gambiae* ORs. Green dot represents ORs missing in *An. funestus* and expanded into 5 ORs in *An. gambiae* including accessions 13, 15, 16, 17, 55. Cyan dot represents ORs missing in *An. stephensi*, duplicated in tandem in *An. funestus* and duplicated in *An. gambiae* with accessions 43-44. Orange dot is missing in *An. funestus* and expanded into 6 ORs in *An. gambiae* with accession IDs of 69-74. Blue arrows and orange triangle show ORs of *D. melanogaster* based on their orthology to *An. gambiae* reported elsewhere^26^.

The lower number of total ORs predicted from both *An. stephensi* and *An. funestus* is clearly at odds with the level of completion of both the genomes. Furthermore, in all three species some orthologous ORs appear in tandem in the genome, as reported previously^3^. For example, the 3 ORs from *An. stephensi* with accession IDs of i) g13438 with orthology to OR 47 of *An. gambiae* and 8961 of *An. funestus*, ii) g13438_2 and iii) g13438_3 are in tandem in the genome reported here. Orthologs for g13438_2 and g13438_3 from IndV3s are missing in *An. funestus* (see green dot in Figure 4) but expanded into six ORs in *An. gambiae* with accessions of 13, 15, 16, 17, 55 (Supplementary Figure S3). On the other hand, the orthologs of ORs 46 and 47 from *An. gambiae* are missing in *An. stephensi* but are present in tandem in *An. funestus* (cyan dot in Figure 4) with accessions Afun_g15671 and Afun_g15671_1. The orthologs of *An. stephensi* with accessions of g15209_2, g15210 and g15212 (orange dot in Figure 4) are missing in *An. funestus* but are collinear orthologs of 6 ORs in *An. gambiae* with accessions ranging from 69 to 74 (see highlighted in orange in Supplementary Figure S3). There are orthologs for 5 *An. gambiae* ORs with accession IDs of 12, 19, 20, 21, 50 missing in both *An. stephensi* and *An. funestus*, which cluster into a group (see highlighted in green in Supplementary Figure S3) and is also reported missing in *D. melanogaster*^26^. While this could result from potential incomplete genomes of *An. stephensi*, considering that the *An. funestus* reference genome is of very high-resolution, it is unlikely that the orthologs to the same genes would be missing in the two independently reported near completed genomes and hence can be considered unique to *An. gambiae*.

A majority of the 54 OR genes in *An. stephensi* show higher expression in early embryonic stages (Supplementary Figure S4). Interestingly, the relatively lower number of ORs that are upregulated in other stages are not upregulated in embryonic stages. The three ORs, g10271, g10269 and g3284 of *An. stephensi*, which are highly expressed in pupa are orthologous to genes GRPor60, GPRor62 and GPRor7 from *An. gambiae*. Interestingly, GPRor7 of *An. gambiae* shows high homology to an OR coreceptor, DmOR83b. The gene g13672 of *An. stephensi with* increasing expression from pupa to adult female to adult male is an ortholog of GORor2 from *An. gambiae* and DmOR43a of *D. melanogaster* that has a known ligand^27^.

### Genetic diversity among individuals using whole genome sequencing

We sequenced at 30X coverage, 6 lab female individuals from one urban city and 1 male individual from another city to estimate the diversity among individuals compared to IndV3s reference. The number of variants in these individuals with respect to IndV3s reference is roughly 2.6 million suggesting an overall SNP density of 1 every 68 bases of the genome compared to 1 in 1,000 for the human genome. This is ten times higher than the number of SNPs (319,751) reported for *An. stephensi* using diverse individuals from a lab strain, which may be because the individuals were taken from the same colony that was used in creating the reference^3^. X chromosome has the lowest SNP density with 1 in 85, which is in tune with what is reported for *An. gambiae* and *An. stephensi*^3,28^. In Table 4, we show SNP density for X chromosome from both female and male individuals to make sure the lower SNP density in X is not resulting from lower coverage of X from hemizygous male. In females, the SNP density is at 1 in 85 bases and is only slightly higher than the SNP density of 1 in 114 bases in male.

**Table 4.**
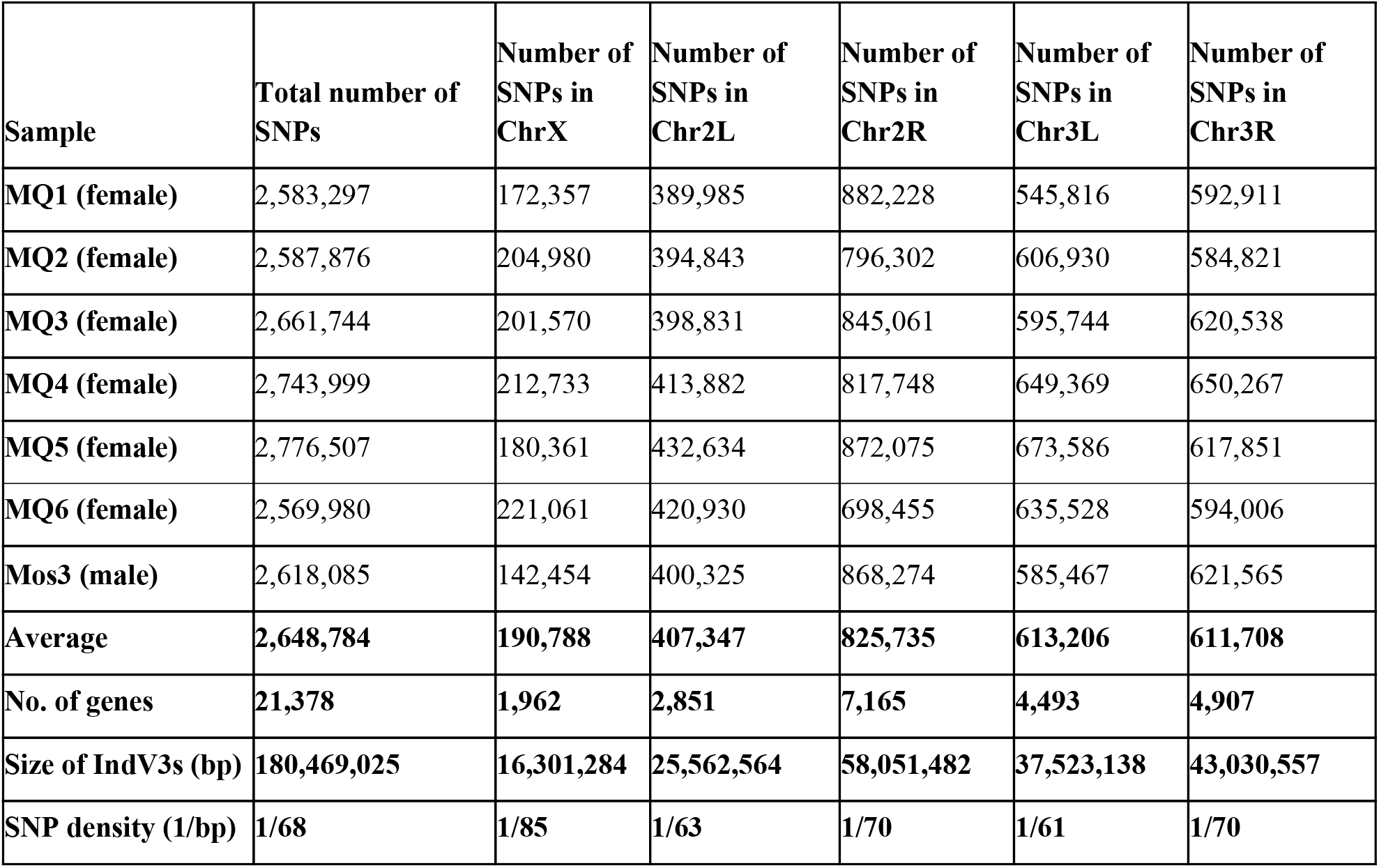
SNP analyses from whole genome sequencing. Distribution of SNPs, SNP density across chromosomes of *An. stephensi* from 7 individuals

## Discussion

Here, we report a near-chromosome-level assembly of *An. stephensi*, a major malaria causing vector in urban India. Draft assemblies of two strains of *An. stephensi* were obtained from VectorBase and their complementarity was used to improve the assembly of each strain using simulated mate-pairs from the other in an iterative fashion. The L50 for the Indian strain improved from 37 to 9 after iterative improvement and to 2 after building pseudomolecules. The longest scaffolds increased in lengths from 6 Mb to 24 Mb to 58 Mb for IndV3s. Considering that the genome size is roughly 225 million bases with 3 chromosomes, the assembly statistics achieved are good enough to be validated and stitched into chromosomes using physical markers^5^. The total genome size after stitching is 180 Mb and 194 Mb for Indian and Pakistani strains, respectively, covering more than 80% of the estimated genome size. Considering that there is a low level of representation from the heterochromatin region of the genome and that the Y chromosome remains unassembled, placing the scaffolds with lengths adding up to 180 million bases, covering 80% of the estimated genome size, onto chromosomes suggests much higher representation of euchromatin regions of the chromosome assembled here. Furthermore, the lower percentage in the assembled genome is also resulting from the low resolution of the physical mapping data, which may be missing in shorter scaffolds (Figure 2b). The linearity of the final chromosomes was validated using synteny to each other and to the chromosomes of *An. gambiae and An. funestus*. The full synteny between the X chromosomes of IndV3s and PakV3s suggests potentially fewer mis-assemblies. There are no major gaps in the synteny between IndV3s and chromosome-level genomes of *An. gambiae* and *An. funestus* (Figure 2b/2e and 2c/2f) suggesting near-chromosome level assembly of IndV3s. Also, synteny of IndV3s to *An. gambiae* and *An. funestus* showed the least shuffling of genes between the chromosomal arms of the three malaria vectors compared here. However, arm switching appears to be common among species in the genus *Anopheles*.

The assembly reported here and submitted to the NCBI repository as early as January 2019 is more complete than the one reported recently based on synteny across closely related species^12^. We observed (Figure 2c) that synteny across closely related species, such as *An. stephensi* and *An. funestus*, can vary significantly, thus questioning the use of synteny alone to assign scaffolds to chromosomes. Since our assembly used only components from various strains of the same species and used protein-level synteny only as a measure of completeness, we believe that our assembly is more likely to be authentic.

We used multiple approaches to assess the completeness and usefulness of IndV3s and PakV3s assemblies reported here. These include assessing i) the percentage of physical maps located on the chromosomes, ii) the percentage of predicted genes validated via databases, iii) gene structure validation using genes of interest, and iv) synteny with other *Anopheles* genomes. Using Augustus gene prediction software with *Aedes aegypti* as a training model, we were able to predict 21,378 genes in IndV3s and have validated 12,148 using multiple databases including transcriptome data from the same species and proteome data from other species. The number of predicted and validated proteins from IndV3s is in line with the corresponding numbers for *An. funestus* with ~20,000 predicted genes, the most closely related to *An. stephensi™* for which the genome is recently deciphered. To assess the robustness of the assembly around gene loci, we looked for completeness of a set of functional genes of interest to vector biologists. All selected genes were predicted full length as validated by the gene structures of the orthologs from other well studied species. Almost all the genes showed conservation in the size of orthologous exons within the folding domain. From the expression profiling data on the browser, it is clear that there are two extra cassette exons predicted for FREP1 gene with no evidence of expression in the stages discussed here and for KDR, a number of cassette exons are missing in the prediction but show expression in adult male (Supplementary Figure S5). Since skipping and insertion of these cassette exons have no effect on the reading frame towards the 3’ end of the exons, they may be skipped or included under different biological contexts not studied here. Table 3 provides direct links to these gene structures on the genome browser (link under Data Availability section).

The end of chromosome 3R of IndV3s near the centromere built using scaffold18 needs special mention. The scaffold18 of IndV3 of length 3.8 Mb and the scaffold19 of PakV3 of length 4.2 Mb could not be uniquely assigned to chromosomal arms based on physical markers. After resolving conflict, the scaffolds were assigned to the end of chromosomal arm 3R. A synteny between scaffold18 of IndV3 with the two completed chromosomes of *An. funestus* (Supplementary Figure S1) shows that the first half of scaffold18 is syntenic to the middle of chromosome 2 and the second half to the middle of chromosome 3 of *An. funestus*. Since the 2L arm of *An. funestus* is syntenic to the 3R arm of *An. stephensi* and 3R arm of *An. funestus* is syntenic with 3L of *An. stephensi* (see table in Supplementary Figure S1), scaffold18 of IndV3 and scaffold19 of PakV3 is suspected of spanning the breakpoint/centromere between the two arms of chromosome 3 of *An. stephensi*. Considering that 3R and 2R arms of *An. stephensi* is syntenic to chromosome 2 of *An. funestus* with a very small centromere as revealed by lack of gap in the synteny (Figure 2c and 2f), we hypothesize that the centromere for chromosome 3 of *An. stephensi* is short enough for scaffold18 of IndV3 and scaffold19 of PakV3 to span across without the need for HiC data (Figure 5). This also explains why markers from both 3L and 3R arms were hitting these scaffolds. Thus, chromosome 3 could have been stitched full length with both arms connecting using scaffold18 using the approach presented here. However, in the assembly submitted to NCBI this scaffold is stitched at the end of arm 3R reaching out to 3L. Chromosomal arm switching may lead to reduced fitness and is proposed as a mechanism for speciation in blood-feeding black flies^30^. By analogy, it can be hypothesized that arm switching may have played a role in speciation in the genus *Anopheles*.

**Figure 5:**
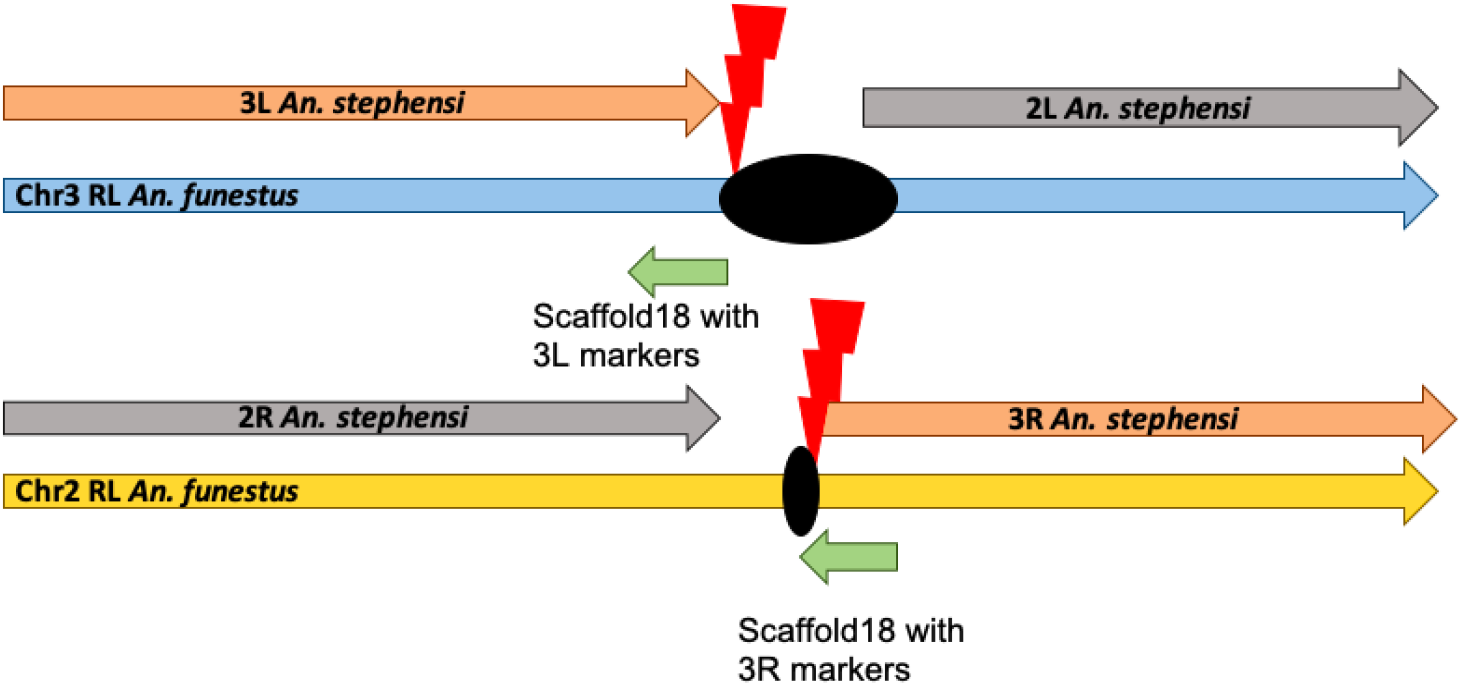
Comparative analysis reveals chromosomal arm switching. Arms of *An. stephensi* chromosomes syntenic with arms of *An. funestus* chromosome. Lightning symbols in red are the potential breakpoints and the ellipses in black are the centromeres.

The evolution of olfactory receptors is of interest among blood feeding mosquitoes. We have identified 54 ORs from the IndV3s assembly and 42 ORs from the recently reported genome of *An. funestus* using the ORs of *An. gambiae*^26^. These numbers are much lower than the 79 ORs predicted for *An. gambiae*. Considering the level of completion of the genomes of *An. stephensi* and *An. funestus*, this steep drop in the number of ORs predicted in these two species may suggest active/convergent evolution among OR genes. Also, the multiple OR genes that are found in tandem within the genomes of all three *Anopheles* species with some tandem OR cluster from one species orthologous to tandem OR cluster in the other, suggests multiple tandem duplication and expansion of OR genes in *Anopheles* genera, as reported for salivary genes^3^. The most conserved OR from *An. gambiae* with an accession of GPRor7 (Agamor7) to the OR from *D. melanogaster* with an accession of DmOr83b is also conserved in both *An. stephensi* (g3284) and *An. funestus* (AFun_g1526) suggesting a conserved function within *Diptera*. These are the olfactory coreceptor (orco), which constitutes a heterodimer with other ORs helping the odour molecules to enter into the insect body by forming a ligand-gated ion channel^31^. Also, the branch marked with orange triangle in Figure 4 with two orthologs in *D. melanogaster* has expanded to include 17 orthologs of *An. gambiae* and 10 orthologs for both *An. stephensi* and *An. funestus*, suggesting expansion of this function in *Anopheles* genera.

A chromosome-level genome also provides a great resource to study genome-wide gene expression across tissues and/or developmental stages. Genome-wide gene expression analysis is also an important source to clone full-length genes of interest for a non-model organism. We have taken advantage of the completed genome to profile genome-wide gene expression from early embryonic stages to adult mosquitoes using publicly available data. As shown in Figure 3b there is a good correlation between various embryonic stages including PEM. Also, while there is good correlation between adult male and female, only adult female retain correlation with all early developmental stages. Not surprisingly, there is no correlation between PBM and FeCa, as FeCa is carcasses left after removing post blood meal ovary (PBM). We observe that the selected metabolic genes of interest, implicated in pyrethroid resistant insects, *CYP6M10a*^32^, *CYP6P4*^33^ and *CYP6P9a/b*^34^ are highly expressed in the larva stage compared to all other stages (Figure 3a3, 3a4, 3a5). This is expected because the majority of insecticides are targeted to control larvae. Also, *ACE1* and *KDR* genes, which are implicated in insecticide resistance, is also differentially upregulated in larva with the exception of adult male where it is highly expressed (Figure 3a1 and 3a7, Supplementary Figure S5). Interestingly, mutations in *KDR* and *ACE1* genes have been shown to cost male fitness^35,36^. The expression profile of *SPO11* gene, with a role in meiosis, is detectably higher in embryonic early stage (Figure 3a12). *Cardinal* and *KH* genes, with the exception in adult female, display inverse correlation across developmental stages. On the other hand, the expression profiles of *MRE11* and *RAD51* genes (Figure 3a9 and 3a11), known to form complexes, are highly correlated with high expression in post blood meal ovary (PBM), perhaps suggesting a role in plasmodium infection.

Despite the importance of olfactory receptors in *Anopheles* evolution and host selection, the functions of individual ORs are not well understood except for a few with high homology to ORs from *D. melanogaster*. Gene expression profiles of the 54 ORs from *An. stephensi* reported here, show that the majority of them are highly expressed in embryonic stages (Supplementary Figure S4). Interestingly, the lower number of ORs upregulated in other stages are not upregulated in embryonic stages, suggesting distinct functions for many ORs during embryonic development. Of the three ORs (g10271, g10269, g3284) upregulated in the pupa stage, g3284 is an ortholog of the coreceptor DmOR83b of *D. melanogaster* conserved in all of Diptera, suggesting functional importance of g10217 and g10269 of *An. stephensi* during its pupal stage. The other OR gene g13672 of *An. stephensi* with increasing expression from pupa to adult female to adult male is an ortholog of DmOR43a of *D. melanogaster* with known a odorant molecule containing six-membered carbon rings including cyclohexanol (camphor like), cyclohexanone (found in human urine/blood), benzaldehyde (almond), and benzyl alcohol (fruits) with a single attached polar group without any effect on other molecules with six-member rings^27^. This would suggest that g13672 of *An. stephensi*, GRPor2 of *An. gambiae* and g8762/g8742 of *An. funestus* may be implicated in host recognition via cyclohexanone.

We believe that this is the first instance to obtain a chromosome level assembly from two draft assemblies of the same species using low-resolution physical marker data. This is also the first chromosome-level assembly of a malaria vector from India. The assembly reported here offers a genomic context to genes and other genetic elements providing a framework for controlling malaria in India with state-of-the-art technologies such as gene editing and gene drive.

## Methods

### Sources of data used in this work

The draft assemblies of both the Indian (IndV1) and Pakistani (PakV1) strains of *An. stephensi* were downloaded from https://www.vectorbase.org/. The sequences are also available on GenBank under the accession ALPR00000000.

The transcriptome data was obtained from public sources (NCBI SRA database: SRP013839).

Individual insects for whole genome sequencing was originally maintained at NIMR, Bangalore and were obtained from Dr. Sushant K. Ghosh (see acknowledgement).

### Homology-based assembly

The *An. stephensi* genomes of Indian strain (IndV1) and Pakistani strain SDA-500 (PakV1) were downloaded from https://www.vectorbase.org/.

Mate-pair libraries containing reads carry information about the arrangement/ordering of the chromosomes, which help to build scaffolds from the first pass assembly that is obtained from deep sequencing of paired-end reads. This implies that the sequence information from the mate-pair reads are NOT directly used in the assembly, but only the knowledge of ordering information is inherited.

#### a) Generation of read libraries

Using the samtools wgsim (version 1.9), mate-pair libraries with varying insert sizes (1 kb, 5 kb, 10 kb, 50 kb, 100 kb, 500 kb, 1 Mb, 2 Mb, 5 Mb) and read length of 50 bp were simulated from PakV1 with flags set for zero base error rate, rate of mutation and incorporation of indels.

#### b) Confident set of reads for subsequent scaffolding

To assess the quality of the reads generated from each library, they were mapped back against *An. stephensi* IndV1 genome using bowtie2 (version 2.3.5.1)^37^. Flags to suppress discordant and unpaired alignments were set. While processing the alignments, only paired reads mapped in the right orientation with the correct insert sizes were used for downstream analyses. Care was taken to remove multiple mapped reads from the concordant list of fastq reads.

#### c) Scaffolding of pre-assembled contigs

Reads from b) were used for super-scaffolding of the IndV1 genome to get the IndV2 assembly using the SSPACE tool (version 3.0)^38^. Now, from the IndV2 assembly, mate-pairs with varying library sizes (1 kb, 5 kb, 10 kb, 50 kb, 100 kb, 500 kb, 1 Mb, 2 Mb, 5 Mb) were simulated again to get the reads concordantly mapping to PakV1 assembly. These reads were used to super-scaffold PakV1 to obtain the PakV2 assembly. This iterative procedure was continued till the L50 and N50 of the assemblies did not improve significantly with further iterations resulting in assemblies IndV3 for the Indian strain and PakV3 for the Pakistani strain (shown in Figure 6).

**Figure 6:**
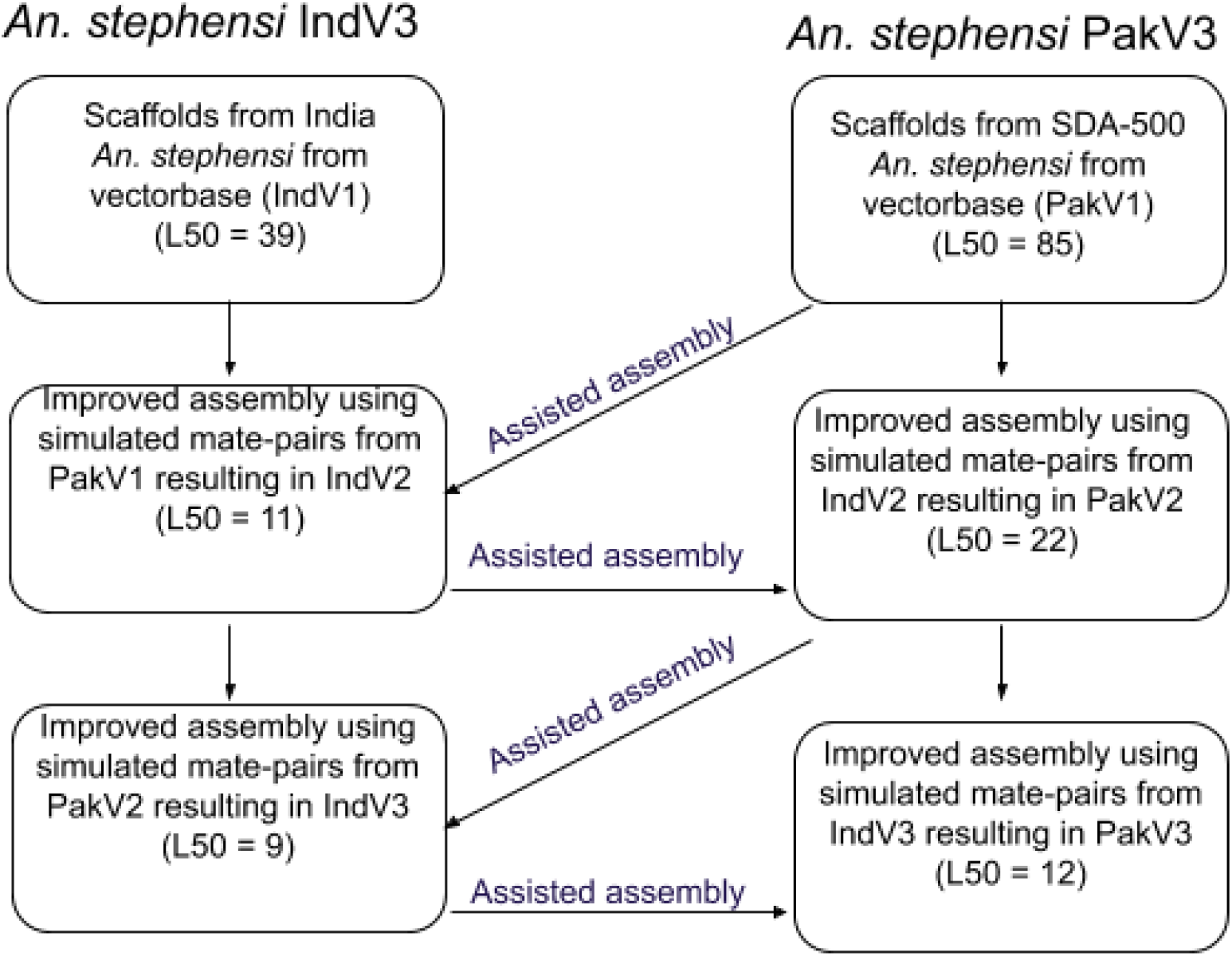
Schematic diagram showing the iterative approach followed to improve the assemblies of both strains of *An. stephensi*.

### Pseudomolecule generation using physical marker data

DNA sequences for physical map data for all chromosomal arms were downloaded from the supplementary material of *An. stephensi* draft assembly paper^3^. These were subjected to analysis using BLAST under stringent conditions (blastn -query 3L.fa -db indV3 -outfmt 6 -evalue 1.0e-25 -max_target_seqs 1 -out 3L_indV3.out) against the assembled genome IndV3. The unique hit of the physical markers onto scaffold IDs was used as a measure of the quality of assembly at this stage. A Venn diagram was generated to check the assignment of physical markers to scaffolds of the assembly (Figure 1a). The scaffold hitting against markers from two chromosomes is resolved using the number and linearity of markers, and in some cases using the level of homology at the DNA level. Chromosomes were put together after orienting the scaffolds using the order of the physical markers. 1000Ns were inserted to separate different scaffolds while stitching. Scaffolds in Supplementary Table S6 are those with doubtful orientation or overlapping markers.

The 5 chromosomal arms for the Indian strain are given below for a total genome size of 180 Mb, covering more than 80% of the estimated genome size.

- IndV3s 2L chromosome: f31 + f41 + r37 + r17 + f25 + r22 + f5 + f15 = 25,555,578 bases
- IndV3s 2R chromosome: f4 + f12 + r2 + r38 + f23 + f140 + f10 + r7 + f117 + r19 + r26 + f24 = 58,040,504 bases
- IndV3s 3L chromosome: f13 + f1 + f6 = 33,320,143 bases
- IndV3s 3R chromosome: r3 + r8 + f9 + f30 + f20 + f21 + f18 = 43,024,569 bases
- IndV3s X chromosome: r16 + r14 + f11+ f28 + f40 = 16,297,292 bases

The 5 chromosomal arms for Pakistani strain are given below for a total genome size of 194 Mb, again covering more than 80% of the estimated genome size.

- PakV3s 2L chromosome: r44 + f50+ f45 + f21+ r33 + r6 + r14 = 22,755,475 bases
- PakV3s 2R chromosome: f4 + r12 + f1 + r5 + f43 + r27 + f24 + r9 + r20 + r29 = 62,559,946 bases
- PakV3s 3L chromosome: f19 + r15 + f10 + r3 + f25 + r7 = 40,347,823 bases
- PakV3s 3R chromosome: f2 + r28 + f8 + r38 + r16 + r34 + r22 + r23 + r72 + r84 + r19 + f71 = 51,425,578 bases
- PakV3s X chromosome: f17 + r13 + f11 + r41 + f51 = 16,419,921 bases

### Physical map conflict resolution and validation using synteny

The scaffolds that were mapped to markers from multiple chromosomes were assigned to unique chromosomes based on the number of mapped markers, percent identity, e-value and alignment lengths. For example, Supplementary Table S6 shows scaffolds of IndV3s (ScaffoldID) assigned to markers from multiple chromosome arms (ChromosomeID) and that were resolved as shown in column Assignment. All the validation by synteny of IndV3s reference genome assembly against PakV3s, *An. gambiae* and *An. funestus* genomes were done using SyMAP (version 4.2)^39,40^.

### Gene annotation

AUGUSTUS (version 3.2.3)^41^, a eukaryotic gene prediction tool, was used to find protein-coding genes in the IndV3s genome. AUGUSTUS uses *ab initio* gene prediction and reconciles the predicted gene structures with orthology to the proteome from a model organism. The model organism closest to *An. stephensi* of that was made available by AUGUSTUS for use in gene prediction is *Aedes aegypti*. AUGUSTUS provides a gff3 file delineating the exon-intron boundaries for each predicted gene along with its protein sequence. This gff file is uploaded to the browser, the link to which can be found under the Data Availability section. The same gff3 file is used to compute gene expression profiles of all these genes across developmental stages.

### Identification of genes of interest

We used orthologs of the genes of interest from public databases by searching for the gene sequences from the closest available species. For the majority of genes, sequences were available in *An. gambiae*. However, for the FREP1 gene, no genes could be found in any related species. We took the amino acid sequence of the fibronectin domain of FREP1 from Nui et al. and found the full length predicted gene containing this domain. The table presented in Supplementary Table S1 shows the database source, accession and related organism used in the identification of these genes in *An. stephensi*.

### Finding orthologs of ORs

To find the ORs from *An. stephensi* and *An. funestus*, fasta sequences of 79 OR genes reported for *An. gambiae* is used^26^. Here, protein blast, blastp (version 2.7.1) was used for identifying orthologous genes from the predicted proteomes of *An. stephensi* and *An. funestus* using the 79 OR protein sequences from *An. gambiae* as query. The top hits are given in Supplementary Table S7. The amino acid sequences of ORs from *An. stephensi* and *An. funestus* were extracted from the respective proteomes and tandem ORs were manually split and provided in Supplementary Text S1. Using clustalW (version 2.1), a multiple sequence alignment of ORs from only *An. stephensi* and *An. funestus* were generated, because these two species are more closely related to each other than *An. gambiae*^29^. Using Figtree (version 1.4.4) we created phylogenetic trees shown in both Figure 4 and Supplementary Figure S3.

### Transcriptome analysis

Developmental transcriptome data was downloaded from NCBI SRA (SRP013839). Mapping was done using transcriptome reads on IndV3s using STAR aligner (version 2.6) (Supplementary Table S3). Read counts mapped per gene were obtained and normalized RPKM was computed providing a normalized expression profile for all predicted genes across the developmental stages. The genes of interest and olfactory receptors were taken out from annotation files and used for plotting the line graph as well as in the heatmap. The entire transcriptome analysis was done using R programming. Packages used in the analyses are pheatmap, stats, corrplot, knitr, ggplot2, dplyr, tidyr, reshape2. A description of the functionality of each tool is given in Supplementary Table S4. For further details on the script and version of packages used, we provide a link to markdown transcriptome report under Data Availability section.

### Isolation of genomic DNA

Briefly, the insect tissue was homogenized, and the genomic DNA was isolated using Qiagen Genomic-tp (Qiagen). Later, fluorometric quantification of DNA was done using Qubit 2.0 (Invitrogen).

### Library preparation and sequencing

Whole Genome DNA libraries with an average insert size of 200 bp were made using NEBNext^®^ Ultra^™^ II DNA Library Prep Kit for Illumina^®^ (New England Biolabs, 2016) using the protocol recommended by the company. Briefly, around 50 ng of DNA was used for library preparation, DNA was sheared using Adaptive Focused Acoustic technology (Covaris, Inc.) to generate fragments of length around 200 bp. The fragments were end repaired, 3’-adenylated, ligated with Illumina adapters, and PCR enriched with Illumina sequencing indexes. The size selection was performed using solid-phase reversible immobilization (SPRI) beads (Agencourt AMPure XP Beads) from Beckman Coulter. The quality and quantity of the libraries were evaluated using Qubit (Invitrogen) and TapeStation (Agilent). The libraries were diluted and pooled with an equimolar concentration of each library. Cluster generation was done using cBot (Illumina) and paired-end sequenced on Illumina HiSeq 2500 platform using TruSeq SBS Kit v3-HS (200 cycle) (Illumina, San Diego, CA) following the manufacturer’s recommendations.

### SNP Analysis for genetic diversity

DNA samples for 7 individuals were extracted and sequenced for 30X coverage. Variant calling was performed using an in-house pipeline, which uses fastqc (version 0.11.5), bowtie2 (version 2.3.5.1), samtools (version 1.9), picard-tools (version 2.18.12), samtools (version 1.9) mpileup^42^ and bcftools (version 1.9) to extract variants. The resulting vcf files are filtered using bcftools (version 1.9)^43^ for variants above a quality of 10 and a minimum depth of 3 reads. Mapping percentages of the 7 individuals on IndV3s reference have been tabulated in Supplementary Table S5.

### JBrowse and Database

The entire web framework of the website was taken from MEGHAGEN LLC, which includes the pre-built platform with the capability of genome visualization, blast, and further bioinformatic analysis. The website is built using HTML, CSS & javascript. Current website is hosted on amazon cloud running Linux EC2 instance provided by Meghagen LLC. It uses inbuilt servers (Apache, Shiny) to establish user side connections.

JBrowse is open source, fast and full-featured genome browser built with JavaScript and HTML5. It is easily embedded into websites or apps but can also be served as a standalone web page^44^. JBrowse can utilize multiple types of data in a variety of common genomic data formats, including genomic feature data in bioperl databases, GFF files, BED files, and quantitative data in wiggle files^45^. The website for IndV3s here has been built using Apache/2.4.29 (Ubuntu), build 2020-03-13T12:26:16 and JBrowse version 1.16.8.

## Acknowledgments

TIGS India for funding researchers involved in this work and for funds for sequencing. Government of Karnataka for funding the computational infrastructure at IBAB. IBAB facility for providing computational infrastructure used in all data analysis reported here. The authors wish to acknowledge Dr. J.J. Emerson for his insightful early review of this manuscript, Dr. Bhagyashree Kaduskar for help in validating genes of interest and Dr. Sonia Sen for enthusiastically initiating work on olfactory receptors, Dr. Pushpinder Singh Bawa for library preparation of individual samples and Dr. Sushant K. Ghosh, Senior Scientist (retd) of NIMR, Bangalore for providing the mosquitoes.

## Author Contributions

AC: Validation by synteny and resolving scaffold conflicts; SR: Developing homology-based assembly method and preparation of the manuscript; SJ: Assembling and stitching chromosomes; KP: Annotation and genes of interest; JS: SNP analysis; CS: Olfactory receptors; SW: Transcriptome and Browser/database (website); NK: DNA extraction from individuals and quantification; RRK: Sequencing; CG: For maintaining lab strains; BC: Overseeing lab work including library preparation and sequencing; SS2 Subramani: For initiating the work, interpretation of results and scientific editing of the manuscript; SS1: Came up with the concept, managed the team and wrote the manuscript

## Competing Interests

The author(s) declare no competing interests.

## Supplementary Information

SupplementaryMaterial.pdf - Supplementary Figures, Tables and Text referred to in this manuscript have been compiled into this file.

## Data Availability

The improved IndV3s and PakV3s assemblies for India and SDA-500 are available on NCBI BioProject under the project ID PRJNA473393. Genome and gene annotation files reported here can be found at http://3.93.125.130/indv3s. Scripts and transcriptome related data can be found at https://github.com/SaurabhWhadgar/indv3stranscriptome.

## Notes

http://3.93.125.130/indv3s

https://github.com/SaurabhWhadgar/indv3stranscriptome

### Competing Interest Statement

The authors have declared no competing interest.

