## SupplementaryMaterial for "A near-chromosome level genome assembly of *Anopheles stephensi*"

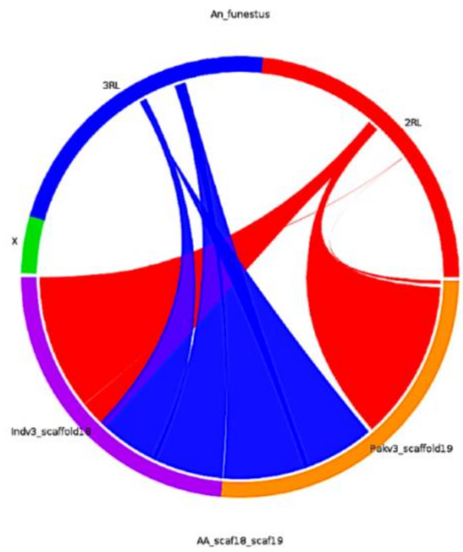


| *An. stephensi* | *An. gambiae* | *An. funestus* |
|---|---|---|
| X | X | X |
| 2R | 2R | 2R |
| 3R | 3R | 2L |
| 3L | 2L | 3R |
| 2L | 3L | 3L |

**Figure S1:** The matrix shows syntenic arms from all three *Anopheles* species. The figure shows synteny of scaffold18 with syntenic arms of *An. funestus*.

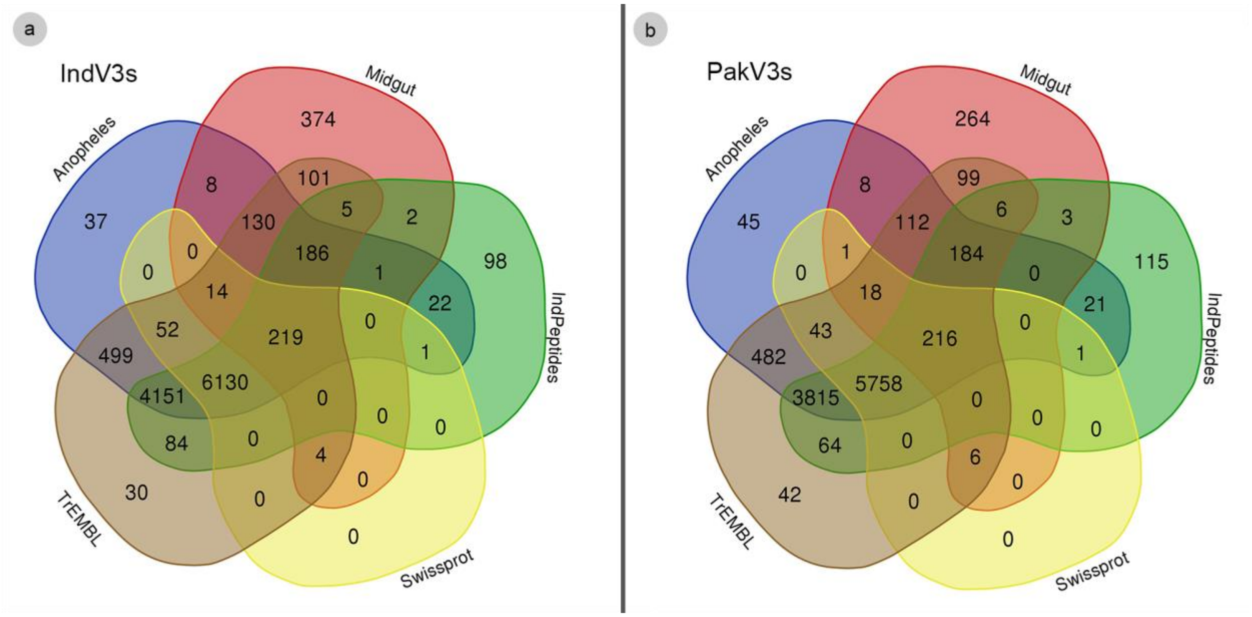


**Figure S2:** Venn diagrams depicting the intersection of proteins validated by 18 other *Anopheles* proteomes, *An. stephensi* female midgut transcriptome, *An. stephensi* Indian peptides gene set, Swiss-Prot and UniProt TrEMBL databases for IndV3s (a) and PakV3s (b) respectively.

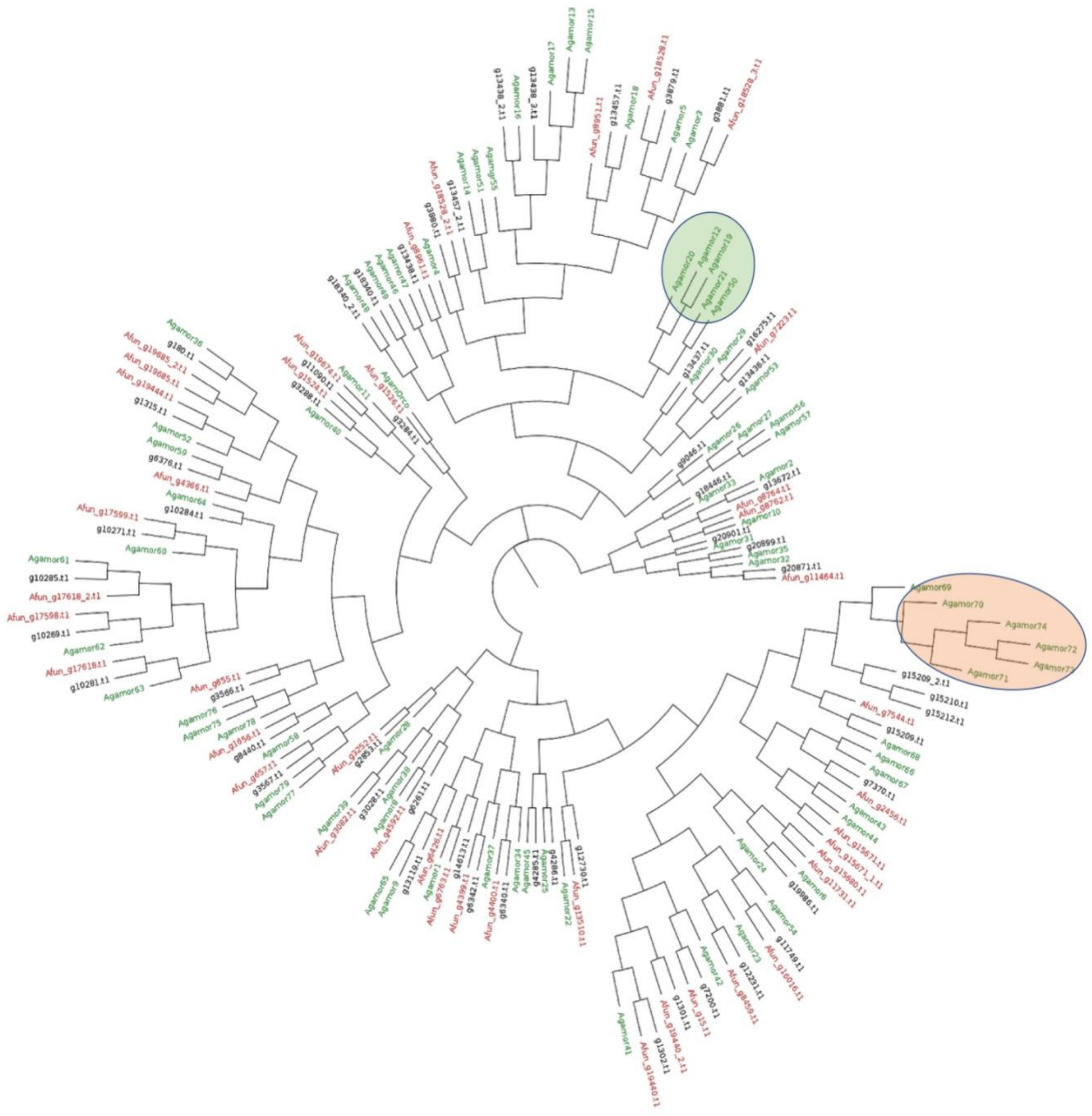

**Figure S3:** Phylogenetic tree of all olfactory receptors (ORs) from the three anopheles species including *An. stephensi* (black), *An. funestus* (red) and *An. gambiae* (green). The yellow highlights the branch that is unique to An. gambiae and missing in the other two species.

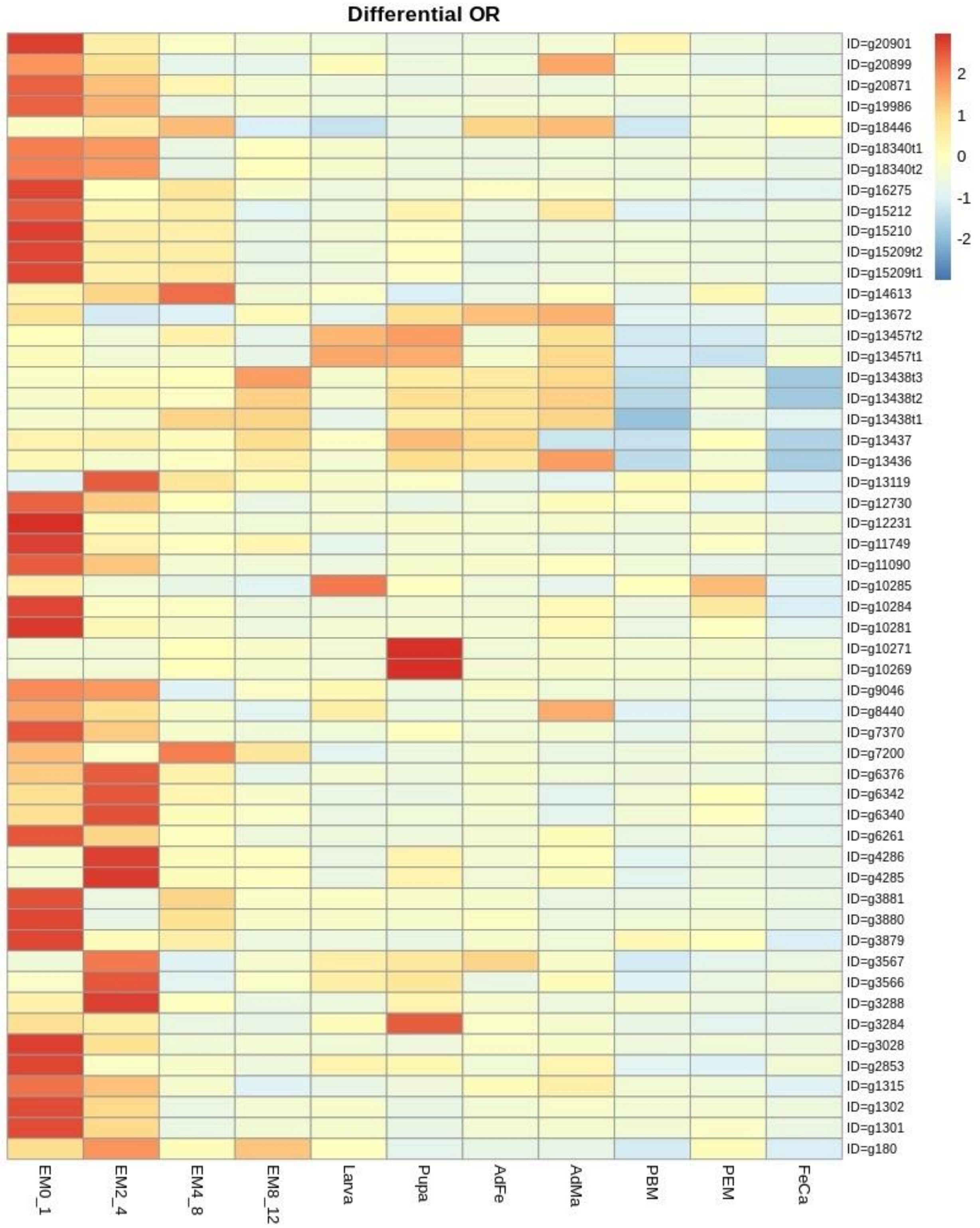


**Figure S4:** Heatmap of gene expression profile of all 54 ORs across developmental stages: EM0-1: embryonic stage 0-1 hours, EM2-4: embryonic stage 2-4 hours, EM4-8: embryonic stage 4-8 hours, EM8-12: embryonic stage 8-12 hours, AdFe: adult female, AdMa: adult male, PBM: post blood meal ovary, PEM: pre emerging ovary, FeCa: female carcasses.

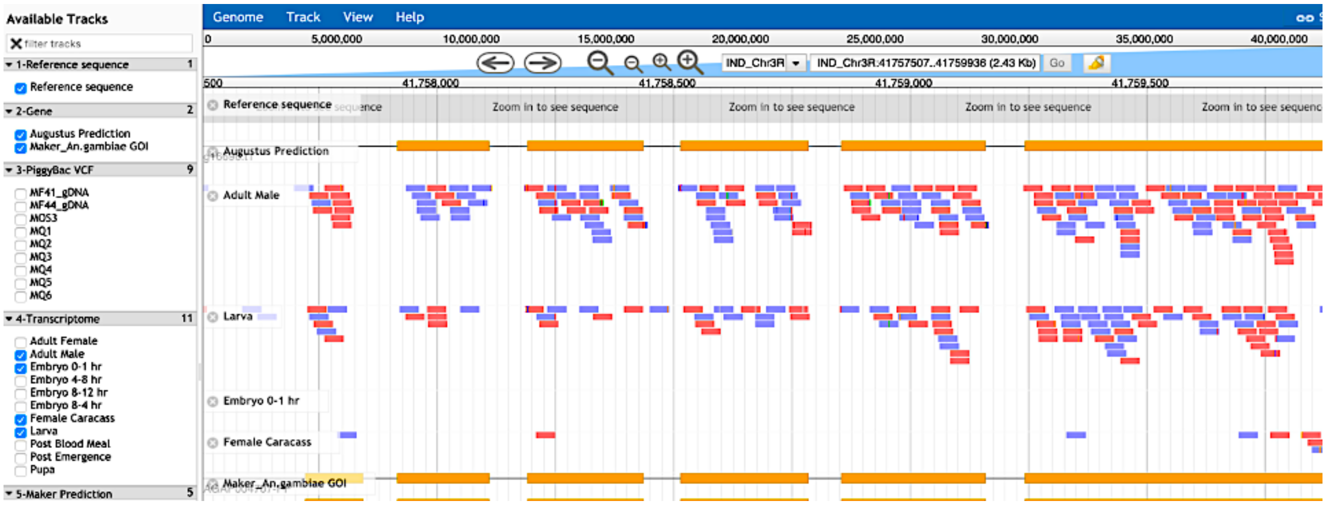


**Figure S5:** Expression of *KDR* gene in adult male stage.

| Gene of interest | Source/Database | Accession ID | Species |
|---|---|---|---|
| SPO11 | UniProt | A0A182YKK2 | *Anopheles stephensi* |
| Cardinal | UniProt | Q9VCW2 | *Drosophila melanogaster* |
| ACE1 | UniProt | Q869C3 | *Anopheles gambiae* |
| CYP6P9a/b | UniProt | B5AII6 | *Anopheles funestus* |
| CYP6P4 | UniProt | Q7QCY8 | *Anopheles gambiae* |
| RAD51 | UniProt | F5HKE1 | *Anopheles gambiae* |
| MRE11 | UniProt | Q7QID8 | *Anopheles gambiae* |
| KH | UniProt | Q7Q6A7 | *Anopheles gambiae* |
| RAB5 | UniProt | A1YSB1 | *Anopheles gambiae* |
| CYP6M10 | UniProt | Q16WQ8 | *Aedes aegypti* |
| KDR | UniProt | Q0N3S3 | *Anopheles funestus (fragment)* |
| FREP1 | Niu et al 2017[13] | ----- | *Anopheles stephensi* |

**Table S1:** Online source of data for genes of interest identified in *An. stephensi*

| S. No | Samples | Accession ID | Number of reads | Read length (bp) | Sequenced bases (Mb) |
|---|---|---|---|---|---|
| 1 (EM4-8) | Embryo 4 to 8 hours | SRR514863 | 8335955 | 39 | 323 |
| 2 (EM8-12) | Embryo 8 to 12 hours | SRR515304 | 14390100 | 38 | 532 |
| 3 (larva) | Larva | SRR515305 | 14725066 | 41 | 574 |
| 4 (pupa) | Pupa | SRR515306 | 12192512 | 41 | 500 |
| 5 (AdFe) | Adult Female | SRR515307 | 14263361 | 41 | 586 |
| 6 (AdMa) | Adult Male | SRR515308 | 16487004 | 41 | 676 |
| 7 (PBM) | Ovary post blood meal | SRR515309 | 16466546 | 41 | 676 |
| 8 (PEM) | Ovary post emergence | SRR515310 | 14422518 | 39 | 567 |

| | | | | | |
|---|---|---|---|---|---|
| 9 (FeCa) | Female carcass | SRR515315 | 11175963 | 41 | 457 |
| 10 (EM0-1) | Embryo 0 to 1 hour | SRR515316 | 5524113 | 41 | 225 |
| 11 (EM2-4)) | Embryo 2 to 4 hours | SRR515341 | 5210129 | 41 | 213 |

**Table S2:** Transcriptome data summarizing sequencing metrics across the various developmental stages including embryonic stages 0-1 hour, 2-4 hours, 4-8 hours and 8-12 hours, larva, pupa, adult female, adult male, PBM (ovary Post Blood Meal), PEM (ovary post emergence), FeCa (female carcass).

| S. No | Samples | Accession ID | Mapping percentage to IndV3s |
|---|---|---|---|
| 1 (EM4-8) | Embryo 4 to 8 hours | SRR514863 | 84.11% |
| 2 (EM8-12) | Embryo 8 to 12 hours | SRR515304 | 89.63% |
| 3 (larva) | Larva | SRR515305 | 87.35% |
| 4 (pupa) | Pupa | SRR515306 | 82.44% |
| 5 (AdFe) | Adult Female | SRR515307 | 87.46% |
| 6 (AdMa) | Adult Male | SRR515308 | 80.10% |
| 7 (PBM) | Ovary post blood meal | SRR515309 | 89.87% |
| 8 (PEM) | Ovary post emergence | SRR515310 | 91.15% |
| 9 (FeCa) | Female carcass | SRR515315 | 82.60% |
| 10 (EM0-1) | Embryo 0 to 1 hour | SRR515316 | 84.56% |
| 11 (EM2-4) | Embryo 2 to 4 hours | SRR515341 | 85.23% |

**Table S3:** Percentage mapping of the 11 transcriptome developmental stage samples on IndV3s reference genome.

R version 3.6.3 (2020-02-29)
Platform: x86_64-pc-linux-gnu (64-bit)
Running: Ubuntu 18.04.4 LTS

Attached base packages:
stats, graphics, grDevices utils, datasets, methods, base

Other attached packages along with the version
edgeR_3.28.1, limma_3.42.2, reshape2_1.4.3, tidyr_1.0.2, dplyr_0.8.4, ggplot2_3.3.0 knitr_1.28, corrplot_0.84, pheatmap_1.0.12

Packages used for Rmarkdown along with the version
Compiler_3.6.3, htmltools_0.4.0, tools_3.6.3, yaml_2.2.1, Rcpp_1.0.4.6, rmarkdown_2.1

knitr_1.28, xfun_0.12, digest_0.6.25, packrat_0.5.0, rlang_0.4.5, evaluate_0.14

edgeR is a Bioconductor software package for examining differential expression of count data. Here we have used inbuilt RPKM function of edgeR, package [40]

Reshape package was used to Convert an object into a molten data frame which was further used to plot using ggplot package

ggplot is used to create visualization for given data such as a heatmap, line chart etc.

Pheatmap was used to create heatmap for the transcriptome data. The heatmap given in supplementary material generally top 100 as the choice range of the data.

Tidyr and dplyr are the data wrangling packages used for data handling and to filter the data based on the condition.

Corrplot is used for plotting the correlation plot between the sample based on read counts.

Knitr is used along with the htmltools, rmarkdown to create HTML output for RCode

**Table S4:** Description of R tools and packages used in transcriptome analyses

| S.No | Sample | Mapping percentage to IndV3s |
|---|---|---|
| 1 | MQ1 | 73.81% |
| 2 | MQ2 | 74.58% |
| 3 | MQ3 | 68.57% |
| 4 | MQ4 | 75.77% |
| 5 | MQ5 | 73.00% |
| 6 | MQ6 | 78.40% |
| 7 | mos3 | 68.56% |

**Table S5:** Percentage mapping of the 7 individuals from whole genome sequencing on IndV3s reference genome.

| Assembly | ChromosomeID | ScaffoldID | Size (bp) | Assignment |
|---|---|---|---|---|
| IndV3 | 3L or 3R | Scaffold18 | 38,83,490 | 3R |
| IndV3 | 2L or 2R | Scaffold12 | 48,18,062 | 2R |
| IndV3 | 2L or 2R | Scaffold2 | 1,81,36,279 | 2R |
| PakV3 | 3L or X | Scaffold3 | 1,06,18,294 | 3L |

| | | | | |
|---|---|---|---|---|
| PakV3 | 3L or 3R | Scaffold19 | 42,66,331 | 3L and 3R |
| PakV3 | 2R or 3R | Scaffold8 | 87,12,747 | 3R |
| PakV3 | 2L or 3L | Scaffold10 | 80,43,466 | 3L |
| PakV3 | 2L or 2R | Scaffold12 | 51,10,278 | 2R |
| PakV3 | 2L or 2R | Scaffold1 | 1,58,21,620 | 2R |

**Table S6:** Scaffolds of IndV3 and PakV3 which show the presence of physical markers from both the chromosomal arms present in column 'ChromosomeID' required resolution and have been designated a chromosome in column 'Assignment'.

>g180.t1
MQLNILLQRCRRLLERLYIERDFFRPYEILLALPGFHLVDGFRRKTWVRVLFFVSRIVLLL
QYAIWADRCYLGLINAPQHSGKALHYGNTFGVLTMMLVRMLVVRWYLPNVEQLMRY
LRGQQRHRKSRSTTHRVSYRKIANIAVTFQLIGLADRVVFCFSRTYREELYQMPSNLVDL
GWPMVVALHVFSFDFASRWAAAFNVSLTGTNSIMMGLYDELVDIADDYSRMFEVRGS
DGEFWTGLERHIAQTVKRHESFIRELNQLKPFLQITFLVMFYSAALFLAIGMFMITANGT
TTYDVILSGFLFALLLECYWCCRLVDRLNDMNTQIGKHLYNLAWPTELQYTWADRSRY
RQARSSLLIMMSSTQKTLGISCGGMFEMSSEAFASLVKMTYTVLTFLRDTQNFS
>g1301.t1
MAAVKESPIDKFNRILSWQLHILRMLGLDAFSCRLVLNPLALTIFLMAGLFMVVSFYDV
LVLFRGDLFGTSFVLTTIFYGFIGWARILGALAYRSKLPLLMQMTRDTYHRAVRDKRQS
AILARYTGIFWRGVMLYSLMFLVGVVIASVGPALLFLYSGKKILPFGVYLPFVDPNSGTG
YELNYLYQMSCILWTPPGLTATQNIYFAFILNICIQYDVLQLQLADLNQLIQWSGVENQD
NAVRKKLREIIVYQRRLEVFVNTIEQVYKMQALVEVLSLTFQLVLTLYVMRTSMWPPGL
ILIPLCTVQLFILCVPGTLIEIKASHLTETIYGIDWHDMHQKNKRIFQLLLHRSQHPRFLTC
ARMAIIDLNLFLSVRRLVGVEMRLGWVYYTDPFRSQVMKKVYSIFMMLENM
>g1302.t1
MVEHPIYPFDRLIKRQRLLLKLIGVDSFDRRYRFNKLTVMVIFLAGFFLVVSLYDLYLFR
HDVFNFVYVLITIFFATIGIGRITVFLWYSSTLSGLLSQTYHTYRLVKEDDERKWNILAW
YTLMFQRAVNAYTILFIGTSIATGILPLGIYLLSGERVLPYGVVLPFVDPSSQKGYELNYL
YQVSCIIWTPPGLVASECMMFALVLNICIQYDILAVQLLDLDQVIRSHDPDREALISQQLR
AILHGQQRLISYISSIEYSHTVVAGVEVLSVGLQIVITLFVMQFLFLFCLVGTIIEQKGEKFS
DGVYNLTFNELSREHKQIFRLLLLCSQQPKTLTCARMTRISLNLFVNVRTMGRIGSFNPRI
QE
>g1315.t1
MTLWSYLRRKLSPILDLRQDSDYFVLLKWLYAFNGIQLQTNRLWLRALYLLYRLLLPA
QCAIWLYRTWAAAYVERNKNLALSLLCGQFALTSIMFRFILLLLRHNQLQPVRSYINAR
RFLRDHTKAQGLRQRVFRTNNILIIGLMIYGMINFLIYEASDLQWHDIFRMPPYLMEMNR
PLAWTLQIIMHPMTLNGLGAYVTSFLSMHTLLTGLQAEFLLVEFAFVGLLKRVEEQVLQ
GAIEDDARQRLLWETFNRELGICVREHCEVVKHIREVHRVHSFSITVQYYTALLSLAIDT
FFISYNGLDFVSLSVLIFSVLLVFEWYYCCKLVEDLQATNKRIGWALYTDDWPAWLQH
GKRQPGALRQFRITMSIVLLVSQRSLSFHGSDIVEVSWETFGSMLKTSYSVMMFLIELRK
LNR
>g2853.t1

MPNEKGSSGLFVLRLGNGTEMVRLVLHEVRYVLIVMFYSTRCLTAKIQNSLVDKYIYW
FLTLIPIAMLCVPQFAYLLVDSKGLIDFVSVLVPFTEILLTNLKMIICNIKREKIIKLINEIQA
EWTEYQKSDQHEIQSLITSTAKKTRIFVIIYTASFVLICLEYASMPLFKFVYFSLFSENHAN
FIVTIPYSSSTDTSTSFSLTYFFVMIAVYMLALTLSGFDSLFATLAMHITTMFQFLKIEIDQ
LGSDMRAGTGRAELRDKMKRIILKHKTNLSLIEELEDGFSFYLMVQFLTSSFVVCVVFYE
LTIVFGWNEDTFKTLTYLPGAILQLYLFCWYAQNITEEARLVSDHIYNTPWYLCDLPLQK
TILTFMVKAQKPTGVTASKFYMVTLQSFQRISSTSYSYFTLLQTINQ
>g3028.t1
MVSTVAPVSDIPNASQTTKNWDIFKLQRKILLVFGLWPADRLVRRWYLKGLIAINLAAL
AICMVGEFLHGLYAYRDGNLSECIESICPTVARISGFLRMVFYLVNEAKIDQVLNNISKLL
QDKHPRDNAICKQMTTLGQQFTFYLFLMMFFAACLYGVTPFCIMAYNWYQGQRPLVK
LLPFKLALPFDSQNSYYFVLTTIFLNYASAPTITSQSGSDALFSGVCLYVYGQFQAIKLDL
EALSATLDKGSLKGSVAETQRVTDELRRISKRHQQIIDLVAEVRTAFTPNVLLAYTATAII
MCIVCVAMLVVEGIYKLTYLPYAFAELTLLFLYSYSGTIIRDSSETLQTVAYDFPWYRFD
RNTRHLIQMIMIRAQHGSNVDVPFFETSMASFSAIVRTASSYITLMKSFL
>g3284.t1
MVHAGKHRVEPASAITSTSMQVQPTKYVGLVADLMPNIRLMQASGHFLFRYVTGPILIR
KVYSWWTLIMVLMQFFAILGNLASNADDVNELTANTITTLFFTHSVTKFIYFAVNSENF
YRTLGIWNQTNSHPLFAESDARYHSIALAKMRKLVVLVMVTTILSVVAWVTITFFGESV
KNVLDKETNETYTVEIPRLPIKSWYPWNAMSGPAYIFSFIYQIYFLLFSMVQSNLADVMF
CSWLLLACEQLQHLKGIMRPLMELSASLDTYRPNSAALFRAISAGSKSELIINEEKDPDV
KDFDLSGIYSSKADWGAQFRAPSTLQTFDENGRNGNPNGLTRKQEMMVRSAIKYWVER
HKHVVRLVSAIGDTYGPALLLHMLTSTIKLTLLAYQATKIDGVNVYGLTVIGYLCYALA
QVFLFCIFGNRLIEESSSVMEAAYSCHWYDGSEEAKTFVQIVCQQCQKAMTISGAKFFTV
SLDLFASEPTVAYGHRRSYETHGNMG
>g3288.t1
MWRSSSEDDQTLSLNFRMLERILRFVAVWPTDYNPYLPKYLRGRYFLSELIDTCYLLFW
FFICVHIAAFHIVSIVVMDLSYDELFLTLITTSIYSIMTLLSLYLRLYESNVRQLYEFTIRHF
RKRSAAGVHYISISSSIRVTNRYQFWWLIICVLGTMHWAIYPILSQERTLPFPCWYPVDV
QQSPMYELAYIFQVLGQLQVSLVYGLAGALFMVFVFMTCSQFDMLCCSLTNIRQSAMIL
NGHYRQELRHHQDNHELDTREYVLKEIFREDLDNVQPTKTASKLDQLSPSQSYLMELSP
ELTCVLEDCIKHHLLLLRFCQLLESCYHPYILLKLFQILLLLCFLSFMATVESLSTMKLINV
LEYFMLTMTELYLYCFLGQILMNQGFKVGDALWKSPWHLCGASYRRRMLIILMNAQRP
VRLTGLKLYELNLETYYTPPTLSITNLNQKLSPQNANELQRNLPNKPNPRQVMRGSRDR
ASDIDGPFIQNPVGPEVHRNLKPFAVPLLHTLSPKIDRVSGGFASLRREQRDNGKARERA
RNNCEKLLLALRSRFAKQNSAPPKKDGRRENKTQRNRERERSREKE
>g3566.t1
MVVKRALLYSFSLLQHHFNVGHPLEHFCLLRCLDVVSPAMLIQRPRSGLEIGIKTLSLSV
LLTHVVGLAYDLTQQEDIRVAMDIFCMLSLFSSLFARNTCLRQYQSHIIAMERLDANPGF
EVGVPYAETIRHRTVTQNNRYLGWYLVSHFLTVTIYVTQNMAMQGSFVKIITHFPIDLSG
YAPALDTMTQFFYTVAGYGWAWYHAAGQLIVIVLLRFVIAEFRVFLHSLATLDEQIHDR
LLLELDGNEERVVRELLYKHARQHSQLIVVVMHLRAILRIYSLVHFSFYMIIMAAFMAR
VLVIPGSSSLGMAIPVLVTIVFFLETFGLCMLVEKLVQLNRRVSVNLYGFGWTRYLQYG
HSIKRTMMLMIMQANNTKDFSAGGLTTVSAELFAKTCRLVYTMMMGMANLAT
>g3567.t1

MSSSGQGSNICFLGWIRWMDTVNGIHLHDESRLARAFQLVFYVLQLSQLLVIYNFLSSCL
HTVSLEEFARQFNQCGGYVLTFFRVATINMYRTDLKETAQFINAAEFHHLNARAEKIRS
GPIRHAGRVLSILFAIQVVAITLWFIMTELQAQAQNVLLPTVTYLPFDASGWPTLLKVMF
RLYVYLSCTQLFLTFFGSYIITSSYLLTLTIELRILNDSYAGAPDDPQQLVAFLEDRVRYK
VTLLHHIGIIKRQMNVGLLFELVLIVCLLAINGLRVCTTSSDLSEIALSGSMIMIYLLEFFQ
YCWQVDEMQQLHEGQAFAVYSTPWVGAMHETKAQLLITTRMAQVPLRFMCGGMYQL
STELFATVVQFIYSLVMMLLHFK
>g3879.t1
MVLPKLADPFAVMPLLLRLQRFVGLWGERRFRYKFRLAFLSFCVLVVIPKVAFGYPDLE
TTVRGTAELIFEWNVLFGVLLFSLKLDDYDDLVYRYMDIAKIAFHKDIPAQLGDYLVHI
NHRIDKFSKIYCCSHLCLAIFYWVAPSSSTYMAYLSPRNKSRPVEHVLHLEEELYWFHTR
VSLVDYSIFTAIMLPTIFMLAYFGGLKLLTIFSNVKYCSATLRLVAMRIQLMNRLDEVQA
EKELVEIIVMHQKALKCVELLEIIFRWVFLGQFIQCVMIWCSLVLYVAVTGLSTKAANV
GVLFILLTVETYGFCYFGSDLTSESLSVARAAYDCYWYQRSVSIQRKLRMVLQRAQKPV
GISAGKFCFVDIEQFGNMAKTSYSFYIVLKDQF
>g3880.t1
MGFVLQLLHLVGLDGPGRASRIRLASVMLFYLTFIVIPELTGGYTDVHQFVRTGVELLFN
CNIFVGGMLFALEVTSFRMFIRELKFLAMLASSLSYKLKHALARFNYRANTFAKLQTIC
MGVIALAYWVAPLPSIYWFYYHPDNATEPPVRLVQHLEVKFYWLENRTMLKDYVAFA
VIMLPVVCMCSAVCNIKVMTISGSIAYCTLFTRLTANAIEQLPDVAPAGWNPKALSHVV
SMHASLLKTIHLLDRALRSVLLLQWLGCGLNWSISLVYLTNTGISFKSSTVCVMFLLATS
ETFLYCWLGSRLATQQERLERAIYAKRWYNYPRNERCSVLTILRQAQKPAVITVGKFFR
VNLEEFSRIVNLSYSAYVVLKDQIKMDAI
>g3881.t1
MVLPKLKDEKAVMPFLLRIQTIAGLWGDRSQRYRFYLIFSYFCLMVVLPKVLFGYPDLEI
AVRGTAELMFESNAFFGMLMFSFQRDNYEKLVHQLQDLAALVLQDLPAELGQYLIAVN
RRIDRSSKIYCCCHFSMATFFWFMPVWSTYSAYRAAATNSTEPVEHVLHLEEELYFLHIR
TSLVHYTFYAAIMWPTIYTLGFTGGTKLLTIFSNVKYCSAMLKLVALRMQCLTGVKRER
VEEELNEIISMHQRALDCVLLLETTFRWVFFVQFIQCTMIWCSLILYIAVTGFSSTVANVC
VQIILVTVETYGYCYFGTDLTTESYGVALAVYDSDWYKFSVSMRRKLRLLIQRSQKPLG
VTAGKFRFVNVAQFGKMLKMSYSFYVVLKEQF
>g4285.t1
MKKFSTSEANLDRMFALIAKHMEVLKLNIFKPEWRLSLRTVLVLVAIGFMPILTAFSVN
KYYEHLEIKVECFTQACTGAQVFIRSYFYLRQRDQCRQLATEIRRQRISYGVNENERME
QLFRRATERMLMLYRLMYAMYCGSFFFVLGPLIMPDSRKASLPLAFRIPYLPPDENLLY
WCLNYVHHIFLIVVGIHHLAPIDGIIVLALISICTRISALELLLNELDGKITESKWQQTEHLE
PCLDRIIELHTDMKRFAELVSSTFEMHFFTIFSMICSIICMCLNVIAGQPRNSIYPLLLASVC
QLFVVCLFGNVLLIVNDRLPKSIYGIQWYRLTVTQQKKILFLLANAQTDIVMSAVFKPVN
MTSFVAVGEKVITSDEDYTYYSYFITGLSGILFVLHHTSLKRMHADEIYIDPNR
>g4286.t1
MAHKHRKEMDQELDRIITFLRRPLQLLGLDVLDPSWKLTPRTVFTIGMFFLQHYASAWF
LTTHLDAFVVFTECFSTSAVGLEIAIRMGLLLYHRELLNETVQIIRNQKRSESFQKLANQF
NKVVSVSVHLIAVMYLSTMMFELIPIVYPDPKKSNLPLALFIPHMPHDVAPYWQINYVY
HTVMNLICVMFLFAVDGTLVLSILAAVYQIKGLKLCLQELDTGAEQSVLQRELVRICKT
HQSIKQFIRLLDQTYYLDLLVDFGLVCLILCMGLNVIAVDVMQPIGVYLIAVAFQLFLLC

FCGNLLLIESDSLSNVAYSIDWHVMPVPEQKLLMFMIAHSQKPQKLSGIFMPLIMSSFMS
VIKASYSYFTLLH
>g6261.t1
MLNLLSVTVGKTPAREELFAKANRVQRLRWEREPSIMPPSPKPSTIGKNNGMSRSPSNGI
ARSCKKKMPPANSTDQLVQFESFIRVPEIFFTMIGVARYGEPRRTLQAHLKQLLFWSSCA
NTGFCLIIEHIYFVKAAGNFTNFLQLTALAPCMGFTALSFVKIMTIQLNGTKLTDMLHRL
EALFPKSAALQERYGVFQYNRESEVVMKSFSILYMTLIWMFNLLPLVSMVAGYCADGT
WHKQLPYFMWYWYDWHEPGYFEVTFLHQNWGGFVSAVFYLSTDLMFCAIVLLVCLQ
FDIVAYRLKHARPDDQQELHECVRIHQAVIELCSELEHMFSPSLLVNFLSSSVIICLVGFQ
ATAGITPADLFKFVLFLVSSLVQVFLLCYYGNKLIVAVRDSGSISSQIPYSAFEGQWIGAS
VAYQRSLLFVMLRSTTVQKLTALKFSIVSLASYSKILSTSFSYFTLLKAIYEPNEKNVK
>g6340.t1
MAAFGTGSHLLSSEYRMPFFPWFFGIPYGDEARVAYYTIFVYQCFGMYFHMLLNTAGD
TQLCYMLQMIGIQLDLLAKRFRSLSNSEEFDRSFVPLVQHYNKIHRVENLFSLAYFVQFS
VSGLVICASAYQVASMFNLYDFSKLMNVFYMMSMTMQIGLPCYYGNEVTLKSYALTN
AIYSSNWYSMKQSNRKSVQMFLVRTNKPFAATAFRYFNFNLPAFTTILNMAYSVYCVL
QRKAKNV
>g6342.t1
MKIETQKTWSTVDKYRWSEYIRPVRMTVWTWRICGLYNAKPQTPLYRAYRIVFNVVL
MVVYLFTLSFNMFVMQTFEQLVLYIMYIVFTEIVMVLKALITYYKFDQICYLYRQTLDS
DFKPVDAEEEQLHRKGIGEINYYFYLYITTTHLAIASSLLYLLHQDYRMPYFPWVMGIAY
GSTERLNFGIMFAYQVVGMYFHMLINVAIDVQLCYFLGMIGIQLDVLGKRFRMIQTSEQ
FGNSFMTLINQYQKLHK
>g6376.t1
MEWIRQQLNIVWHLHEGRDYLHCLRPFQLVAGYPINLRPVLAKLVTGVRIVVYLLYLAS
LIHKICYVLYRPEDINYVSFVSGGITVLVAVLLLMIIFTIHYDAFVQLGDFLNDRSFARDH
PQAARIRERWYRWSNCLILGPQCGIILILMQTWFSRQHRKKHTMLVIRGEPIGTEFDQLL
YVSFLYFPTVGFFMGCSIVNAILVGFMGEMELLATCLGDVFETVEKQPTVQKAADNRST
YWITLHEQLRQCAKRYCEIFTMLPKLQRMASFVFLQHHIFSLGLVTAGCYVTLRGPALR
ENVVLSEYPISVVLEYFIFCQLVERLQDMYARFDFIHSPSIQNANAAAYVMSLGLYLPSL
VEARFRLCATITTAGQVLSTVTAKRSNLGRCWYVCSSLTSGEVCSGVPIYEDKHNDLGL
GRLVISPAGLSIRARVANRWVRRTSENIKM
>g7200.t1
METSIEKHHSLERFRATLAWQNKILALFGCYVYVRERRVTSRIVAICFIAISFIVLSVYSA
VQSWGDMGQVLLSIVAVFYAIVGVARLAVAISDPAGCYHSIKLAEEMYQHANGSHRAE
CTVLAKYTDLFCKSVHLYTFGFMLSVVLVSVMPFAFYLFRGERFLPLGIVFPFTDGENM
YGFWSTLAVQLAYILSGPLALVPSQNIYFAFVFNICLQYELLIERLKQLDETIRSSGSIEQG
PKRSTVRDQLVKIIQLQQRSTNYITHIENFYQMQSFVEFLCNSLQAALTLNELHRNFWLP
GFFILPMAVGQMLILCSLGTLIELKSDQFKDQLYDIAWSEMELPQQAMFKYVLQSAQQP
MRLTCGRFTVINMNLFLTVGIVGVLFLEE
>g7370.t1
MEAAEKFHQYERYLRTLCNVLGFDVLRKGWKKTFRTYVTIFLCGQYFLWMVWSIIIAS
DTFELLKSLSFIGFFFQCSSKMYYTIANAAHYSTNFAGLQETIYTAHMDGTEEQKTVIDR
VITVVLLATKATTVLFTSSLFIFSLYPAYMYFVMNVKVTIFPLYIPGINIYSSYGYGITNSL
HMLIAVYGLLGALTSDTAFMLFVLHFISYVELFRIECEQFARDLDAFGQQWEYHTVEYK
TFCRDWMRALYQYHQQVIVYLSSLQECYHSICVYQVASCSFSIMFNLFLALTTDWYAT

YSFMVISWFQLFVYSLLGTVMQVMNDRLNSYISNLPWYLLPTDEQLRYNFMLGRSQLP
AEMVIRSVGPMNMETFTDIMQKIYSAFTMMYSFLVDLG
>g8440.t1
MSSLVRLAKQYTQRVTDAGQLVIVNRMDRFIGFFSWDVQQRFTWLKIALIVFAVTYETT
AIAAMALASIKGVFTERSFTMSFVTLTGAMCIVMWISLAVFRRDLTTTVAFLQQRQSAIH
KHNAPRKALLDRVTRYLWLFYLQNIAQVFFWINLLRDCSPLAVFELSLLDSANVLLYPIA
MTLMSLMFIHTIMIVSTLLSGLTLEFYWLGQEFEQVFAECSSISVTWHQRYWDALEQRIG
MCVMEHQRLLGQISKLRNNLKLYLLLNLVADFSLITFAGCQMVISSEGDQHLYSILAALT
ACLNMLNFGGLCDLLKIQVHAIKFHLYSSQWTDYLRPVSGPLYQRCRRIRSSILIVMTRA
EHELRISCGSVYDMSLAT
>g9046.t1
MELSPGKGKFPLVDLTIRGLKVMRFWNEKPAQTFSIFGLLLVVVYPIVWLIPSWLFIISSQ
DNLTLLMKAANEQIVFMAIFFKLCSFVINFHRWEQLFYDLQRAFTSVMDDQSLDIQGILG
HVEKTAHFLTKGYTSVLCFNCALYGVFPMLFVAVKYAITGSHDVPLSTPIEANYFIPGYR
THFWLWLPLNVMLNVLLEMHGIALFLIECFTWSLVHATSCLFRVLQIQAHELSNQNERK
DQWYAKFESFVSLHESVLRSARTLEEILSFQMLFLYMSTVFALCLMVMVLSLAFNDVFL
LIAMICVIGYCLFQTFSFSYLGTELIEESGAVADAIFHSTWYNQSVNRQKDLCFVLMRAK
KPVKLTAGKLFIVTRDSFTEVIKQAYTIFTLMSQFLEESAN
>g10269.t1
MAKRAMVPERIPGKPLVRKYWDKFFTFTSTVDYFNLLNTFGTVFALHYHSPNTRWTWK
KLLWMVYRTFYLLSYLSYCYKTYWTFSNWEYSTASANVLGALGLCSGALLRLVLVEL
NYPTIRKLQAFLNDRTYLNEDQWAQDQRSQLYRHNNRFLVVLISAITVESLCFLARLLLT
RPEFMFQYNGRVLGGPAVQIVYGMVTACWGIIYVLSFIGFYMLLAVFRLEMELLARSFQ
QLEETLLPDQERMDTMDQDQTERAYWNKLQAELTIRIKRHVELLEYCVPSAASPFAFLQ
YYCTFGLIADSFFVVSFEGFTGYSMAYVLFASFLILESLLLCRGVEDLNDLSSSVDRGAG
LYTAGLDLRGLALPFLAFCNFPAQAYREAVWIEFELRSCSVIGAVIALAAGSPHAGLAIK
QSVRVSDCKWYCAMVPAMAHCWGSPVLVLLARGLSSGSIIITLNNGRFPMHSHVPSVV
VVMLLAAFCIEKRKGIIIS
>g10271.t1
MELFLSQIYRKFHDSELRFANNPDQFVILRYLTFLYAIRCDSPLAMWQRVLWYCHRSGL
MLVFSSYCLKAYWHMNLGAYTFPMFNIIGTIWIFGGALVRRMLFDRSVLVRLERFLNDR
SFRGDEPAATIARRTVQRQNTRYLVGTALTLLLETFLFSGTNLMLQPEFMLTYRGRVVG
GVVVQILYGFATCYWGSLYVLIFSFIYVILNAFRVEMSILVQSFEQINQILHQHCPSVSAS
ETSIAEERKLWKDLRNLLKKNVQRHVELLENLIVFRSILGPFSFVQYYGSFVLIAYYCFII
MYKGITSLTVVYIAFIVFLVVESFLFCHIISNINELNAKIGMVLYAMEWYNKLHFSKRFAS
DYRHVRSSLLTIAIRTQSPLSFTINGLGTISRGRFVDLLNSSYSFMALMLQLKNEIAS
>g10281.t1
MNFMWLQSFSDRISHYSESADFFIIQRYFEKIYAIHYSARSWRDRTLWYLYRALYSLIYL
SYIYKTHWVLHHWQDSLSSANILGVMWFFSAVILRVAILEWHYPLMQRLQTFLNDHSY
QCSHPAIVTKRAQFYRRTNRLVLAVMAINFVEIVCFTATNVMKLKDFMLQYRGAIAGG
WPVQVVYGVLTMFWGGTYCMGFMVCYLLMCIFQLEIDILIQSLEDVGRSLRSGRESEAF
WDNIIDRLRPHIYRLEDLSITYLVIADCCFIVVSHGLSSYSIVYFISMMVFLTESFFLCHSVE
NLRNLKPRVASVLYDFDWMLQMQCSHDPNLSSHYRHVKRTFLLIIAQSDQPIHFSFAGI
GEISMNSFAQLLEKSYSMLTFLLQFAK
>g10284.t1

MSYIDRWASFVPVWFGSTFEFTKDADYFVLIQPLLKWLHLFANPVPYLGRMLSVRGVL
VQLYHFLVLLSYTSFVYRVYWQLFHPSYVAQLIIMVGAVLLYTLAIVRIWTLNKFGREL
QELRLFFKDRTYGEHCGWAHQNRAAVYRRWNWTIITLMLGSIINHFVFIGTNWHNPEFR
LQFRGVTVTPTIVRTIIEFCSCYIALGLFVSSSLIHVTLELFQTELKILVHSFKQALDTEQQR
ELEAHNAHIAYKLFCEQFYANIRRHTHLLQMFSKFARMLNLFGMFVYYGTLVIMTCTCF
FVMHHQFSSTVVSFVVFAVCLMIDTLLFCKRIDNINELHNSVGDIVYSGYWPALLRLAD
RGLSKNDQRSFRRSILIVLQRCQQPLGLGYGEFGSLSMHRFGELVQSIYSLITFLAQFD
>g10285.t1
MCHQQKRNVVQLLFRRYITFTDQSDYFALYRTLATISAIHYDAHCWFDRTLWIVYRFLP
ILVNVSYFYKAYRIIIHPEDNTSAAVIVASIWGFTEGTLRIAIIELWYDKLCSIMSFLNDRT
YRQQDALVRQQRAALFAANNRIQLVLVTTMLTIAAWFMTTQLFNRDAFMLQINGHVV
ESASVQIVYGLLCNVWGMIYVLSFAIFYIIMNILQLEMSILLDGIASVQSTVIDRAGRQIAT
LEATGHSTQMKQQVFWDILQPELNRHISRHVNLLDNLKEFSTIVGPFSFVQYYGTFALIA
DCGLILAMEGLSTNGMIYLIFVTVLVFQSFIICRGIEKINDLNEAIGHALYAGFNWPELLQ
YNVRFRAQYVTARHTLMLVIGRSQKGFQCSYGGLGGISMERFAQL
>g11090.t1
MELKEEWILPDAVYESPLLKRTLLGLRYYGLLLGQSQPLKKAHCLRGMVFTVSMILFNC
TQYVDLWQVWGSVSDMTANAATTLLFTTTIFRIIFFYLHRASKSLCIERLTFSFLVLGFN
DIIKVAHTGIERILGDGWDDEKDIVTSNVRYLSRLAVVFWSCALVTANMMCIYSLVLYL
MYDGPVDGQRNSTVPNSTSVLQQYPTPILRSWYPAADGKANHFLEIYLIQLYIMYVGQL
IVPSWHMFMVTLMIYGRTECSVLNHRLCFLDRYHKLGQNTPKPSGPDSVDNTERRTLII
DCIKRQANLVAFTRELEQLTRAAVFLDFVVFSVLLCALLFEASMTTSGVQVFIDVCYITT
MTTILFLYYWHANEIHACADQLSMSAYKSDWYRYDRGTNRMLQIFILYSNRPLKMQAF
FISMSLDT
>g11749.t1
MYLLERLRTVVRRRLERHSPDPREQYNSIVTSVNRIGGLVGIDVFTPNFKPGNMHLRLVL
LNSFAFFWINLYNLTTTYGNLVDFMFCFETLLYVFIAWIKMHVFVKHKSLILQLHQFMV
QFFDQFHGDPEQDALLVRTLVDTYLLVALFGFCSSAAAMLIFVSSLIWSVCVEYALPLGF
YIPTVGMDYLKGFALNFAFQLFESGLMVSGIISSETAFFIFLQNACLQVDMLRLELDRLG
RLGALNTDGRHTREIRSRIQSIIEHHIEHLDYSKSMCSLFELHFFIVFGCIFCQLISIVVVIVS
VPDWYPGYFLFIMLTAQLFFSCALGQVFNIKSDELTVAIYNVPWYNMEVCDQKAMKLL
LLASQHPGRLSYGFGTVNMRAFFEIYRKTYSIGMMMISVNEED
>g12231.t1
MKIWQRYLKRQRALFRTQYQSPKQLFDSACEMLIKCFAVCGGERMKPGYTRRNPRLIFL
VTDLILYLFVNLYSIAIVWGSLMDVVFCFVTLGIAIQGLAKIEAFTCPELNDLHLYNVARF
TMPPRFPEVEEALFHTAAMCKVFIRILAVAFSIVGIAIYSYAILMPLVEGELSLAFGFYLPF
IDYRTPIGFAINWVYQFIQVSEGCIGLMACDTCLLFLIVNATGQMDLIIIYLRRLTELIDSN
DAGQNDKKIADLLGDIVIKHLEHTKYVTDMDKLLKKQFFINFSCIIFELVASLAIVVRFP
WYPGMAICLICTIQLFVNCTLGTFLSSKNDKLVEEIYNVNWYGLSTKHQKTLQQVLLTS
QHPVVLSDGFSAIDLYNFVESWSPEADGDAGLPESTH
>g12730.t1
MSFFRKSHTQYLELSYSQIYKVFHWFLKLTLLRIFDDDFLVSPITTVLFQFFVSEIVISLGTI
AMHAVRYRTDLDTVILSVSAFVSALEVLLKLNGMVYRRKEIMQMIGTVLADRSYLRGPI
ERAICGKYMRLARKLLFITIASYLGTATMLLIYPTLPGVIQERTLPVGYSIPFVDYYKAPW
YMLNYVLQIVQLNWVAFLFIGLDGPFYLFVCYSASQLEMLIVPDNVPEQRRLIRKVYAI
HTNLSKFLSKCSYIYREIYLMQVLCSSVHICVSLFHIQLKLKNGSYGMLLTNVNKMWLF

CYCGELVVSKSTAFSTAVYTNQWYRLWNRRDLQDILFMLQNAQRNYGFSVGGFGFLSF
ALFTVGKCNKAENVDLNYLGRQVGKLSASLRQSLPKVRNLKDVIMSAWCVTYGGVET
KRKSI
>g13119.t1
MARIFFWKTKVEKFFTTREVQQIKLLPSLRMIFFIFGIFYAWPDERMEKSSLWWYRLKG
VLFRMFFIYLCTATQVAYNFTVTTREELFEGMFILLTQLVLILKMEFFYKNVFKIQKLIRR
LEGELYQPRTAEEDYPLERARKKTTTFWVLYFIFSDGLVTKWLIISCIYTIMLVPAWPVV
DHTAPYGVFLMVLGYQFVAMFLNASFNISWDSLVAALLALTNAHLHRLQIQLMKVGHP
QLRGSSDDKRLATVETIPDQKNSKDDVYNELLRCIVFHQEITGFLRDVLKLFSGPMLLQL
YCSVFILCITEFRLLTDVNTTTETIRALTYLICLIIQVVQYCYFGNEVNYMSQKVHQATAF
MNYPAMDIRTRKLLIAFQQLTAKGIHCSAKKIFTIELSMATFVTKIIGSDRFGRYSHAYTF
YAVVRKRVRSSKKSSSRFAPVSLGRCNECNNNNFPAFGKGAFARLTLALWACESIDKPI
LFRFPWS
>g13436.t1
MKFLRLDDAREIIPIGCRLLRLFGLGNNERFKLVYWIQIAIYLVFSLIPRFLLQLDDTVMV
LRFSSEIMFISYLCFQMVALYFRRAHLYQLVDMLKQCAGQPCSEDIQAFFIRSNVKINKS
SVSYVRFFLILYILYCTMAPIASIGVYMRNARNETSEKEEFIISSEMNLYYLDIRYNPLHYS
IYAASIFVLSGISSLSLCTKDVVDIAAIKTTTLMFQLSAMQIRELYGQFTQKQLNKAINMH
RDTLLCKKKLQDALNLSLLFQLACCSGIWCFMMFYILLMGLDSRILNLVVLLVIVSIETY
AYCALGTELTDSGEDVLMALQQLSWYDQSVTLQRQILFMIQRSQQPIVMTAGKLFSASV
QQFSEIVQKSYSFFLVLKNVF
>g13437.t1
MKFLQIDGPRDVLSIGCRMLKLFGLPRDDSFQLRFWFQFVFFFVFGIITRFLTDIDEPIALV
RVGSEIVYAVYLLVQMAALYGRRDDLYQLVDMLRECVNKSYSGHIYTFLVRTNGQINS
SAVKYCKYFMGVCVTYFAMPSIATFVVCVRNLRNQTGEQEEYVLPTELNFYYLDIRYN
LLHYLVYIAAVGVLDVTGSLLLCTKDVVDFSLIRTTSMLFQITAMQIRDLPPLASQADFN
VVIQSHRNTLECATKLQKAMHSALMIQLTFCTAIWCLMLFYILLMGFSSKILNVSLLLLIL
TYETYSYCQFGTHFTDSADEVLRALQQLTWYDQPVTIQKQIYFMIQHSQRSIVLKAGKL
FPVNIAQFSELVKKSYSLYLVLKDVF
>g13438.t1
MGQLRMLLRDLKLFTDTGKREPLDDPSKLFPLSLKLFEMVGVRQHAANYIRLCIINIYM
LSSFFLPKLLLGYDTIPQCFRSIAEVSVTEEYKRILIKLNKAIHNFTKHYFNLTILVIVAMLC
STSFGAFYMYFAQTPGQTLSYPQIMEHRLYALDAHHNLLHWFLHQFFIMPALVILVVIY
TGKAGIFFGAIRFCSTVFSILVLKIERLHLLISNEQYAAELKEIIMLHQLAIRCSKLLQEILM
EILLAQFTGCVLIWCFFLYSVMISGINAEGVTVGAMLFSFSTETFIFCLLGNELTSQGEQIS
TAIYETDWYDRPVEIQKLIVPIIQQSQQRIGITAAKFYYIDFNRFGKVARTTSDSRIACQQ
>g13438_2.t1
MLRFFSINKPPGVPHIALKLLRLVGVSGVRTERYRYVPMFVVFLAFIAVPKIVFGYPNFET
SIIGLAELFFQTNRFVGVLLLVLYSDTLFQLVRRSESYTKTVLTEPSSAAHYLKATDAKIT
KITRLYLTALLVPANFYSYSPILSTLWKYYNAHENDTTAVEFILHMEENFYGLNIRETFSH
YLIFGAIMVPTSFLCALVGTAKLVSILSLIKYCTINFQLVTLKVREMVQQQRLSCEMKSIF
RMHQNAIDCADLLRTLTAPIMLLQLLLCILVWSCLLLYFTISGFNTQFINLFVLFLFDTTE
TFGYCFLGNQLSDESARVARAVYECNWETLSPKTQKDLQVALVRAQRPARITAGNFCS
MN
>g13438_3.t1

MEQFAEPIFKRALLLTTHCHIGCSSFNSYTPRSCNSFTGYIVKMNFLKLETPPGVPHIPIKL
LKVIGIIGRQRERIRILPIFLVYIFIIAVPKSFYGYPSFEVAIIGIAELFFQTNTFCGIFLLFLNG
YKLERLTDGARAFSQKVLQETSPAIVQHLNTQHKLIHKVTRVFFIVVTCAANFYVFAPIL
STLFTLYGPHYLLFSVIMTPTCYLCAFAGTVKVLTIGNFIVYCTLYFQLVQVKLQTVAQE
NSFHQELKAIVTLHQDALNCAKLLESITSLVLLQQLLLCVLIWCSMLLYFTVSGFNTNFM
NLFVLFVFDTTETFAYCYLGELLSNEVTRRDNSSGEVLMLSKFYFRVLTLPVWCTNVVG
NCKPTAIQKDLKLVLVRAQTPVGITAGKFCYMNMEQFGIILKTTYSFFVVLREQF
>g13457.t1
MGFKPCDETAVMPLALRLLRVIGVWGDLTRYRYILVFVCYCFGIVIPKVCFGYPSAEASI
RGYTELILETNVFAGMLMLYVVRYDHFKLLVVELRSFVSIGKMEADVGSDLNFTNSPYFC
PQSNLNVRLHKYTMLYCLYMCCVCTVYCFAPLWSNYSGYTSAINHQDNGSTVFQFNLY
LEQGFYWLDNRTSLVGYCICTVFMFPLMYLCAYTGTVKVVAVFNLIKYCQAAIQTVAIK
LEHLKTFRDVRRRTEAMSEVRDLHQRAMRCAELLELVLQPLLLMQLVLCILIWCTMML
YFSVSGINVKFINMFLLFLFVSIETFGYCYLGTQLSQESINVGKALYDADWIDYDAVMRK
KVAFMIMRAQCRVGLTAAKFCFVD
>g13457_2.t1
MEQFVADDRAVLPLILYLQKHISLGDGSKRLYYPIVFSAFCASVVVPKLMTHYTNLETFI
CSMAELVFVGNVICGTMLLWMEYVSFAQFIEQTTSLTKYLYRDHQLYSVREYVLQFNR
RIHRYTNKYCAFMGVLVVFYCLAPILTSLTAYFQSLVKSAGNDTSTTMAKVEFTLHMEE
TFYGLEQHTNIVHYILYTLAIVPMMFMTAVSVHTKLLTICCSVLYCETLLHIITLKVDNLH
RMPMGNAARDELADIIQVHQRTLACIALLVKALRPVLLMQLVFCIFIWCLMMLYFTLAV
SSPGDSICKCFNSFVCNGMQDIGVKFINVGILFFVITIETLGQCYVGTRLSKQAEELTKCV
YACGWQSMDREIQQELLMVLHRTQWPVGIQAGKFCFVDMERFQKMVNVSYSFFIVLK
DAF
>g13672.t1
MLIEECPIIGVNVKVWLFWSYLRRPRLSRFLVGCIPVAVLNVFQFLKLYSSWGDMSELII
NGYFTVLYFNLVLRTSFLVINRRKFETFFEGVAAEYKLLEKNDEIRPVLERYTRRGRMLS
ISNLWLGAFISACFVTYPLFVPGRGLPYGVTVPGVDVLATPTYEVLFVLQVYLTFPACC
MYIPFTSFYATCTLFALVQIAALKQRLGRLQLDTSVAGRSPGTLFAELKECLKYHKQIIQ
YVQDLNSLVTHLCLLEFLSFGMMLCALLFLLSISNQLAQMIMIGSYIFMILSQMFAFYWH
ANEVLEQSLGIGDAIYNGAWPDYDEPIRKKLVLIIARAQRPMSIKVGNVYPMTLEMFQK
LLNVSYSYFTLLRRVYN
>g14613.t1
MKLNKLSPRWNAYDRRNSFWLQLVCLRHLGLWPPEDSDEATRKRYIAYGWFLRVVFL
HLYALTQALYFKDVKDINDIANALFVLMTQVTLIYKLEKFNYNISRIQACLRKLNCTLYH
PKQREEFGPVLRSMSGVFWLMIFLMFVAIFTIIMWLVSPAYDKERRLPVPAWFPVDYHR
SNTAYGVLFLYQTIGIVMSATYNFSTDTMFSGLMLHVNGQIVRLGSMVKKIGHEVPTEQ
QLVKVAVPIDGEWIEMRKRIKHHSRVYGKTYAEVTECVLFHKDILSFSDEVQDIFQGSIF
AQVCASVIIICMTLLQATGDDVTTADLLGCAFYLLVMTSQVLVFCYVGNEISYTTDKFTE
FVGFSNYFRFDKLTSQAIIFFMQIHGYGRSVGRGDTSTGLHTAGPWYKSHPDRLPLYRTD
YGFNYGLNQVTKSQNWLAKNSRGCRAT
>g15209.t1
MRTAIELYHASFDRLKLSSRLIGAGLWEQKYGFTWGRIASLTQIVMFLGLHVWTGFKYR
EDALEMLESQSLICTGLALMIKYFTMIRNRDPVRELTGNIERETYTNYQETSNEYPVVQK
YSRVLYIAGHIMIGGYFGSLFIIWINPLFVVYFTEGRVMLLFFCEIPYVDWTVNRGYWITVT
LQIAFYVTGTCGLILVDYLCAFFTINGSLYVDLLRFHLNELSELLAEPGYQTRSSPEIIEKV

DHKWRICLVEHQHIVEYYDKFSDLWSMINLAQVGCSVFGICINMLIIFLVYLSTDWYAA
YAILFALFIDLSVHFVLGAIIERKVDELLISLVHFPWYLLDDRRQKEYKLLLLRAQQPSGM
SIAGLTPVNYETYTQTDWYPTYLFFLV
>g15209_2.t1
MFTQLTVYFVVGHIVELKIDEMYNKVISMPWYKLPVKEQKEFCFLMAKQQRPMMLTA
YGFHPMNFEAYMSLYVKFFVGIAKGNDVKLLTEKIELEVLERYENGTKEEMAVLERTG
RYLWIIFRMTRISSACAAGGFMLYPLFAYFTTGRVVPLLLYELPFIDCNTTAGYIVNMLF
QINLLVYGVMGIIFADFLYIMYATYAMTAADIFMVHLVELEILLNDPMQHDTTRSEVRE
LWLKCMYDHQLTTSLLNIIEDIFGLQCLAQVAMGVFTICDCLLLVTLTDWYPTYCFFLV
MFTELTVYFIVGHIVELKIDEMYHKVISMPWYKLPVKEQKEFCFLMARQQRPMMLTAY
GFHPMNFEAYMSYVG
>g15210.t1
MGVEMWTAPGKFLPASCYLSVQMAIYLISTGFTVAKYRHDPLHMMKVLVTLGTALQL
YVKFFVALTKGHDVKLLTEKIELEVLERYQNGTKEEIAVLERTGRYLWIIFRIMRPVCSS
TAVAFMLYPLFAYFTTGERVPFLLYELPYYDCSTIPGYVLNMLFQLNLIINGVMGFILAD
FSYVVIVMYAMTAADIFMVHLVELEILLNEPLKDRTKSSEVREMWVQCIYDHQLTTRLL
NITEDLFGLQCLAQVVLGVFTICDCMLLVTLIDEMYNKVISMPWYKLPVKEQKEFCFLM
ARQQCPMMLTAYGFHPMNFEAYMSSVHNSNGGPVSEAKTVPVFTWQDQGSNPIKIFYR
LPERRVDYFATGEIKSQKARNGRPRSLEVLVPRKKKKYTIRYFRLN
>g15212.t1
MGVEMWTAPGKFLPASCYLSVQMAIYLISTGFTVAKYRHDPLHMMKVLVTLGTALQL
YVKFFVALTKGHDVKLLTEKIELEVLERYQNGTKEEIAVLERTGRYLWIIFRIMRPVCSS
TAVAFMLYPLFAYFTTGERVPFLLYELPYYDCSTIPGYVLNMLFQLNLIINGVMGVILAD
FLYLLIAMYAMAAADIFMVHLDKLEIRLIVPLKDNKKRSDVREMWVQCMYDHQLTTSL
LSIAEDILGLQCLAQVMLGVFTICDCMLLVTLIDEMYNKVISMPWYKLPVKEQKEFCFL
MARQQRPMMLTAYGFHPMNFEAYMSVHLLDLLELLMDRETPTVPSWCHQSMLAHAD
STTTTDGETMDERRKSPRARGAGRTR
>g16275.t1
MKLLHINDPREVIPIGCRLLKLFGLGRSEKLKLLYWTQVVFYLVFSLIPRILMKIEDTVTL
LRLGAELAFVSYLYSQILALYIRRKNLYHLVDMLQQCANKQYSETIDTFLIRSNAKINKF
SVICCKYFFLMYILYCVMPPLVSTGVYFRNTGNLTDEREEFIISTEMNLYYLDIRYNVLH
YSFYTLIICLLTVTSALSLCIKDVMDVSVIKTTALMFQVTAMQIRELREYISQTQLNTVIAS
HRDTLLCAQSLQDTLNLSLLFQLTFCSAIWCLMLFYILMMVWKGLGFDSRILNVVILLLI
VTIETYTYCTLGTHLTDKGEEVLMALQQLAWYDQSVTVQKQILLMISRSQKPIILTAGKL
FYASVLQFSEMVQNCDRVQQKVSIPKNQPGPNFHIIIIIIGIIGMVETNMNEFDSGVMSSV
EPADNDVPQDSIASPFRRLVTSVVFRGSPSDFPSHVLVGRDPGDYRRTNVPYAMLMIRA
HLLRSNGXVVQRKTKPSTAVGSLCLETNFMLHHPVRRMPPGRTARAVNKSFHFSEPHQ
NTHAESFALECECEQTRTIFTLIFSINFTCPQANSRIPEEKGGKCSV
>g18340.t1
MFRRLTQTLLLQPSSYRPDTSDFTVMPLGLRLLEYSGLWGDPRRKLSFALMLIGTILLLIV
PKIVLGTGSDSFDSIARSTAEFIFCYNNYLMMAIFAVQPKPFEQLIGTVQQLFDKHRTHQ
QHGTASGYVVFVNRQIMRFSRLYITVQGVYFLIFNLLPAFVTYHAYFTSNGTTTVEFLLP
VESRFFFMDIRHSIVDYTIFSIVACPAFLFTAYLTVVKGLVFIGIIVYNTLQYQLVSKAIRE
LDADELDTAQFRHRLTEIIDLHGMATRCTKLLDSVLNLMLLVQFTNTVLMCCLFLFYISK
NVNSGAVNVLLLFLALTVENLCFSYFGNRLSTENHNVAVTIYNTAWYKYPTTYQKQFQ
QMIRHAYIPRGITVGKFYIVDVASFGQPETLRAV

>g18340_2.t1
MAKILTRTLSDNFRVMPYNLRVFAMVGLWGDRRKQYRLCVVLLGALVVIIFPKPVRISD
RHPFESIVRSVAELIFAALCYLTVIILAIKSEPFRQVIAKLEQTLALFRDTNDQCSQLIVEVN
ANIHRFSLSYAKLHLFYVLLFNVAPPVYNYPHYLLQSKLPPANRTLEFMLPLMQDLYGL
DVRHNIAHYTISWMSITPFCAFFTLILWYKGSLFMLIRYNTLLYQLVNLLLQQFEYEGPG
ARLRQKHHRLQRIVELHHRAIECTTLLDSILSLILLIQFVGCLLLLCLILFYVSRNHNLIVIN
LGVLFSSIFIEMMCFSYLGNQLTEENATISNSAFNCHWYEEPIVIRKYFLRIILQAHRKATI
TAGKFYNVNIVTFAQLIKTSYTYYMLMKEMF
>g18446.t1
MKSRFRLSGTDKKRSAELVSVADEGRISSYANCAIPAMEYIYRWIVRDDKFAEEVFVRY
NFRKLRYDRDASMVPLASRVRFYALELFFLLQISAIVWDLATVLNDIGLFGDNMCILAG
LLLSMAKKWHCVLNIRELSECIDQLQTYHEHYLQQGERFVRRMRRHDLQERVLQDASS
LLAMLLAACLLVNALFSNGETLILRATYPFSTSTTLGYGCVFLCQAFLIVYLLFTIVLIDC
TGAQILSQMALLFCMQRMAFETIGADLTLPPEGSLHGDQQLRRSVHQLIASHQQLLSFC
DRLKRLYEPNIMAQFVCSMLIICLTAFELMFAKGDPMQMSTGISDATIRCNWIVLDDGL
KKDLRFTTMRSQKPFVIDVYWLFPLTYETFIAPRTRVTLARVLLE
>g19986.t1
MHLTFEETLKNTNIMLLMMGIPPCEEPYPDGVIAAIRRNLGFISSFLLLSYTTFGELIYLLQ
MFEREVNFLEVTFQAPCIGYCIIGVLKMIILAGRRNMIAELVAMFRTKWSQAIVHDEHW
KVCEDTMKPAIRVTSVTALVNVVMGISFTVLPIAEMIYHHYYTGSWNRQLAFNIWWPF
DVLSGAKYFWFVYPMYVVIGFTGIIIHMAFDCLFCILAAHLCMHFRILEHNLTHVVEVET
RWEDSNDGNMSRLQTTVVNHQALIGYWCSVFMQSVFGNALFVNFLGSSIIICIQAFMITT
VSGYTLVKFILFILCFLIELLMLCAYGEDIVQSAMSNGYVRLRLWRINLGAEGAAARSLI
GEGNPVVAL
>g20871.t1
MPCLEEFPAYRFWYMRYCGEVERPTWWFRCRLWYAFILLCSFGPLHVLYLVKTPALDL
MDTCEEIMLLQLCATVVLKFSLFVTHRAEIYDLVEGFKSILKGISVDEFPRFVKCNEVHA
KLARLYVIGTTIVLMLYELVAVVTSVTISLQQHEVHFVTPFNFPFNYQHPVAYAVCFLH
NLDAMLITVFVSVTIDACYSEMASNLSIHFHIVRERFERLDISAGQPFAERELSEVISYHG
DVLTLAQRMTELFQESVFYLLLLLSTILCLLGYEFVMVNNIYKRAQVATLAGIMIGQAVI
YTYHGSVIRDQSVNVSDAIYGTNWYEARTDRKKQIHISLMRAQKPVIVKSGFIEASLPTL
KKILSSSASYITMLMSLEADVSDDGLAEELFKMGLERITVEMHQLITR
>g20899.t1
MFALTFLFNCDSIHMSIFISGSVDTCFSELATSVTIHFRLIQRRFRALDFTAGAAEAELEAV
VAYHKDVLQLCLAMTNLFQYTVFYLLLLDSVLLCVIGYQFVIFMNTPRVLMLASMAFV
MVLQAVIYCYHGSMMYDESLKVADAIYQSNWYEARPAVQKRLRLCMMRAQKPIVTK
GGFMQATLPTLKKVCYGEAETLTR
>g20901.t1
MSGPDKTVDFFRVQSICLRAIGIARVDTRVGRAVFAVSFFTVLVMKLGTVMFAVKHIDE
IMLLCDCLGPTFTAYLGLVRQYNLRLHRAEVWSIVDEFVALRRLLKSNEIRIVKKYNRID
RFLAWAYLITAMSTGVLFVGVALVLVVLSDRADWKLPLLMDVAIFWVVLDCVACDSS
FGTFSSCLVAHFVIIQDRFESLQFGTGNSALGQLIEYHKHILHVSDRVIDAYKNVILNQLLI
SSILLCMLGFQLVISAGTSIMVVYVAYGTAIIIQVTYYCYYGSQLYHESTLVHDAVFKSN
WYEADVRTQKMLINCMMRAKKPVNAKSGFTEASLPTLKAVA
>Afun_g15.t1

METSIGKHHSLERFRATVAWQNRVLALFGCYVYVAEHRVTSRIIVICFISISFIVLSVYSA
VQSWGDMGQVVLSMVAVFYAIVGVARLAVAISDPAGCYRSIKLAEEMYQHANGSDPA
ECSVLARYTDLFCKSVQLYTFCFILSVVLVTVMPFAFYLFRGERFLPLGIVFPFTEGENVI
GFWCTLGVQLIYIFSGPLALVPSQNIYFAFIFNICLQYELLIERLKQLDVTIRNSGGFRENS
HSLPIRDQLVKIIQLQQRSTNYITHIEHFYQMQSFVEFLCNSLQAALTLNELHRSFWLPGL
FILPIAVVQMLMLCCLGTLIELKSDQFTDELYDIAWSEMELPQQAMFKYVLQSAQQPMR
LTCGRFAVINMNLFLTYYEMSYGLNVEMHKQIRLAPSQS
>Afun_g655.t1
MALQRIVQCGLSFLQDHFNVGHPSKHFCLLRCLDVVSPAMLIQRPRSNLEIGIKKICLSILI
AHLVGMAYDLTHQEDVRVVMDIFSMLTMFSSIVVRNICLKQYQSHINAMELLDENPNF
EVGTPYAETIRRRTVKQNNRYLGSYLVCQFLCATVWVCQNMAVKDSFVTIITHFPIDLS
ERAPTFDTLTQLCYACAGYFWAWYHAMGQMIIIVLLRFTITEFRVFLHSLATLDTQIHDQ
LLLDPEGNEERVVREVVYKHARRHSELIVAVMHMRMIIRNYSLVHFSFYMIIVAAFMAR
VLVIPGSSSLGMAVPLLATAVFFVETFGLCMLVETLVQLNRSVSFQLYGFNWTRYLPYG
NSIKRTMMLMIMQANNTNDFSAGGLTTVSADLFAKTCRLIYTIMMGMANLAT
>Afun_g657.t1
MDTSGQPSNICFLRWIHWMDTINGIHLHDESRLARTFQSAFYILQLGQLLIAYNFGVICTN
TISLEEFAQQFNQFGGILLTFFRVITINIYRTELKDTAQFINTSEFHHLNERAEQIRSRTIRPA
GRFFSTLLGLQIVTLIFWFILTELQAYKQNVLLPTITYLPFDASDWPTLLKVMFRLYVYLS
YTQLMFTFFGSYVITSSYLLTLTIELRILNDSYAGAPADPQQLVTFLKDRVVYKESLLQHI
GTIKRQMNVSILLELVLIVCLLAINGLRLCTTTSDLSELVLSCSMIMIYLLEFFQYCWQVD
EMELLHEEQASAVYSTPWVGAMRQTKALLLITIRMSQVPLRFMCGGMYQLSTELFTAV
VQFIYSLVMMLLHFK
>Afun_g1524.t1
MTLRSSSEDDPTLTLNFRTLERVLRFVALWPTDYNPYLPKYLRGQYLLSQLIDSCYLLF
WFFICVHIAAFHIVSILTIDFSYDELFLTLITTSIYSIMIPLSAYLRLYESNIRQLYEFATKHF
RKRSAAGIHYISISSSIRFTNRYQFWWLVICVLGTMHWAIYPILSQERTLPFPCWYPVDVQ
QSPMYELAYIFQVLGQLQVSLVYGLSGALFMVLVFMTCSQFDMLCCSLTNIRQSAMILN
GSYQTELRHCQGNHEIDTREYVLEEVFREDLENVQSTKTISKLDQLSPSQTFLMELSSELT
CVLEDCIKHHLLLLRFCQLLENCYHPYILLKLFQILLLLCFLAFMATVESLSTMKLINVLE
YFMLTMTELYLYCFLGHILMSQGLKVGDALWKSPWHLCGASYRRRMLIILMNAQRPV
RLTGWKLYELNLETYYTDVLGIRKDGFIDSEHVSSTGDIVVIE
>Afun_g1526.t1
MQVQPTKYVGLVADLMPNIRLMQASGHFLFRYVTGPILIRKVYSWWTLIMVLMQFFAI
LGNLASNADDVNELTANTITTLFFTHSVTKFIYFAVNSENFYRTLGIWNQTNSHPLFAES
DARYHSIALAKMRKLLVLVMATTILSVVAWVTITFFGESVQNVFDKETNETYKVVIPRL
PIKSWYPWNAMSGPAYIFSFIYQIYFLLFSMVQSNLADVMFCSWLLLACEQLQHLKGIM
RPLMELSASLDTYRPNSAALFRAISAGSKSELIINEEKDPDVKDFDLSGIYSSKADWGAQ
FRAPSTLQTFDENGRNGNPNGLTRKQEMMVRSAIKYWVERHKHVVRLVSAIGDTYGPA
LLLHMLTSTIKLTLLAYQATKIDGVNVYGLTVIGYLCYALAQVFLFCIFGNRLIEEVRSFR
TFHHICIPKAILVRHIFIYHFNLGLGRYVQSSSVMEAAYSCHWYDGSEEAKTFVQIVCQQ
CQKAMTISGAKFFTVSLDLFASVLGAVVTYFMVLVQLK
>Afun_g1656.t1
MSALVSLVKNQIQRVTDAGQLVIINGMDRFIGFFSWDVQQRITWLKLVLIVFAVAYEITA
IVAMALASMRGMFTERSFTMSFVTMTGAMCIVIWVSLAVFRRDLTATVAFLQQRQRTI
HNEKAPRKALLDRVTRYLWLFYLQNIAQVFFWINLLRDCSPLAVFELPLLDSANVLLYP

VAMTLMSLMFIHTIMIVSTLLSGLTLEFYWLGQEFEKVFGECSTIAAIYHKRYWDALERK
IRTCVTEHQLLLAQISTLRNNLKLYLLLNLVADFTLITFAGCQMVMSNEGDQHLYSILAA
LTACLNMLNFGGLCDLLKIQVSSRDRAKQTKDFNQHPTTHKHTRILQVHAIKFHLYSSQ
WTDYLRPVSGPLYQRCRRIRSSMLVVMTRAEHELRISCGSVYDMSLATCWAVLQFSYS
VFTLLLSFFENTPRSQ
>Afun_g2456.t1
MDTLEQFYKYEYYFSMLCNVIGFNILDKGWKKTYRTYISFFLCGQYFLWMVWSIIIASD
TFELFKSLSFLGFFFQCSLKMYYSSAKATQYGISFDGLKQTIYIGHTNGTVEQKSVIKRIFL
VVQLITKVTTVLYTSSLFIFSLYPLYMYFIVGVKVTIFPLYIPGIDIYSAYGYGITNSLHMLI
AVYGCIGAIASDIAFMMFVLHFVTYGELFRIECEQFQQDLSEATERWERDTPEYKTFCRQ
RMREIYQYHQNVILFLESLQECYQSLSVVQVGSCSFSIMFNLFLALTTDWYATYSFMFIS
WFQLFIFSLFGTIMQVMNDRMLRYIWNLPWYMLPSEEQRRFHFMLARSQLPAEMVIRS
VGPLNMETFTDIMQKIYSAFTMMYSFLVDLS
>Afun_g3082.t1
MLSNVAPVSDFRDTKNWNIFKLQRTILLIFGLWPADRLVRRWYMKVLIAINLLTLALCM
VGEFLHGLYAYWEGDLSETIESICPTVARISGFLRMVFYLINEEKIEQVLNNIRQMLADK
HPREKEVTKRLSKLGQQFTFYLFLMMFFAACLYGVTPFFIMAYNWSQGQKPLVKLLPF
KLALPFDSQNSFYFVLTTIFLNYASAPTITSQSGSDALFAGICLYLYGQFQAIRLEVEALSV
TLDDRSLKESFSQTQRINKELRRISKRHQEIIDLVAEVRRAFMPNVLLVYTATAIIMCIVCI
ALLVVEGIYKLTYLPYAFAELTLLFLYSYSGTIIRDSSEAVQTVAYDFPWYRFDRNTRHLI
QMMMIRAQYGSNVDVPFFETSLASFSTIVRTASSYITLMKSFL
>Afun_g3252.t1
MARLVLHEVRYVLMAMVYISRCLVEKIQNSVIDKFIYWVLTLIPIAMLCIPQFSYLLVDT
KDLVEFVTVLVPFTEIVLTNLKMAICNMKREKIINLINEMQAEWNEYQKSEHLEIQRLITS
TAKKTRIFVIIYTTAFVVIVLEYASMPLFKYTFLNVFMKNHSNYTVTIPYKSSTESTTNFS
LTYFFIMIAVYILALTLSGFDSLFATLAMHITTMFKFMKIQIDQLELDLRAGTNSTEMHAK
LKGIILKHKTNLSLMDQLEDCFSFYLMVQFLTSSIVVCVVLYELTIVFGWNEDTVKTLTY
LPGAILQLYLFCWYSQNITEEASMVSDHIYNIPWYLSDRALQKTILTFMVKAQKPTGVTA
SKFYMVTLQSFQRITSTSYSYFTLLQTINQQ
>Afun_g4366.t1
MLVIRGEPIGTEFDHLLYMNFLYFPTVGFVLGCSIVHAMLMGFMGEMELLAVCLEEVFK
TVENQLIAQKALHDQAQFWSILHEQLSKCAKRHSEIFTLIHKIYYVRYHPEDINYISFVCG
GITVLVAVLLLMLIFTNHYDAFRELGDFLNDRSFARDHPLAAKIRERWYRWSNWLILGP
QLGIMLILVQTWISQQHLKKHAMLVIRGEPTGTEFDQFLYMNFLYFPTMGFFMACSIVN
AMLTGFMGEMELLAVCLGEVFKTVENQLITQKALHDQAQFWSILHEQLSKCAKRHSEIF
TMLPKLQRMASFVFLQHHTFSLGLVTAGCYVTLRGPALRENIVLSEYPISVVLEYFIFCM
LVEKLQDMISLIAFALLLVNANAQQYGQQLGRRSQERLNQLRSYDDGVRSSRNYNDQY
NEQRYSGRNQDQDLQDRRESDYDRDDYSYGYAVRDELSGDIKSQQEVRNGDRVRGQY
RTLESDGTERIVDYTADDVRGFNAVVRHQPSVGSRAQLVHTLQPAVLLRQPTVGQLVS
GNHRPALLTTPQQTSTVLLRN
>Afun_g4399.t1
MTIETSSTDKYRWSDYIRPVRMTIWVWKICGLYNAKPQAPLYRAYRIVFNVVLMVVYL
FTLSFNMFLMQTFEQLVLYTMYIVFTEIVMLLKALITYHKFEQICDLFRQTLDSDFKPMD
AEEEKLHRKGVGEINYYFYLYILTTNLAIASSVLYLLHQDYRLPYFPWMLGIPYGPTERL
NFGIMFAYQVVGMYFHMWINIAIDIQLIYFLGMIGIQLDLLGKRFRMIRTSEQFGQSFVQ
LINQYQKVQRMTQDIEHLFSPAFFAQFSASGLVICATAYKTSSMFNVYELTAIQNLLYML

SMMFQMFLPCRFGNEVTRKSHILRTSIYSSTWYEMRLKHRKTLHMLLQRMNKPLTLKA
YYFFNYNLQAYTTVLKHYIKCHA
>Afun_g4400.t1
MCETGRFIFPMRFAMWSWKLCGFFNAPVPKSIPYRVYCYLFYFWIMALYLFALMLNVF
VPQPFEQRVFFIMYISLTETAMILKTRTIYRHFDTIWSLYETTLGPSFQPRDEFERELQNRT
LQVFNRWYYVYIFVSHMAAFGTGSHLLSSEYRMPFFPWFFGVRYGDDARVAYYTIFAY
QCFGMYFHMLLNTAGDTQLCYMLQMIGIQLDLLAKRFRSLSSSEEFDRSFVPLVQHYNK
IHRMLCRVENLFSLAYFVQFSVSGLVICASAYQVASMLNLNDFSKLMNVFYMMSMTM
QIGLPCYYGNEVTLKSYALTNAIYSSNWYAMKHSNRKSVQMFLVRTNKPFAATAFRYF
NFNLPAFTTILNMAYSVYCVLQRKAKNV
>Afun_g4592.t1
MEPPSVGLVQFESFIRVPEIFFAMIGVTRYGEPKRTLRAHLKQALFWSSCANTGFCLIIEHI
YFVKAAGNFTNFLQLTALAPCMGFTALSFVKIMTIKLNETKLTDMLHRLDALFPKSSVL
QERYGVFQYNRESQVVMKSFSILYMILIWMFNLLPLVSMMEGYFTDGSWHKQLPYFM
WYWYDWHQPGYFVITFLHQNWGGFVSAVFYLSTDLMFCAFVLLLCLQFDIVAYRLKH
ALPDDHQELIECVRIHQAVIELCNELEHMFSPSLLVNFLSSSVIICLVG
FQATAGITPADLFKFVLFLVSSLVQVFLLCYYGNKLIVASSQIPYSAFEGQWIGASVSYQ
RSLLFVMLRSTTVQKLTALKFSIVSLASYSKILSTSFSYFTLLKAMYEPNEKNVK
>Afun_g6426.t1
MAKFLFWKTKVDKIFITREEKLKKLLPSLRMIFFIFGIFCAWPDVNMEKNALWWYRLKG
ILFRIFFIYLGTASQIVYNFTVTTREELFGGMFVLLTQLVLILKMEFFYKNVSKIQTLVRRL
EGELYLPRSDEELEYSIAHSPTVAARWSISAPPTDDPIYPMAVARMESEIIQLLKNCQMPT
ESETETEPVAAKGLSDRVVGVGFRPSVRLRFSTPKNGTRDVANFPPVWWKLEKSRKKR
KAKENIQIAQDMTALSSFVALDCRKFAAGGD
>Afun_g6763.t1
MKLNKLNPRWDAYNRRDSFWLQLVCLKYLGLWPPEDSDQATRKRYIVYGWALRVVF
LHLYALTQALYFKDVKDINDIANALFVLMTQVTLIYKLENFNYNISRIQACLRKLNCTLY
HPKQPEEYRPVLRSMSGVFWLMIFLMFVAIFTIIMWLVSPAFDKDRRLPVPAWFPVDYQ
RSSTVYGVLFLYQTIGIVMSATYNFSTDTMFSGLMLHVNGQIVRLGSMVKKIGHEVPPE
RQFLKEAPPSDDEWKEMRMRINYHSRVYGKMYAEVTECVLFHKDILCFSDEVQDIFRG
SIFAQVCASVIIICMTLLQATGDDVTLADLVGCAFYLLVMTSQVFIFCYVGNEISYTTDKF
TEFVGFSNYFRFDKLTSQAIIFFMQMTLKDVHIKVGSVLRVTLNLHTFLQVRYRLYV
>Afun_g7223.t1
MKLLQIDDPREVIPIGCRLLTLFGLGRDEKPKLLYWCQVVFYLVFSLVPRVLVKIDDTVM
LLRLGSELAFVSYLYSQILALYFRRENLYRLVDMLQRCANKRYSEAIDTFLIKSNGKVNK
FSVTCCKCFFLLYILYCVMPPIASSGVYIRNTHNLTNEPEEFIISSEMNLYYLDIRFNVLHF
AFYTLIICLLTVTSASSLCIKDVMDVSVIRITCLMFQVTAMQIRELKEHISQTQLSTVIDLH
RDTLLCAQSLQDTLNLSLLIQLTFCSAIWCLMLFYILLMGFDSRILNVLILLLIVTVETYTY
CTLGTQLTDKGEEVLMALQQLAWYDQSITVQKQILFMIHRSQKPIILTAGKLFYANQYK
CKLQLPRQDKKLTQLLSVRANPSLSCFPALPKGWP
>Afun_g7544.t1
MRSAIEFYNVSFNRLKFSSRLIGAGLFEKKYGFTFGRIASLTQIVLFLAFHAWTGYIHRQN
ALEILESQSLIYTGIALMIKYFTMIRNRDPVRELTGIVESEMYMKYQETCNEYPVMWKY
GRLLYFAGQIMIGGYYAPLFIIWIYPLFVYFTEGRVLLLFSCEIPYVDWTMTSGYWITIAL
QLAFYVTGVCGLILVDYLCAYFTINGSLYVDILRFHLDEMSELLADPVYQMRSSPDLVE
KGNRKWRQCLMEHQHIVE

>Afun_g8459.t1
MLIKCFAVCGGERMKPGYTRRNARLIFLVTDLILYLFVNAYSIAIVWGSLMDVVFCFVT
LGIAIQGLAKIEAFTCPELNDLHLYNVARFKLDPRFPEVEDALFHTATLCKVFIRILAVAF
SIVALAIYSYAIMMPLVEGELSLAFGFYLPFVDYRTPIGFAINWVYQFIQVSEGCMGLMA
CDSCLLFLIVNATGQMDLIIVYLRRLTELIDSNSFGQNDKQIADLLGEIVLKHLEHTKYVT
DMDKLLKKQFFISFSCIIFELVASLAIVVRNDKLVEEIYNVNWYGLSTNHQKTLKQMLLA
SQHPIVLSDGFSAIDLFNFVEVFSREENYD
>Afun_g8762.t1
MKILDCPLLSVNVRVWHFWSFVLVHNWRRYISIIPVTLLNVFMFADLYRAWGNIEEVIIN
AYFAVLYFNAVALNDEVITKLVATYTKRARYLSISNLILGAVISSCFVIYPLFTGQRGLPY
GMFIPGVNNFNSPQYEIIYLTQMVLTFPGCCMYIPYTSFFASSTLFGLVQIKTLQHQLRNF
RSENLAEKTKGLNRKLQKLIEDHKRIIRYVQDLNDLVTYICLIEFLSFGLMLCALLFLLNII
SVMAQIVIVGAYIFMILTQIFAFYWHSNEVREESMAIAEAAYSGPWVNVNDAIKKKLLLI
TLRAQRPLEITVGNVYPMTLEMFQSLLNASYSYFTLLRRVYN
>Afun_g8764.t1
MLIEECPIIGVNVKVWLFWSYLRRPRLSRFLVGCIPVAILNVAQFLKLYSSWGDMSELIIN
GYFTVLYFNLIKNDEIRPVLERYTRRGRMLSISNLWLGAFISACFVTYPLFVPGRGLPYG
VTIPGIDVLATPTYEVVFILQVYLTFPACCMYIPFTSFYATCTLFALVQIAALKQRLGRLQ
PDRTAPGRAPGTLFAELKECLKYHKQIIQYVHDLNSLVTHLCLLEFLSFGMMLCALLFLL
SISNQLAQMIMIGSYIFMILSQMFAFYWHANEVLEQSLGIGDAIYNGAWPDFDEHIRKKL
ILIIARAQRPIAIKVGNVYPMTLEMFQKLLNVSYSYFTLLRRVYN
>Afun_g8951.t1
MRFKPCDDTAIMPLILRLLKVFGVWGETRYRYKYILVFMCYCFGIVIPKVFFGYPTLEAS
IRGYAELILETNVFAGMLLFYVRNDHFKLLVAELRSFVSIVFRAYQPKFIRSNLVKLNLQI
HKYTMFYCLCMCGVCTMYCMAPLWSNYYGYTIAQAARNNGTPFEFSLYMEQDFYWI
DSRTSLLGYSVCTVFMFPLMYLCAYTATVKMLAVWNIIKYCQSVLRIVVMKLKYMKTF
PDPREQIDALSEIRELHQRALRCAELLELVLQPLLLMQFVLCILIWGTMMLYFSLSSINVG
QELYDCDWSNFDGETRKRIAFMIMRAQCRVGLTAAKFCLVNMEQFRADDRAVLPLILY
LQKRVSLWSDSK
>Afun_g8961.t1
MDQMEMLLRDLKLWTNIVSITDEYKQILITLNTGIHKYTKYYFIFTNCIVLAMFASSTAG
VLYIYTSQTPGQTVSFPLMMEHRLYILNAHHNLLHWFLYHMLLMPAILILVIIYTAKAG
MFFGSICFCSTLFSILALKIDRLRFLITTEQYTAELKEIIVLHQLAIRCAKLLQKILMDVML
AQFTGCVLIWCFFLYSVMISGVTAEGFTVATMLFSFSTETFIFCLLGNELTNKGEQISAAV
YATDWYEQPIKIQKLIIPIIQQAQQRIGITAARFYYIDVKRYGKNLKTAYSFYLLLRDIF
>Afun_g11464.t1
MYRDELYSTVCLGDMSQPNEFPAYQFRYVRYCGVAEQPTWWYRCRLWYAFILLSSFCP
LHVFYLVKTPLSDLIVTCEEIMLLQLNATVVLKFILFVSHHEEMYNLVHSFKSVQKCITV
DELPRFVKCNEIHAKLVRLYVIGTTIIVALYELNAIATSVSLSLQQQHVHFVTPFSFPFNY
QHPVVYALCFLHNLDSMLLTVFISVTIDVCYSEMATNLAIHFDIVRERFERLDISADQPFA
DRDLKQVISYHSDVLSLAQKMTHLFQASVFYLLLLVSTILCLLGYEFVMVNNIYKRAQV
VFLAGIIIGQAVIYTYHGSAIRDHSVNVSDSVYATNWYEATTAIKKQIHISLMRAQTPVII
KSGFIEASLPTLKKILSSSASYITMLMSLEPDV
>Afun_g11731.t1
MYRAHLTFEETIENINIMLLMMGIPPCEEPYPDGVLGAVWRNIGFIGSFLLLSYTTIGELIY
LLQMFERDVNFLEVTFQVPCVGYCMIGVLKMIILAVRRNTIAELVGLFRTRWKQVIVDD

THWKVCEDTMRPAIRITSVTALVNVVMGISFTMLPIAEMIYHHYHTAIWNRQLSFNIWW
PFDVFAGAK
>Afun_g13510.t1
MYFFHKSQQYLELNYNRIYDVFHWFLKITLLRIFDDDFLVAPIPTMVFHFYTSEIVICLVF
VLLHALSYRNDLDAVILSVSAFVSAFELFLKINGMVYRRKEITQMIRTVLNDRSYLNGP
MEAAICAKYQRLASTAIMLLIYPLLVGGIGERILPVGFSLPLVDYHKNPWYLINFVLQIVQ
VNWVALVFIGLDGPFYLFVCYSASQLEILIVYLRQIGENPDNVADQRRLIRKVFDIHSGLS
NCSKIYREVYLMQVLCSIVHICVSLFHIQIKFKNGSYGMLLTNVNKVWLFCYCGELVVS
KAADFSTAVYANQWYQLWNRRDLQDVLFMLRNAQRNYGFSVGGFGFLSFATFTAVM
W
>Afun_g15671.t1
MATKYGTQILYTRSRTTIKFVVSIIILTLTPGVQYILEGEKLMPYTIIIVFTDPEITSHFLMNI
TIQYYLLVVGLAGFIAAESMLILFVTSVAGYADVLKNKIDEMNDVLLEAENSKDRTPVK
LKLREILLLHQRVLEYENDLEKGYYLNNWVQVASIVFNLTGALFGCYVSNSFTMYALAI
AVVIQLFELCLLGTILSIKNEEIEQAFYNSLWYLMDRIEKKDFMIMFHKSQHAIEMTVAS
>Afun_g15671_1.t1
MAPLNIVLFIAGVAKFYTALRYHKFFEVMYNRLDRFHYEYRHHEKYNSTLLLLMERICL
VTKLITVQLVVSGLVLALTPVVQYIFKGEKLMPYAIVIGFTDPEITSHFLMNITIQYYLLFV
GIPGFLAAESVLILFVTSVAGYADVLKNKIDEMNDVLLEAENSKDRTAVKLKLREILLLH
QRVLEYENDLEKRYYLNNWVQVASIVFNLTGALFGCYVSNSFTMYALAIAVVIQLFELC
LLGTILSIKNEEIEHAFYNSLWYLMDRIEKKDFMIMFHKSQHAIEMTVASMAPLNVVLFI
ALRKPVSTLSIPGSMKTPCAGNRETVSAACTEEAGIALVGVHSEYPRYMEERGLGGGVC
KGPGTIQGGPGQKHKPNVGYRLGRRKALFEKRKRISDYALVMGMFGIIVMVIENELSSA
GVYTKLDSLVRPEQKTKNRRTVSQKGLEWFFEISNWELDDTNTTHQSMWL
>Afun_g15680.t1
MAMYYWIITVGSDITTIYNLLTAYGQLDVLMTIIDELDEQLERNECPEKIRAKIIEIVHQH
QHHRSYLQQLVDFLNPYHFVTLGSTVPTMVISVLGLVLLDWYPGAAIVFLGSVQIFYICF
LGTSLEIKTDALTLKVGAIHWNKLSTRDMKYMNLVLAMTQKPKMLAVATLPLNITAFL
KVLIVIVLIEVVRKNREWV
>Afun_g16016.t1
MFIVERLRTAVRRQLERVSLDPRRQHTAIVEGINWVGGLLGLDVFTPNFSAGNLHLRLV
LLNTFAFFWINLYNLSTTYGDLVEFMYCLETLLYVGIACIKIYVFVKHKTLIVNLQKFIEQ
FFESFHGDPEQDALLVRSLQNTYLLVALFGFCSCSAAMLMFVYSLMWSIAVEYTLPLGF
FIPTVGMDHLEGFALNYAFQLFESTLMVIGIISSECAFFMMLQNACLQVDMLLLELQRLG
RLGASNTDGHHTGEIRTRIQAIIVHHIDHLDYSKIMCRLFEVHFFIVFGCIFCQLVSIIVVIV
AVPDWYPGYFLFAMLTVQLFFSCALGQVFDIKCEELTVAIYNVPWYNMEVCDQKAMR
LLLLASQHPGRLSYGFGTVNMRAFFEIYRKTYSIGMMMISVNEEN
>Afun_g17598.t1
MVPFRILERPLVRKYWDKFFTFTDSVDYFNLLNTFGTLFALHYHSPESRWTVKKLLWTV
YRTFYLLSYLSYCYKACWMFSNWEYSTASANVLGALGLCSGALLRLILIEFNYPTIRQL
QAFLNDRTYLHEDKWAWIQRSQLYRQNNRFLVVLITAITVESLCFLARLLLTRPEFMLQ
YNGRVLGGSAVQIVYGMVTACWGIIYVLSFIVFYMLLAGFRLEMELLGRSFQQLEDTLL
LDHEHPCIMEDEDERSYWSKLQAILTIRIKRHVELLENLRLFRSIVAPFAFLQYYCTFGLI
ADSFFVVSFEGFTGYSMAYVLFASFLILESLLLCRGVEDLNDLVSV
>Afun_g17599.t1

MALFVSRIYRKLNNLERKFANDPDQFVILRYLTFLFAIRYDRPSGVLQRTLWYCYRSVLL
CVFSSYCYKAYWHMTHTAYNVSMFNILGTLWIFVGALVRVILFDRSVLARLERFLNDRS
FRGMEQKVITARHTVQRQNNRYLVAVALALLLETFIFLGTNLMLQPEFMLVYNGRVVG
GFAIQILYGFTTCYWGSLYVMIFFFIYVMLNAFREEMSIVVESFTHINQVFDHYHPYSDA
PNTSTAQEQEFWDELRYRLKKNVQRHVQLLEHLVAFRAILGPFSFIQYYGSFMLIAYYC
FIMMYKGITSLTVVYIGFIVFLIVESFLFCRIISEINDLVSHAQIGTVLYDMEWYNKLRFST
RFASDYRHVRSTMLIIICRTQTPLSFTINGLGTISMSRFTDLLNSSYTFLTMLVQFKREIAS
KLMEKAEN
>Afun_g17618.t1
MNFASLESTRIKFLNCISRYTECSDFFIIQRYFEKIYAIHYNARSWRDRTLWYLYRALYGL
IYVSYVYKTYWVLHHWENSLSSANILGVLWFFSAVILRVVILEWHYPLMERLQTFLNDH
SYQREDPWAVANRANFYRRTNRLILAVMVINFAEIVCFTATNVMKLEDFMLQFRGATI
GGWPVQIVYGVLTMCWGGMYCMGFMVCYLLMCIFKLEIDILIHSLENLGKSLNSRQDL
TSDISDTFWDNVIHRLRPHMQRLEDLLIHLQLLQAVIGPIAFVQYYSTYLVIADCCLILVS
HGLSSYSIVYVISMTVFLTEFYFLCHGVENLRNLKLRVATVMYGFDWTLH
>Afun_g17618_2.t1
MQSSNQRFSSQYYHVRRTFLLITAQSDRTIHFTFAGIGEISMNSFAQLLEKTYRLILLPED
NTSAAVVIASVWGFTEGTLRIGIIELCYGTLSKIMSFLNERSYRCQDALVRQQRAALFVR
NNRIQFTLVATMLIVAAWFMTTQLFSRDAFMLQINGQVVDSAAVQILYGLLCNVWGLI
YVLSFAIFYIIMNTLQLEMMILLDGIASVQFTVMNRAKRQLEILEPSGHSSQMQQQVFWR
MLQSELNKNISRHVNLLDILKEFSSIVGPFSFVQYYGTFALIADCGFILSMEGLSTNGMIY
LIFVTVLIFQSFIICRGIEKINDLNEAIGHELYAGFNWPELLQYDERFRYQHAAARHSLML
VIVRSQKGFQCSYGGLGGISMERFAQLMQKSYSLLTLLLQFTK
>Afun_g18528.t1
MVLPKLDDSFAVMPLLLRLQRFVGLWGERRYRYKFRLAFLSFCILVVIPKLAFGYPDLE
TTVRGTAELIFEWNVLFGMLLFSLKLDDYDDLVHRYMDIATIAFRKDLPSELANYLVHI
NHRIDKFSKIYCCSHLCLAIFYWVAPSSSTYLAYLSRRNTSIPVEHVLHLEEELYWFHTR
VSLFDYSIFTAIMFPTIFMLAYFGGLKLLTIFSNVKYCSATLRLVAMRIQLMDRLDEVQA
EKELIEIIVMHQKALKCVELLEIIFRWVFLGQFIQCVMIWCSLVLYVAVTGLSTKAANVG
VLFILLTVETFGFCYFGSDLTSESLSVARAAYGCYWYGRSVSIQRKLR
>Afun_g18528_2.t1
MVLQRAQKPVGISAGKFCFVDIEQFGNLTGGYSDIHQLVQASVEFLFNCNIFVSSLLFAH
KAATFRAFVRELKILAQFACSMSYKIKHTLVRFNRQADIFAKLQTTCMTVIALCYWVAP
LTSIYWFYLGSSNSTEPLQLVQHLEVKFYWLENRTILRDYVIFVLIMLPVVFMCSSMCNL
KVMIVSCSIAHCTLFTKLTVKAIEELPDATPYRRTSKSLSNVVLMHARLLKCIHLLNRTL
RSMLLLQWLICGLNWSISLVYLTNTGISLKSITVIVMFVLATSETFLYCLLGSRLATQQER
LERAIYAKRWYNYPRKVQRNILTILRQTQKATDITVGKFFRSVKVKS
>Afun_g18528_3.t1
MVLPKLKDEKAVLPFLLRIQSIAGLWGDRSQRYRFYLIFAYFIVMVVMPKVLFGYPDLEI
AVRGTAELMFESNAFFGMLMFSFQRDNYEKLVHQLQNLATIVLQDLPAELGQYLIAVN
RRVDRFSKIYCCCHFSMATFFWFMPVWTTYSAYSAITNNSEPVEHVLHLEEELYFLHIRT
SIVHYTFYAAIMWPTIYTLGFTGGTKLLTIFSNVKYCSAMLKLVALRIQCLAGAKREHVE
NELNEIISMHQRALDCVFLLETTFRWVFFVQFIQCTMIWCSLILYIAVTGFSSTVANVCVQ
IILVTVETYGYCYFGTDLTTESFGVALAIYDSDWHKFPVSTRRKLQLLLQRSQKPVGVTA
GKFRFVNVAQFGKMLKMSYSFYVVLKEQF
>Afun_g19440.t1

MVEHPIYAFDQLIKRQRLLLKLIGVDSYDPKFRFHGLTFLFVCLALFFFVVSLYDLFLFKN
DMFNFVYVLITIFFATIGLGRISVFLVYSKVLPNLLLQTYYTYQVIKNDERELRILGWYTQ
LFQRAVNGYTLVFIGTSIAAGILPIGIYLMTGERVLPYGVVLPFIDPSSQTGYELNYIYQVS
CIIWTPPGLVASECMIFALVLNICIQYDILGVQLQDLDELIRSQGPMREVMIAQKLRTILH
GQQRLFSFITTIEDSHTVLSGVEVLSLGLQIVITLFVLQFSLWIPGLVLIPAFTLQLFLFCLL
GTIIEDKGVKFSDGVYNLTWNELSLGDQKIFRLLLLSSQQTKTLTCAR
>Afun_g19440_2.t1
MTPICLNLFVNVLVSESPIDRFDRILSWQYHILRMLGMDAFTKRLTLNPLSVTIMTMAGL
FMVVSFYDVLVLFRGDLFGKSFVMATICFGFIGWGRIVGAWAYRSDVPKLMQMARDT
YLSGVNDERQIALLRWYTEIFWRGVMLYTMIFLFGAIMASFGPVLLYLYNGEKILPFGV
YLPFVDPNSGTGYELNYLYQMSCILWTPPGLTATQNIYFAFILNICIQYDVLQLHLADLD
VLVQRTDLEGKDDAVRAKLCDIIVRQRRLEQFVQSIEQVYSKQAFVEVLSLTFQLVLTL
YVLRASLWLPGLFLIPLCTIQLFILCVPGMLIQVKASNLTDTIYGIAWHELHQQNKRIFHL
LLHRSQHPSGLTCAGMANIDMNLFMSVR
>Afun_g19444.t1
MTLWNYFRQKLKPLLELQEDSDFFVLLNWQYIFYGVQLKTKRPWLRALFLLYQLLLPT
QCAIWLYRTWAAAYIEHNTTLALSLLCGQFALTSLLFRCVLFLRSYDQLQPVRSYINSKR
FLHGHSKAHELRQRAYRTNNILILGLMVYGLINFFIYEATGLQWHEIFRMPNYLMQTNR
PLAWTLHIIMHPMTLNGLGAFIASFLSMHTMLTALQAEFLLVEYAFVGLLKRVEEQVQG
VPTEDDFKQRLLWQSFNREIGKCVREHCEVVKHIRDVNRVNSFSITVQYYTALLSLAIDT
FFISYHGVDFVALSVLIFSVLLVFEWYYCCKLVEDLQATQNKRIGWTLYNDDWPAWLQ
HGKHQQKSLRQFRITLSIILLASQQSLSFHGSDIVEVSWQSFGGMLKTSYSVMMFLIELR
KLNR
>Afun_g19674.t1
MELKEEWILPDAVYDNPLLKRTLLGLKYYGLLLGHSQPYKKAHCFRGMVFTVSMVLFN
CTQYIDLWQVWGSVSDMTANAATTLLFTTTIFRIIFFYFHRARFNSIIQAAHAGIERILGD
GWDDEKDIVTSNVRYLNRLAVVFWCCALVTANMMCVYSLVLYLMYEEPSVNLDGLIS
NGTVPQQQYPTSILRSWYPAADGKDNHFLEIYLIQLYIMYVGQLIVPSWHMFMVTLMIY
GRTECSVLNYRLCFLDRYHAPGQADKPKASVPEHVDNDERRSLIIDCIKRQTNLVAFTRE
LEQLTRAAVFLDFVVFSVLLCALLFEASMTTSGVQVFIDICYITTMTAILFLYYWHANEIN
AYADQLSMSAYKSDWYRYDHGTNRMLQIFILYSNRPVKMQAFFISMSLDTFLAILRASY
SYFTILKQLTD
>Afun_g19685.t1
MDSHLQEKARKRLLERLYIERDFFHPFEILLALPGFHLVERFRKRSWMRVLFVLTRVIQL
LQYALWVDRFYLELIDSSGSSEKTLHYGNTLSALTMMLVRMFVVHWYMPNVEEFKRY
LRRQRRLRVTKPNSGTHRISYRKIVNIAIMFQLVGLADRLVFCFSSTYRQELYELPSNLVE
LGWPMVIVLHVVSFDFESRWIATYSVSITGMNSIMMGLYDELVDIAEEYQELFARSKDD
SSEFWSCLERNIVQAVKRHEAFISQLDQLKPFLQATFLVMFYSAALFLAVGTLLLITNGM
TVFKVIFSGFLFALLLECYWCCQLVDRLND
>Afun_g19685_2.t1
MNAQIGMHLYSLPWTTELQYTIPDDSRYRQVRLSLLIMMSKTQKSLEINCGGMFEMSRE
AFASLVKLAYTMLMRLLERLYIERSFFHPFEILLALPGFHLVERFRKRSWMRVLFVLTRV
IQLLQYALWVDRFYLGLIDSSGSSEKTLHYGNTLGVLTMMLVRMLVVRWHMPNVEEF
KEYLRRQRRLRVTKPNSGTHRVSYRKIVNIAIMFQLIGLADRLVFCFSSTYRQELYELPS
NIAELGWPMEIVLHVISFDFASRWAAAYNVSLTGMNSIMMGLYDELVDIAEEYRQLFAG

SKDDSEFWACLERNIVQAVKRHEAFISQLDQLKPFLQATFLVMFYSAALFLAVGTFIITA
NGTSTYDVILSGFLFALLLECYWCCQLVDRLND

**Text S1:** Protein sequences of all 54 ORs from *An. stephensi* and 42 from *An. funestus*

***SPO11***
>g3704.t1
MASTSADFFNDCESTDDLFASSESSLLLPSSFTDLSSASDEYSTDNFPFESLAWDKNLVNL
FNDFTEATTHEQQGTNTNTVTRDERQHDASTDLSQHLDASLHHTQTASSKRLFDGTLAE
RATIRTDNCLNSTVHNCSEPDERDHCNHVERWHDVGKLCETLHRHRTHFAHQHNTTCT
GNTPHHTNTVIGTADAQLLAAQSDAECWNTDTQYSEAHDSAKDSGYDEPTATTTTQQH
LTDATEVSGQHREIQARIIDLLNQIERSVNEGIPLNIRRKPRWETCLIEDGILQTTDSAREG
RTIRPARTRRLQLMVKLLATIYQLLATGTCCTKRELYYLHLELAQTPGYTYAALDDICA
LLDADPWELNVFNTSKGLIAGPIVLTVSGGTTIDCGTNRWGTAVPLDVGSVVAIQLSAR
LVLVVEKDTVFKRLVEDGIFDQFPNTVVLITAKGYPDVSTRLLLKKIADWTKVPIYALM
DADPHGIEIFCVYKFGSLAMVHQQQSLAVPSMRWIGLFPSDIELLGLQGVPLREHELKRI
EQMVKRPYTEGHIQRELLLLRQLATKAEIESLYNIASDFITTVYLKGKFNECLKHSSMKE
HPFLG
***Cardinal***
>g5128.t1
MVMVDERTPLTSDLSGPLPLVSGPSGNVHHLKSHESVRERQVRTFQCWICSAIMGAFAL
AIVISISYIIFGDATKPPLDGANATAADFPELLNLISFPLVDELPPQWNGTDVSEDAKAAAI
AEGEKALGDKELLEETLSSPPVNSPSFRHQKSVGATVAARLAAKVGFVEDRATKALVR
KLDIRHRGSVGRGPVMNLPRTHRHPQCDFNARYRSANGTCNNKERPYEYGVAMIPFRR
QLNPDYGDGISAPRASVDGTELPSARQVSLDIHRPSYHSDPNFSVMLAVWGQFLDHDIT
STALNQGVDGKPIECCDPGQPQHPECFPVPLGPGDPYYHQYNVTCMNFVRSVPAPTGHF
GPRQQLNQATAYIDGSVVYGSDEERMKKLRTGEGGRLRMLRTPDGRQLLPVSTDPLDG
CNEQEMNAAGKYCFESGDTRANENLHLTSMHLIWARHHNSLADGLARVNPHWDDERL
FQEARRILAAQMQHITYAEFVPVIVGNATAARMDLLPESTGRDDTYNASVDASIANVFA
GAAFRFAHTLLPGLMKKTRNPTSSSSGIELHRMLFNPYSLYAHDGLDNALGGAMSTSLA
KYDQYFSTELTEKLFEKADEHLLHNHPCGLDLVSLNIQRGRDHGLPAYPRWRKHCHLTP
ADSWAELERIVDPESFRQMKSIYRDPANVDVYSGALSEPPVKDGIVGPLLTCLLADQFLR
LKQGDSFWYERRRGPQRFTEGQLQQIYDTKLSSIICRNSDNIEQSPVHLMKRTDSRTNPE
TDCKQLDTFDFEPFREDKDAEPQHTRSAKIATDRVKVLVMEPKSAGTTTTRTTIEPEMV
ERDKATDSTTMITTTESLPTTLSNTVSGV
***ACE1***
>g6054.t1
MARTNTGETLTARYRLSPHCDGDFVREEPQQPGEPQHSPNGGYHSPPPVPMEIRGLLMG
RLRLGRRAIPLGLLCVTALLLLLPPSAIVQGRHHELNNGAALGSHQLSGAGGAGLSSQSA
QSGSLASGVISSAPAASSSSALSSGEEDLARITLSKDADPELGTLEREHVHSGATPRRRGL
TRRESNSDANDNDPLVVNTDKGRIRGITVEAPSGKKVNVWLGIPYAQPPVGPLRFRHPR
PAEKWTGVLNTTTPPNSCVQIVDTVFGDFPGATMWNPNTPLSEDCLYINVVAPRPRPKN
AAVMLWIFGGGFYSGTATLDVYDHRALASEENVIVVSLQYRVASLGFLFLGTPEAPGN
AGLFDQNLALRWVRDNIHRFGGDPSRVTLFGESAGAVSVSLHLLSALSRDLFQRAILQS
GSPTAPWALVSREEATLRALRLAEAVACPHEPSKLSEAVECLRGKDPHVLVNNEWGTL

GICEFPFVPVVDGAFLDETPQRSLASGRFKKTEILTGSNTEEGYYFIIYYLTELLRKEEGVT
VTREEFLQAVRELNPYVNGAARQAIVFEYTDWTEPDNPNSNRDALDKMVGDYHFTCN
VNEFAQRYAEEGNNVYMYLYTHRSKGNPWPRWTGVMHGDEINYVFGEPLNPSLGYTD
DEKDFSRKIMRYWSNFAKTGNPNPNTASSEFPEWPKHTAHGRHYLELGLNTSFVGRGP
RLRQCAFWKKYLPQLVAATSNLQVAAPPSAPCESSAFFYRPDLIVLIVSLLTVTPEQTPK
RPSDFEREDNQNDDDDEHSPEPAAARSQPPKKVNRTEQTATARGNPVLPAKEPQISEEE
RLNILKFVETEEPDGEVLDESGLKKMLLLFEKRVLKNQEMRIKFPDNAEKFMESEIELND
AIQELHAVATVPDLYPLLVELNGVASLLDLLSHQNSDISVAVVNLLQELTDVDILHESLD
GTETLIEALRNQQAAGLLVQNLERLDESVKEEADGVHNTLAIFENLIEVKSDIAKEVAEQ
GLLQWIMKRLRAKIPFDANKLYCSEILSILVQDTNENRITLGNIDGIDVLLQQLAAYKRH
DPNSAEEQEFMENLFNSLCSALMAKENREKFLKGEGLQLMNLMLREKKLSRNGSLKVL
DHAMAGPDGRDNCNKFVDILGLRTIFPLFMKTPKRSKKRLLSTDEHEEHIVSIIASMLRN
CKGSQRQRLLSKFTENDFEKVERLMELHRKYLDKVEAMDREIDQEMRVDDEDEQDDD
MVYVKRLSGGLFTLQLVDYVILEISCTDVVKQRVLKILNLHNGSMKMIRNVMREYAGN
LGDASDSDWREQEQAHILQLIDRF

***CYP6P9a/b***

>g6842.t1

MELINAVLAAFIFIVSTVYLFIRNKHNYWKDNGFPYAPNPHFLFGHAKGQTQTRHAADI
HQELYKKFKQLGERYVGMSQFIVPSVLVIDPELVKTILVKDFNVFHDHGVFNNAKDDPL
SAHLFALEGNPWRLLRQKLTPTFTSGRMKQMFGTIWDVALELDKYMEENYNQPEIEMK
DVLSRFTTDVIGTCAFGIECNSLRTPESDFRKYGNKAFELNPIIILKLFLSSSYPGLIRKLRL
KITYNDVEAFFMKIVRETVNYRESNNVKRNDFMNLLLQIKNKGKLDDNDDGTVGKGEV
GMTEAELAAQAFVFFLAGFETSSTTQSFCLYELAKNPDIQERLRQEINQAIDENDGQVTY
DVAMSIQYLDNVINETLRKYPPVESLSRVPSVDYLIPGTKHVIPKRTLVQIPVHAIQRDPD
HYPDPERFDPDRFTPEEVKKRHPFTFIPFGEGPRICIGLRFGVMQTKVGLITLLRKFRFSPS
ARTPDRVTFDPKMITLSPSAGNYLKVEKL

***CYP6P4***

>g6843.t1

MVLVEVILLSVVLLLSVAYLFLRERHSFWRKRGFPYKPNPSLLFGQMQGNGTTRHAAY
VTQEIYNYAKDRGERFVGYSFFFMPLVMVCDIELVKTILVKDFAVFHDRGMYSNARVD
PLSAHLFALEGHEWRALRQKLTPTFTSGRMKQMFGTMMQVAEELHRHLLANIGQELE
MKEILARFTTDVIGTCAFGIECNTFQHPDSDFLKYGKRVFEHKLLGVVKMTFAMLCKDT
ASKLGVKVTDPELEKFFLNLVHETVEYRERNDVQCNDFLDLLLQIKNKGCLVEQEEGHT
EQPDSSTGLTMNELAAQVFIFFVAGFETSSTVMNFCLYELAKNPDIQERLREELNRVIES
NGGELTYETVMGNEYLGQVVNETLRKYPPLETTLRVTAQDYTIPGTEHVIPRKVGVQVP
VFAIHRDPDHYPDPECFDPDRFTVEQCKNRPAYTFLPFGEGPRMCIGMRFGLMQIKLRGI
AEWSKMELLSYVLTAFVFVVSIAYLFLRSRHNYWRDRGIPYARAKPHLFMGHMEHFRT
KHGAIINEEIYRDLKSRGETIGGMSFFIIPGLVAVDPELIKTILVKDFNVFHDRGVYNDAK
ADPLSAHLFALEGHEWRVLRQKLTPTFTSGRMKQMFGTIQQVAEEFLKYMNENCHREI
EMKNVLARFTTDVIGTCAFGIECNTLKNPDSDFRKYGNKVFEQDALLMMKFVFAMMF
KSIAKGIGVKLTDEGVERFFLQVVRDTVEYRELNDVQRNDFMNLLLQIKNKGYLDERDL
VSADDAKGKAGLTLNELAAQVFVFFLAGFETSSTTMNFCLYELAKNPDIQERLRDEIER
AVEDHNGQVTYEMVMNNQYLDNVINETLRKYPPIESLTRVPMRDYTIPGTKYVIPKDTL
IQIPVYAIQRDPEFYPEPDQFNPDRFLPEEVKQRHPYVFLPFGEGPRICIGMRFGMMQAKL
GLITLLRNFRFSPSSQTPAEIVFDPKSFILSPTTVLRLQLIYWTSPTQCWFSFGSAVKHRKA
LTITMEPITLVLTGFIFIVSIVYLFVRSKHNFWKDQGVPYAPNPHFFYGHVKGQSRTRHG

ADINQELYKHFKQRGVPYGGISLFIMPSLIVVDPELVKTILVKDFNVFHDRGVFSNPKDD
PFTGNLFGLEGNPWRLLRQKLTPTFTSGRMKQMFGTIWEVALELEKYMEENYNQPEIE
MKDVLGRFTTDVIGTCAFGIECNTLKTPDSEFRKYGNKAFELDPVTLTKFFFASSYPHLA
RKLHVRTTQQDVEDFFMKIVRETVDYRESNNVQRNDFMNLLLQIKNKGKLDDQGTIVG
KGEVGLTHNELAAQVLIFFLAGFETSSTTLSFCLYELAKNPDIQDRLRDEITGAIDDNGGE
VTYDVAMNIQYLDNVINETLRKYPPVETLTRKPSQDYVIPGTKHVIPEGTIVQIPIYAIQR
DPDHFPDPERFDPDRFAPEEVKKRHPYVFVPFGEGPRICIGLRFGVMQTKVGLINLLLKF
RFSPSARTPDRVAFDPKMFTLSPIGGNYLKVEKVV

***RAD51***

>g6908.t1
MAQMEKSLQSASTVEDEEDYGPLLIGKLEGNGITNGDIKKLAEAGFHTVEAVAYAPKK
QLLAIKGISEAKADKILQEATKHVPMGFTTATEYHQKRSEIIQLTTGSKELDKLLGGGIET
GSITEIFGEFRTGKTQLCHTLAVTCQLPVSQNGGEGKCLYIDTEGTFRPERLLATAERYKL
VGADVLDNVAYARAYNTDHQMHLLMVASAMMAESRYALIIVDSATSLYRTDYSGRGE
LAARQTHLAKFLRMLLRLADEFGVAVLITNQVVAQVDGAAMFNPDPKKPIGGNIIAHAS
TTRLYMRKGRGEARICKIYDSPCLAEGEATFAINPDGIGDVKE

***RAB5***

>g12024.t1
MASSPRAGGAAQRPNGATQNKICQFKLVLLGESAVGKSSLVLRFVKGQFHEYQESTIGA
AFLTQTLCIDDTTVKFEIWDTAGQERYHSLAPMYYRGAQAAIVVYDIQNSDSFARAKT
WVKELQRQASPNIVIALAGNKADLANSRVVDYEEAKQYADDNGLLFMETSAKTAVNV
NDIFLAIAKKLPKNEGAGPQQNIRPTQNETNRQNSGCCAVSK

***CYP6M10***

>g12863.t1
MLSLFDFAFLVAALVAGLYYYLDRKRSYWKDRGVPGPKSELLLGNFGTVGTKEHITVP
MKKIYDEHKGKHPFAGIYQFVKPVALITDLELLKCVFVKDFQYFHDRGTFYNERDDPLS
AHLFNLEGQKWRTLRNKLSPTFTSGKMKMMFPTIVTAAKEFKDFMEETVKRENVFELK
DLLARFTTDVIGMCAFGIECNSMRNPDAEFRAMGRKIFEISPGTFKTMLMNGMPSLAKM
LRMKQTDQEVSDFFMNAVRDTINYRVTNKVKRNDFVDLLITMMSKDENSSDDESLTFN
EIAAQAFVFFLAGFETSSTLLTWTLYELALSEEIQEKGRQCVREVLKKHNGEMTYEAILD
MKYLDQILNESLRKYPPVPVHFRIASKDYQVPGTKSVLEAGTAVMVPVHAIHHDPAVFP
EPERFDPERFSPEEEAKRHPYAWTPFGEGPRICVGLRFGMMQARIGLAYLLDGFRFEPSP
KTTVPMELSTESFIMAPKGGLWLKVDKI

***KDR***

>g16696.t1
MTEDSDSISEEERSLFRPFTRESLQAIEARIADEEAKHRELERKRAEGEIRYDDEDEDEGP
QPDPTLEQGVPVPVRMQGSFPPELASTPLEDIDGFYSNQRVLRSAGPWHMLFFIVIIFLGS
FYLVNLILAIVAMSYDELQKKAEEEEAAEEEALREAEEAAAAKAAKLEAQQAAAAAAA
NPEIAKSPSDFSCHSYELFVGQEKGNDDNNKEKMSIRSEGLESVSEITRTTAPTATAAGT
AKARKVSADMTQDCTDDAGKIKHNDNPFIEPAQTQTVVDMKDVMVLNDIIEQAAGRH
SRTSDHGEDDDEDGPTFKDKALEFLMKMIDIFCVWDCCWVWLKFQEGVAFIVFDPFVE
LFITLCIVVNTLFMALDHHDMDPDMEKALKSGNYFFTATFAIEATMKLIAMSPKYYFQE
GWNIFDFIIVALSLLELGLEGVQGLSVLRSFRLLRVFKLAKSWPTLNLLISIMGRTMGALG
NLTFVLCIIIFIFAVMGMQLFGKNYVDNVDRFPDHDLPRWNFTDFMHSFMIVFRVLCGE
WIESMWDCMLVGDVSCIPFFLATVVIGNLVVLNLFLALLLSNFGSSSLSAPTADNETNKI
AEAFNRISRFSNWIKMNVANALKFVKNKLTSQIASVQPTEHGENELELTPDDILADGLLK

KGIKEHNQLEVAIGDGMEFTIHGDLKNKAKKNKQIMNNSKDDDTASIKSYGSHKNRPFK
DESHKGSAETMEGEEKRDASKEDLGIDEELDDEGEGEEGPLDGELIIHAEEDEVIEDSPA
DCCPDNCYKKFPVLAGDDDAPFWQGWGNLRLKTFQLIENKYFETAVITMILLSSLALLS
LINLAAIWVGAADIPAFRSMRTLRALRPLRAVSRWEGMRCVDKNKTTLPHEIIPDVNAC
KAENYTWENSPMNFDHVGKAYLCLFQVATFKGWIQIMNDAIDSRDVGKQPIRETNIYM
YLYFVFFIIFGSFFTLNLFIGVIIDNFNEQKKKAGGSLEMFMTEDQKKYYNAMKKMGSK
KPLKAIPRPRWRPQAIVFEIVTNKKFDMIIMLFIGFNMLTMTLDHYKQSETFSAVLDYLN
MIFICIFSSECLMKIFALRYHYFIEPWNLFDFVVVILSILGLVLSDIIEKYFVSPTLLRVVRV
AKVGRVLRLVKGAKGIRTLLFALAMSLPALFNICLLLFLVMFIFAIFGMSFFMHVKDKSG
LDDVYNFKTFGQSMILLFQMSTSAGWDGVLDGIINEEDCLPPDNDKGYPGNCGSSTIGIT
YLLAYLVISFLIVINMYIAVILENYSQATEDVQEGLTDDDYDMYYEIWQQFDPDGTQYV
RYDQLSDFLDVLEPPLQIHKPNRYKIISMDIPICRGDMMFCVDILDALTKDFFARKGNPIE
ETAELGEVQQRPDEVGYEPVSSTLWRQREEYCARLIQHAWKRYKQRHGGGTDGSGDD
LEIDACDNGNDGGDGNDNDGSGGAAVGSGDNGSQIAGGSIGGGGGGGTPGSGKSKGIL
GGSQANVGVMEHSLSSKESPDNNGDPQGRQTAVLVESDGFVTKNGHRVVIHSRSPSITS
RTADV

***KH***

>g18313.t1
MAPASDKYKRTNTNGMQHQPLDVAIVGGGLVGSLLALHLGKKGHEVNLYEYREDIRT
AELVIGRSINLALSARGRRALAEVGLEEALLDHGIPMSGRMLHDVNGNRKIVPYDGNTN
QCIYSVGRKHLNEVLLNAAEKYPNIHLHFNHKLVSANLDEGNLSMVDPLTKEVKSARA
DLIVGCDGAYSAVRKEIVKRPRYDFSQTYIEHGYLELCIPPTASGEFAMPHNYLHIWPRG
QFMMIALPNQDRTWTVTLFMPFTQFHSITDPGRLIDFFRQYFPDAIELIGRERLIKDFFKT
KPQPLVMIKCRPYHVGSKALIIGDAAHAMVPFYGQGMNAGFEDCSVLTDLFNQYGTDL
TRILPEFSEKRWEDAHAICDLAMYNYIEMRDLVTKRSYLLRKKLDELLFWLLPNTWVPL
YNSVSFSHMRYSKCIANRAWQDKILTRVLYGASFVSVAAIGGLAYRHMTVGHLERLSS
AILSTFQLLKPNTASV

***FREP1***

>g18569.t1 (gray: NO homology, green full homology, yellow highlight from Nui et al.[13])
MVPKAPKMVYSFVPAVVIILGLVSFSHSIALNGDVPGNINKAAVSVNSPQPALEGVLTA
VLPDEPNADSERFDALTLEDQVRLLSKQLNALTYQRREDYKMLENSLKKYVRKNAAEI
TDGQIREELDQLSEGSPEASKPGLAELTDGCYGTDRLSQSHQVCPRLLLLLLAVREQKQ
DKKKATLKTTEDVNQLREASSPSKERLTVQWLSQSISEIRSELAELQSSFGGGSKDAQFR
NQLLEDLSTLRSEFGTAKLELESLRSRQEKTEVLVRELQEEAVQSADDIRRSLNMRHEKH
PTSDCERYYGRWYPRVRSLGMVMIMIMADRSESIFSHGEDAAGIMCEHCVKIMTSKFSA
RFVCAPGIGEPLGNGQGMGPIPLLATVECQFTQQKSDRFVPLMGRGELDGTTAQQDKSI
LPHTIDFVEPEADHRMRHQRFIRQQLHELEVKQSVMKRQLSELHGHRLADRLRSVEIEQ
RRLASASFNVSRQIAGLDKLHGSMLELLEDVEAIQGKFEKTVPDMRREIAKVEFGVAQA
ASEQGLVREEVHNAAKSIQAMAVSVSALQEERDTVKRLQGEVHELKDELARIRSASVL
HREMAHNRLEKLEAGSKSDYNGSTTAPHRTLTELERTTKLVQQLESVENEYESIINKLPR
DCSQIERMRTTGTANQPGDGGLYLIAPAEQHHPLMTQCFGEWTTVQRRQDGTVDFNRS
WEEYAQGFGTPAGEFWIGNQALHHLTQDNCSRLRIVMQDIYDNTWFADYATFRIDSRD
AGFRLDLGGYSGNASDAFEYQNHMQFSAIDVDRDISNTHCAGNYEGGWWFSHCQHAN
LNGRYNLGLTWFDASRNEWIAVKSSHMMIARRPECDANVTELPVEQQQQPVTSGLAAA
TSTPSDASRSLHGTNPRQQHDSTTTAPTFS

***MRE11***

>g24450.t1
MSESASQTADINPDDTIKILVASDIHLGFNEKDPIRGDDSFIAFEEVLQHALENEVDALLL
GGDLFHVANPSTNTLDRCFRLLKTYTLGDKPIRLEFLSDQNDNFLESLSHTVNYEDPNM
NIAIPVFSIHGNHDDSGGAGKVSSMNLLSTNGYVNYFGKWTDLSKIDIRPILLRKGETKL
ALYGLSYMSDARLCRLLDDAKVFIEKPDEPGFFSIMVLHQNRAERGPKNYLPESSLPQFL
DLIIWGHEHDCRIEPEENSAKKFYVSQPGSTVATSLSEGEAIQKCCGLLSIHKGLFRMDPI
PLKSVRPFVFESVDLATVQEELALDEGDVQQRVQDFATERIEAMIERAKTKITGYARQP
KLPLIRLRLGLTEIEQQFNAIRFGFRYHGRVANPQDMVIFKKKPKVKVKDELGNALDKA
ALQEAYRNQREQRAQRAEEIVDRYFREADVVNQLEVLNPRSMAELCRRMVDYEDDDA
PEKIIKFYEDKALSFLRSQDNTSEEGICEALAGFHTADPDIHEKVLSMLDARSNRQDATD
PLQRFRDLDDDGMAGGNDGSSVRGDPPNTTVAAGKPAARGARGGRGSRGGGTARGA
ASKAASTAGSTRGRSQTSITSMFSQQSTNNSIASVATRTSSRKTATKKARQMDFDSDDE

**Text S2:** Protein Sequences for the Genes of Interest listed in Table 3.

**Supplementary Table S7:** Level of homology between orthologous ORs from *An. stephensi* and *An. funestus* compared to *An. gambiae.*

| Gambiae ID | Steph ID | Steph Start | Steph End | e-value | Funestus ID | Funes Start | Funes End | e-value2 |
|---|---|---|---|---|---|---|---|---|
| Agamor1 | g14613.t1 | 1 | 379 | 0 | g6763.t1 | 1 | 404 | 0 |
| Agamor2 | g13672.t1 | 1 | 375 | 0 | g8764.t1 | 1 | 347 | 0 |
| Agamor3 | g3881.t1 | 1 | 389 | 0 | g18528.t1 | 692 | 1081 | 0 |
| Agamor5 | g3879.t1 | 1 | 389 | 0 | g18528.t1 | 1 | 375 | 0 |
| Agamor7 | g3284.t1 | 20 | 481 | 0 | g1526.t1 | 1 | 513 | 0 |
| Agamor8 | g6261.t1 | 69 | 475 | 0 | g4592.t1 | 1 | 399 | 0 |
| Agamor9 | g13119.t1 | 1 | 408 | 0 | g6426.t1 | 1 | 133 | 3.99E-57 |
| Agamor11 | g11090.t1 | 1 | 425 | 0 | g19674.t1 | 1 | 424 | 0 |
| Agamor16 | g13438.t1 | 454 | 823 | 0 | g18528.t1 | 377 | 685 | 2.04E-38 |
| Agamor18 | g13457.t1 | 2 | 384 | 0 | g8951.t1 | 3 | 343 | 5.13E-153 |
| Agamor22 | g12730.t1 | 6 | 363 | 0 | g13510.t1 | 1 | 357 | 0 |
| Agamor23 | g12231.t1 | 92 | 385 | 0 | g8459.t1 | 61 | 330 | 2.45E-158 |
| Agamor25 | g4286.t1 | 9 | 371 | 0 | g3252.t1 | 169 | 389 | 5.82E-15 |
| Agamor26 | g9046.t1 | 1 | 400 | 0 | g3252.t1 | 214 | 388 | 3.75E-14 |
| Agamor27 | g9046.t1 | 1 | 400 | 0 | g8764.t1 | 160 | 344 | 3.71E-11 |
| Agamor28 | g2853.t1 | 21 | 413 | 0 | g3252.t1 | 1 | 394 | 0 |
| Agamor29 | g16275.t1 | 1 | 377 | 0 | g7223.t1 | 1 | 362 | 0 |
| Agamor31 | g20901.t1 | 6 | 338 | 0 | g8764.t1 | 160 | 344 | 3.71E-11 |
| Agamor32 | g20871.t1 | 1 | 382 | 0 | g11464.t1 | 15 | 393 | 0 |
| Agamor36 | g180.t1 | 1 | 404 | 0 | g19685.t1 | 475 | 852 | 0 |
| Agamor38 | g3028.t1 | 1 | 409 | 0 | g3082.t1 | 1 | 405 | 0 |
| Agamor39 | g3028.t1 | 1 | 409 | 0 | g3082.t1 | 1 | 405 | 0 |
| Agamor40 | g3288.t1 | 1 | 438 | 0 | g1524.t1 | 3 | 439 | 0 |
| Agamor42 | g7200.t1 | 1 | 378 | 0 | g15.t1 | 232 | 615 | 0 |
| Agamor48 | g18340.t1 | 396 | 792 | 0 | g18528.t1 | 701 | 1081 | 2.51E-43 |
| Agamor49 | g18340.t1 | 1 | 388 | 0 | g18528.t1 | 704 | 1081 | 7.76E-60 |

| | | | | | | | | |
|---|---|---|---|---|---|---|---|---|
| Agamor52 | g1315.t1 | 35 | 417 | 0 | g19444.t1 | 35 | 418 | 0 |
| Agamor54 | g11749.t1 | 1 | 404 | 0 | g16016.t1 | 1 | 403 | 0 |
| Agamor56 | g9046.t1 | 1 | 400 | 0 | g3252.t1 | 99 | 388 | 2.53E-17 |
| Agamor57 | g9046.t1 | 1 | 398 | 0 | g3252.t1 | 32 | 391 | 1.22E-17 |
| Agamor58 | g8440.t1 | 180 | 582 | 0 | g1656.t1 | 1 | 430 | 0 |
| Agamor59 | g6376.t1 | 1 | 327 | 0 | g4366.t1 | 97 | 365 | 6.16E-144 |
| Agamor61 | g10285.t1 | 1 | 420 | 0 | g17618.t1 | 398 | 751 | 0 |
| Agamor62 | g10269.t1 | 16 | 356 | 0 | g17598.t1 | 3 | 340 | 0 |
| Agamor63 | g10281.t1 | 8 | 384 | 0 | g17618.t1 | 8 | 403 | 0 |
| Agamor65 | g13119.t1 | 26 | 403 | 0 | g6426.t1 | 18 | 147 | 2.84E-48 |
| Agamor66 | g7370.t1 | 1 | 393 | 0 | g2456.t1 | 1 | 393 | 0 |
| Agamor67 | g7370.t1 | 1 | 393 | 0 | g2456.t1 | 1 | 393 | 0 |
| Agamor68 | g15209.t1 | 1 | 377 | 0 | g7544.t1 | 1 | 258 | 4.12E-136 |
| Agamor75 | g3566.t1 | 2 | 409 | 0 | g655.t1 | 3 | 409 | 0 |
| Agamor76 | g3566.t1 | 2 | 409 | 0 | g655.t1 | 3 | 409 | 0 |
| Agamor77 | g3567.t1 | 1 | 381 | 0 | g657.t1 | 1 | 381 | 0 |
| Agamor78 | g3566.t1 | 2 | 409 | 0 | g655.t1 | 3 | 409 | 0 |
| Agamor79 | g3567.t1 | 1 | 381 | 0 | g657.t1 | 1 | 381 | 0 |
| Agamor53 | g13436.t1 | 1 | 384 | 9.80E-179 | g7223.t1 | 1 | 362 | 1.26E-138 |
| Agamor41 | g1302.t1 | 1 | 353 | 4.29E-176 | g19440.t1 | 370 | 739 | 9.47E-102 |
| Agamor4 | g3880.t1 | 1 | 381 | 1.23E-175 | g18528.t1 | 374 | 687 | 7.89E-136 |
| Agamor14 | g13457.t1 | 381 | 796 | 1.31E-171 | g18528.t1 | 1 | 376 | 8.45E-73 |
| Agamor60 | g10271.t1 | 5 | 416 | 1.84E-171 | g17599.t1 | 5 | 428 | 0 |
| Agamor30 | g13437.t1 | 1 | 384 | 1.61E-169 | g7223.t1 | 1 | 362 | 4.01E-126 |
| Agamor6 | g19986.t1 | 1 | 332 | 1.16E-166 | g11731.t1 | 6 | 189 | 6.20E-103 |
| Agamor51 | g13457.t1 | 381 | 795 | 1.72E-164 | g18528.t1 | 1 | 376 | 1.18E-72 |
| Agamor34 | g6340.t1 | 1 | 240 | 7.60E-156 | g4400.t1 | 1 | 380 | 0 |
| Agamor10 | g13672.t1 | 1 | 375 | 7.61E-154 | g8762.t1 | 1 | 346 | 0 |

| | | | | | | | | |
|---|---|---|---|---|---|---|---|---|
| Agamor15 | g13438.t1 | 861 | 1238 | 6.16E-150 | g18528.t1 | 700 | 1080 | 1.43E-76 |
| Agamor45 | g4285.t1 | 5 | 370 | 4.14E-149 | g2456.t1 | 20 | 386 | 5.09E-15 |
| Agamor64 | g10284.t1 | 1 | 416 | 1.42E-141 | g17618.t1 | 54 | 405 | 3.50E-29 |
| Agamor13 | g13438.t1 | 861 | 1239 | 3.36E-139 | g18528.t1 | 700 | 1081 | 2.73E-79 |
| Agamor17 | g13438.t1 | 861 | 1238 | 5.83E-139 | g18528.t1 | 700 | 1080 | 1.90E-71 |
| Agamor70 | g15210.t1 | 2 | 319 | 7.52E-138 | g2456.t1 | 1 | 389 | 1.37E-39 |
| Agamor72 | g15210.t1 | 2 | 319 | 9.32E-138 | g2456.t1 | 1 | 389 | 3.39E-44 |
| Agamor37 | g6342.t1 | 1 | 252 | 1.49E-137 | g4399.t1 | 6 | 372 | 0 |
| Agamor73 | g15210.t1 | 2 | 319 | 2.88E-135 | g2456.t1 | 1 | 389 | 2.93E-44 |
| Agamor46 | g13438.t1 | 106 | 442 | 1.42E-132 | g8961.t1 | 1 | 302 | 6.82E-124 |
| Agamor69 | g15210.t1 | 2 | 319 | 2.36E-132 | g2456.t1 | 57 | 389 | 1.46E-35 |
| Agamor47 | g13438.t1 | 106 | 443 | 1.92E-130 | g8961.t1 | 1 | 302 | 6.67E-124 |
| Agamor71 | g15212.t1 | 2 | 320 | 2.76E-127 | g2456.t1 | 1 | 389 | 2.51E-41 |
| Agamor74 | g15210.t1 | 2 | 319 | 5.99E-125 | g2456.t1 | 1 | 389 | 4.04E-41 |
| Agamor15 | g13438.t1 | 454 | 822 | 2.11E-117 | g18528.t1 | 11 | 372 | 8.20E-67 |
| Agamor13 | g13438.t1 | 454 | 822 | 6.66E-116 | g18528.t1 | 377 | 685 | 8.66E-33 |
| Agamor16 | g13438.t1 | 861 | 1239 | 4.44E-110 | g18528.t1 | 710 | 1081 | 5.88E-79 |
| Agamor17 | g13438.t1 | 454 | 822 | 1.15E-107 | g18528.t1 | 11 | 372 | 1.10E-58 |
| Agamor35 | g20899.t1 | 1 | 202 | 1.48E-105 | g11464.t1 | 15 | 392 | 2.95E-97 |
| Agamgr55 | g13438.t1 | 453 | 823 | 1.35E-97 | g18528.t1 | 11 | 374 | 1.54E-63 |
| Agamor18 | g13457.t1 | 385 | 796 | 3.88E-86 | g8951.t1 | 3 | 343 | 5.13E-153 |
| Agamor48 | g18340.t1 | 17 | 388 | 6.27E-77 | g18528.t1 | 8 | 374 | 1.27E-35 |
| Agamgr55 | g13438.t1 | 861 | 1237 | 1.45E-76 | g18528.t1 | 710 | 1079 | 5.75E-68 |
| Agamor20 | g3879.t1 | 8 | 387 | 2.57E-69 | g18528.t1 | 700 | 1079 | 8.14E-63 |
| Agamor19 | g3879.t1 | 8 | 387 | 3.66E-69 | g18528.t1 | 700 | 1079 | 1.38E-60 |
| Agamor12 | g3881.t1 | 7 | 387 | 5.40E-66 | g18528.t1 | 700 | 1079 | 4.28E-61 |
| Agamor21 | g3881.t1 | 7 | 387 | 2.73E-65 | g18528.t1 | 700 | 1079 | 2.63E-60 |
| Agamor50 | g3881.t1 | 7 | 387 | 2.19E-63 | g18528.t1 | 700 | 1079 | 6.37E-59 |

| | | | | | | | | |
|---|---|---|---|---|---|---|---|---|
| Agamor49 | g18340.t1 | 402 | 792 | 6.48E-63 | g18528.t1 | 9 | 374 | 3.65E-51 |
| Agamor51 | g13457.t1 | 7 | 382 | 3.70E-62 | g18528.t1 | 694 | 1079 | 5.07E-70 |
| Agamor14 | g13457.t1 | 7 | 382 | 9.20E-59 | g18528.t1 | 694 | 1081 | 1.92E-70 |
| Agamor33 | g18446.t1 | 294 | 400 | 1.92E-51 | g8762.t1 | 249 | 342 | 1.79E-13 |
| Agamor68 | g15209.t1 | 459 | 750 | 2.00E-41 | g7544.t1 | 1 | 258 | 4.12E-136 |
| Agamor16 | g13438.t1 | 7 | 443 | 1.76E-34 | g18528.t1 | 17 | 374 | 1.64E-71 |
| Agamor46 | g13438.t1 | 453 | 825 | 6.05E-34 | g8961.t1 | 1 | 302 | 6.82E-124 |
| Agamor46 | g13438.t1 | 1 | 101 | 2.04E-31 | g8961.t1 | 1 | 302 | 6.67E-124 |
| Agamor47 | g13438.t1 | 453 | 825 | 3.14E-31 | g8961.t1 | 1 | 302 | 6.67E-124 |
| Agamor47 | g13438.t1 | 1 | 101 | 1.00E-30 | g8961.t1 | 1 | 302 | 6.67E-124 |
| Agamor47 | g13438.t1 | 863 | 1239 | 1.21E-30 | g8961.t1 | 1 | 302 | 6.67E-124 |
| Agamor15 | g13438.t1 | 109 | 443 | 7.89E-30 | g18528.t1 | 377 | 686 | 1.25E-34 |
| Agamor46 | g13438.t1 | 863 | 1239 | 1.50E-29 | g8961.t1 | 1 | 302 | 6.82E-124 |
| Agamor43 | g1301.t1 | 75 | 373 | 3.07E-27 | g15671.t1 | 250 | 553 | 5.33E-170 |
| Agamor13 | g13438.t1 | 190 | 443 | 3.51E-26 | g18528.t1 | 700 | 1081 | 2.73E-79 |
| Agamgr55 | g13438.t1 | 108 | 443 | 4.63E-26 | g18528.t1 | 356 | 685 | 2.23E-35 |
| Agamor33 | g18446.t1 | 113 | 213 | 8.34E-26 | g8762.t1 | 249 | 342 | 1.79E-13 |
| Agamor44 | g1301.t1 | 83 | 409 | 1.89E-25 | g15671.t1 | 250 | 548 | 2.97E-170 |
| Agamor24 | g12231.t1 | 41 | 369 | 3.88E-24 | g15680.t1 | 1 | 181 | 2.60E-107 |
| Agamor17 | g13438.t1 | 220 | 443 | 1.41E-23 | g18528.t1 | 377 | 685 | 7.53E-34 |
| Agamor68 | g15209.t1 | 361 | 460 | 4.18E-14 | g7544.t1 | 1 | 258 | 4.12E-136 |
| Agamgr55 | g13438.t1 | 1 | 101 | 0.006 | g18528.t1 | 356 | 685 | 2.23E-35 |
| Agamor13 | g13438.t1 | 8 | 106 | 0.52 | g18528.t1 | 377 | 685 | 8.66E-33 |
| Agamor17 | g13438.t1 | 8 | 106 | 1.9 | g18528.t1 | 377 | 685 | 7.53E-34 |
| Agamor15 | g13438.t1 | 8 | 102 | 5.3 | g18528.t1 | 377 | 686 | 1.25E-34 |
